# Structure-function-dynamics relationships in the peculiar *Planktothrix* PCC7805 OCP1: impact of his-tagging and carotenoid type

**DOI:** 10.1101/2022.01.04.474796

**Authors:** Adjélé Wilson, Elena A. Andreeva, Stanislaw J. Nizinski, Léa Talbot, Elisabeth Hartmann, Ilme Schlichting, Gotard Burdzinski, Michel Sliwa, Diana Kirilovsky, Jacques-Philippe Colletier

**Affiliations:** Université Paris-Saclay, CEA, CNRS, Institute for Integrative Biology of the Cell (I2BC), 91198 Gif-sur-Yvette, France; Univ. Grenoble Alpes, CEA, CNRS, Institut de Biologie Structurale, 38000 Grenoble, France; Max-Planck-Institut für medizinische Forschung, Jahnstrasse 29, 69120 Heidelberg, Germany; Univ. Lille, CNRS, UMR 8516, LASIRE, LAboratoire de Spectroscopie pour les Interactions, la Réactivité et l’Environnement, Lille 59000, France; Quantum Electronics Laboratory, Faculty of Physics, Adam Mickiewicz University in Poznań, Uniwersytetu Poznańskiego 2, Poznan 61-614, Poland

**Keywords:** cyanobacteria, flash photolysis, photosynthetic pigments, structure function relationships, X-ray diffraction

## Abstract

The orange carotenoid protein (OCP) is a photoactive protein involved in cyanobacterial photoprotection. Here, we report on the functional, spectral and structural characteristics of the peculiar *Planktothrix* PCC7805 OCP (Plankto-OCP). We show that this OCP variant is characterized by higher photoactivation and recovery rates, and a stronger energy-quenching activity, compared to other OCP studied thus far. We characterize the effect of the functionalizing carotenoid and of his-tagging on these reactions, and identify the time scales on which these modifications affect photoactivation. The presence of a his-tag at the C-terminus has a large influence on photoactivation, thermal recovery and PBS-fluorescence quenching, and likewise for the nature of the carotenoid that additionally affects the yield and characteristics of excited states and the ns-s dynamics of photoactivated OCP. By solving the structures of Plankto-OCP in the ECN- and CAN-functionalized states, each in two closely-related crystal forms, we further unveil the molecular breathing motions that animate Plankto-OCP at the monomer and dimer levels. We finally discuss the structural changes that could explain the peculiar properties of Plankto-OCP.

**Highlights:**

- Complete functional characterization of *Synechocystis* and *Planktothrix* OCP
- Hitherto unknown structures of ECN- and CAN-functionalized *Planktothrix* OCP
- Insights into fs-s timescale photodynamics of ECN- and CAN-functionalized *Synechocystis* and *Planktothrix* OCP

## Introduction

Photosynthetic organisms have evolved to make use of nearly all photons absorbed by their light-harvesting antennas. Under light stress conditions, however, the photosynthetic electron transport chain becomes saturated leading to the formation of harmful reactive oxygen species (ROS), e.g., singlet oxygen (^1^O_2_) [1,2], that can damage the photosystems and other cellular machineries, eventually leading to cell death. Accordingly, photosynthetic organisms have developed a variety of mechanisms, altogether referred-to as non-photochemical quenching (NPQ), that are aimed at reducing the amount of energy reaching the photochemical reactions centers thereby avoiding accumulation of ROS [3]. In a vast majority of cyanobacterial strains, the main light-harvesting antenna is a large soluble complex, the phycobilisome (PBS), and the soluble 35 kDa photoactive Orange Carotenoid Protein (OCP) is at the center of the NPQ mechanism (for review: [4–6]). OCP is capable both of dissipating the excess energy harvested by the PBS [7], and of quenching the produced harmful singlet-oxygen [8,9]. For the energy quenching mechanism to be elicited, OCP must be photoactivated, which triggers the changes in protein structure and pigment position required for PBS binding and discharge of its excessive energy [10,11]. Specifically, upon absorption of a blue-green photon, OCP converts from an inactive dark-adapted state (denoted as OCP^O^, due to its orange color) into an active light-adapted state (denoted OCP^R^, due to its red color). OCP^O^ is characterized by two absorption maxima at 475 and 495 nm (vibronic structure), while OCP^R^ displays a single broader absorption peak between 510 and 530 nm [11]. The photoactivation quantum yield of the protein is notoriously low, viz. 0.2 % [12], meaning that the OCP-supported photoprotective mechanism is at play only under high light conditions and that the concentration of OCP^R^ is null, or very low, in darkness and under low light conditions [11,13]. Phylogenic studies of OCP sequences allowed their classification into three distinct clades, viz. OCP1, OCP2 and OCPX [14]. Members of the OCP1 clade are characterized by a slow OCP^R^ to OCP^O^ thermal recovery (at 8°C) that is accelerated by the presence of the fluorescence recovery protein (FRP), whereas OCP2 and OCPX exhibit a faster thermal recovery (even at 8°C) that is not affected by the presence of FRP [15,16].

The best characterized OCP are OCP1 from *Arthrospira maxima* and *Synechocystis* PCC 6803, hereafter referred to as *Arthrospira* and *Synechocystis* OCP, respectively. For both, the dark-adapted structure was solved by X-ray crystallography [8,17], revealing a conserved two-domain modular architecture. The fully α-helical N-terminal domain (NTD, residues 1-165), unique to cyanobacteria, and the C-terminal domain (CTD, residues 187-320), structurally belonging to the nuclear transport factor-2 superfamily (NTF2), encase at their interface a ketocarotenoid pigment, e.g., 3’-hydroxyechineone (3’-hECN) [8] or ECN [17]. Notwithstanding the presence of a linker that covalently attaches the NTD and CTD, the dark-adapted state is stabilized by two main protein interfaces, viz. (i) the central interdomain interface, which features two highly-conserved H-bond (N104-W277) and salt-bridge (R155-E244); and (ii) the interface between the N-terminal helix (also coined, N-terminal extension or NTE) and the CTD β-sheet [8], which features six to seven H-bonds depending on species. Additionally, the ketocarotenoid pigment buries ≈ 95% of its highly hydrophobic surface into the binding tunnel spanning the two domains (buried surface area (BSA) of 786 Å^2^; a surface complementarity of ∼83%), thereby contributing to the stabilization of the OCP^O^. The sole polar interactions between the ketocarotenoid and the protein scaffold are the H-bonds established between the carbonyl oxygen of its β1 ring and the side chain hydroxyl and amine of CTD residues Y201 and W288 (*Synechocystis* OCP residue numbering), respectively [8,17]. Rupture of these H-bonds is the first event along a photo-activation cascade that involves several ‘red’ intermediate states spanning the ps to second time scale [12,18–20] and culminates with dissociation of the two domains following the 12 Å translocation of the carotenoid into the NTD [21]. Dissociation of the two domains is essential for the energy-quenching function, as only OCP^R^ is capable of binding to the PBS [10,11]. This activity is measured as the quenching of PBS fluorescence, itself induced by exposure to blue-light.

Despite the considerable knowledge acquired on OCP in the last two decades, the photoactivation mechanism is still under debate. Notably, the very first instants of the photoactivation mechanism remains elusive. Numerous computational and time-resolved spectroscopic studies have recently sought to shed light on the structure, formation and decay of the carotenoid excited states associated with OCP photoactivation ([19,22–24]; Figure 1). Notably, it was demonstrated that upon photoexcitation, three ps-lived intermediate states are formed following the sub-ps decay of the initial S_2_ state, viz. an S_1_ and an intra-molecular charge transfer (ICT) excited states [25,26], and an S* state [19] that was initially proposed to correspond to a vibrationally hot ground-state (S_0_) population [27]. Recently, however, it was proposed that the S* state, characterized by a longer lifetime than the S_1_ and ICT states [19,28], is also an excited state which serves as the precursor of the first photoproduct, P_1_, in which the H-bonds between the carotenoid and the protein are broken and the protein is (therefore) already ‘red’ (difference absorption spectrum peaking at 565 nm) [19]. The debate however remains open concerning the nature of the S* state [29–31] and its putative role as the precursor of P_1_. Indeed, recent results from our laboratories show that while the photoactivation speed of OCP^R^ is independent of irradiation light (470 nm versus 540 nm), the concentration of S* decreases by ≈ 30% when 540 nm light is used to trigger photoactivation [32]. Hence, S* cannot be the sole precursor of P_1_. Evolution of P_1_ (50 ns lifetime), wherein the carotenoid is likely untethered from its H-bonding partners in the CTD, leads to a repositioning of the carotenoid in the tunnel, in close vicinity of the dark-state position (P_2_; 0.5 – 1 µs) [19]. After a first partial movement into the NTD (P2’; 10 µs) [19], the ketocarotenoid completes its translocation in around 10 µs, reaching the position that it occupies in the final OCP^R^ (P_3_) [19,23]. Conformational changes in the NTE and CTT ensue (P_M_ and P_X_ ∼10 ms and 35 ms), followed by an opening of the protein upon dissociation of the two domains (∼100 ms) [23]. Thus, the formation of the photoactive OCP^R^ is a multi-step reaction spanning at least twelve decades in times [12,19,23]. All steps, including the P_2’_ to P_3_ and P_3_ to P_M_ transitions, are accompanied by recovery to the initial OCP^O^ state, explaining the low quantum yield [12,19,23]. About 1 and 0.2 % of molecules reach the P_1_ state [19] and the final OCP^R^ state [12,19,23,32] (Figure 1), respectively.

**Figure 1:**
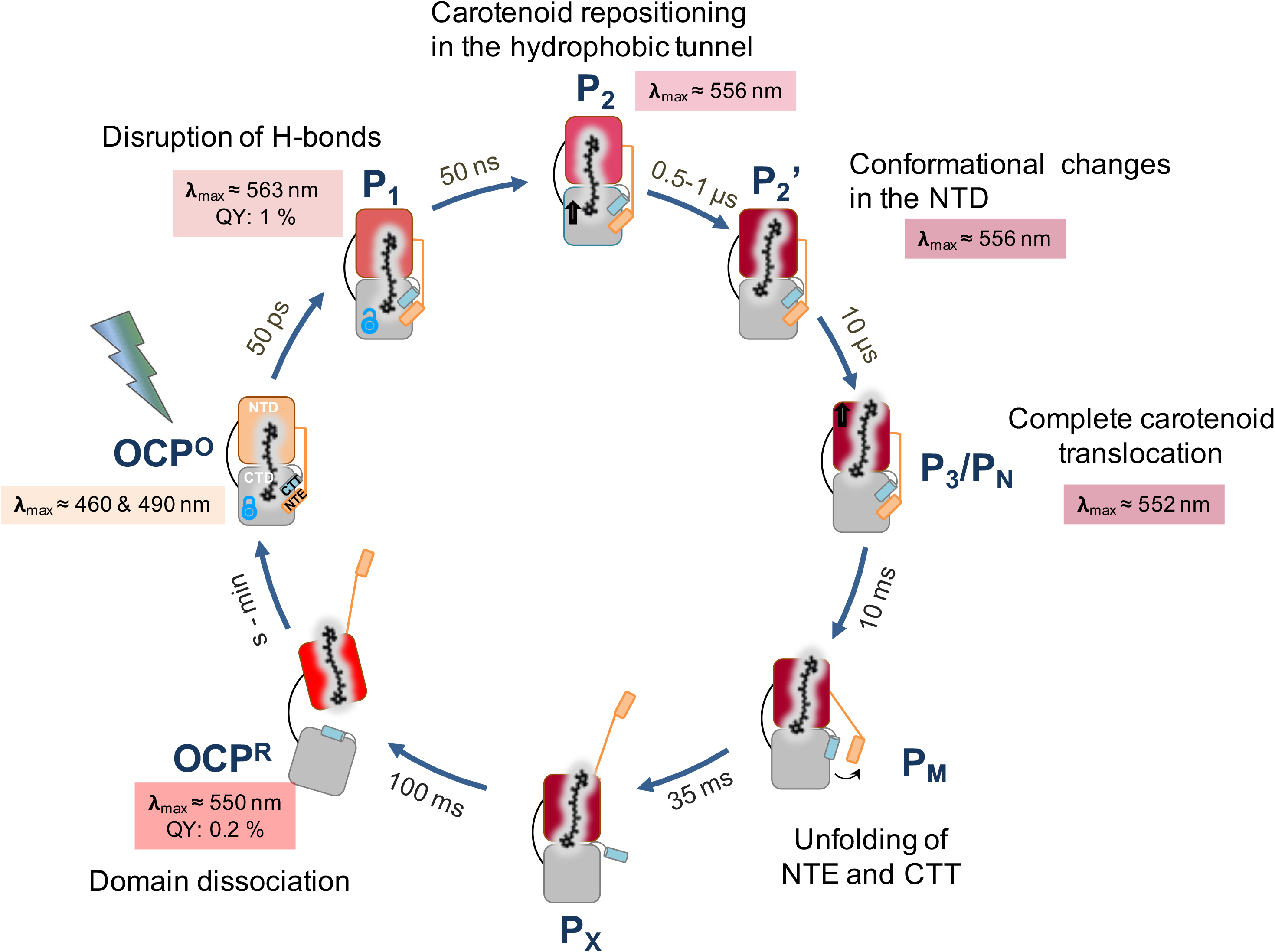
Formation of OCP^R^ is a multi-step reaction that spans twelve decades in time. The presented model for OCP photoactivation summarizes findings from multiple studies [12,19,23,32]. The wavelength of maximum difference-absorbance (after subtraction of the OCP^O^ signal) is indicated for all intermediate states characterized to date. The P_1_, P_2_, P_2’_ and P_3_ were observed by Konold et al. 2019 in transient absorption UV-Vis experiments upon excitation at 470 nm of Ntag-Syn-OCP_ECN_ [19]. Of important note, P_2’_ was only observed in transient IR experiments, suggesting that it is characterized by a conformational change in the protein scaffold that does not influence the carotenoid electronic properties. P_N_/P_M_ and P_X_ were observed by Maksimov et al. in time-resolved fluorimetry experiments conducted on the ECN-functionalized OCP-3FH mutant (W41F, W101F, W110F and W277H) featuring a single tryptophan at position W288, following excitation at 262 nm [23].

In all recent studies on OCP, the protein was expressed recombinantly [33] and a six-histidine tag (his-tag) was introduced at the N-terminus (OCP produced in *E. coli* cells) or C-terminus (OCP produced in *Synechocystis* and *E. coli* cells) to accelerate purification. It yet remains unclear whether or not presence of a his-tag at either the N- or C-terminal extremity of the protein influences the kinetics of photoactivation, PBS quenching and thermal recovery, respectively. Indeed, after migration of the carotenoid from the NTD/CTD interface into the NTD (≈ 0.5 µs), formation of OCP^R^ will requires, prior to domain separation, the detachment from the CTD β-sheet of the N-terminal helix αA (or NTE) [19,34,35] and the C-terminal helix αN (or CTT) [35–37], respectively. Supporting the hypothesis that his-tagging could affect OCP function is our recent report that the his-tag position affects i) the photoactivation efficiency, with OCP^R^ accumulation being slower when the his-tag is present at the C-terminus (C-tagged OCP); ii) the lifetime of excited states, with those being shorter-lived for C-tagged OCP); and iii) the spectral signatures of the S_1_ and S* states. It was also found that the position of the tag does not influence the P_1_ formation quantum yield (QY), suggesting that the more efficient photoactivation of the N-tagged protein is related to molecular events occurring on the nano- to millisecond timescales [32].

The influence of the carotenoid type on OCP the various facets of function is also uncharacterized. It has been shown that while the naturally-occurring pigment in *Synechocystis* and *Arthrospira* OCP is 3’-hydroxyechinenone (3’-hECN), the protein may also be functionalized by other similarly-length ketocarotenoids, the palette of which will depend on species. Indeed, upon knock-out of the gene coding for the hydrolase converting ECN into hECN, *Synechocystis* OCP binds ECN [38], and likewise when the protein is overexpressed in *Synechocystis* cells, due to the low amount of hECN (≈ 1% of all ketocarotenoids) [11,39]. When expressed recombinantly in *E. coli* cells [33], *Synechocystis* OCP can be complexed with ECN, canthaxanthin (CAN) or zeaxanthin (ZEA), depending on the set of carotenoid-producing genes that are co-transformed with the gene coding for the OCP. This feature holds true for all OCP variants produced recombinantly in carotenoid-producing *E. coli* cells, including OCP1 from *Synechocystis*, *Arthrospira* and *Tolypothrix* [15,33], OCP2 from *Tolypothrix* and *Synechocystis* 7508 [15,16], and OCPX from *Scytonema* and *Synechocystis* 7508 [16]. To date, only *Tolypothrix* OCP was found to bind canthaxanthin (CAN) in the natural context, when overexpressed in *Tolypothrix* cells [30]. It is tantalizing to envision that functionalization of OCP by different types of carotenoids could enable regulation of its activity. Early studies showed that the relative populations of the carotenoid S_1_ and ICT excited-states depend on the carotenoid that functionalizes OCP, with virtually no ICT in *Synechocystis* OCP functionalized with the fully symmetric CAN (identical β-ionone rings at the two extremities), but up to 50% ICT in OCP functionalized with the non-symmetric ECN [30,40]. It was then proposed that the absence of the ICT state is a consequence of the fully symmetric nature of the CAN pigment, which is absent in ECN [30,40]. However, the P_1_ state was not identified at the time, so that it remains elusive whether or not the change in functionalizing carotenoid also affects the yield of this presumed photoproductive intermediate. Moreover, it is unknown if the rate of OCP^R^ accumulation and recovery, and the yield of intermediary photoproducts are influenced by the nature of the functionalizing carotenoid. Clearly, a thorough structure-function-dynamics study must be conducted to provide answers to all of the questions introduced above. Ideally, this study should be conducted on different OCP variants from different cyanobacterial strains to understand how subtle changes in the structure may affect the function, its regulation or both. For example, it was recently reported that despite a lower intracellular OCP concentration and a quasi-strict conservation of the amino acids lining the OCP carotenoid tunnel, the amplitude of the OCP-related PBS-fluorescence quenching is larger in the cyanobacterium *Planktothrix* PCC 7805 than in *Synechocystis* PCC 6803 [41], and likewise for the PBS-fluorescence recovery.

Here, we address these gaps in knowledge by performing a comparative structure-function study on the OCP1 from two different strains, viz. *Synechocystis* PCC 6803 (Syn-OCP), and *Planktothrix* PCC 7805 (Plankto-OCP). For both OCP1 variants, the kinetics of photoactivation, thermal recovery and PBS-fluorescence quenching were assessed in the native (non-tagged), N-tagged and C-tagged states, and with ECN or CAN as the functionalizing carotenoid. We observed that the presence of the his-tag at the C-terminus has a larger influence on photoactivation, thermal recovery and PBS-fluorescence quenching than its presence at the N-terminus. We found that the nature of the carotenoid influences the yield and characteristics of excited states, the ns-s dynamics of photoactivated OCP and the thermal recovery, leading to different rates of OCP^R^ accumulation, and of PBS-fluorescence quenching. As only the structures of ECN- and CAN-functionalized Syn-OCP [17,21] were available, we solved the Plankto-OCP structures in both the ECN- and CAN-functionalized states. At 1.4 – 1.8 Å resolution, these structures shed light on the molecular breathing motions that animate Plankto-OCP monomers and dimers, and point to subtle changes outside of the carotenoid tunnel explaining the peculiar properties of Plankto-OCP.

## Materials and Methods

### Construction of plasmids for *ocp* gene expression in *E. coli* cells

The plasmids pCDF-OCPSynCtag and pCDF-NtagOCPSyn featuring the *Synechocystis ocp* gene and coding for C-tagged Syn-OCP and N-tagged Syn-OCP, respectively, were described in [33]. To construct the plasmid *pCDF-*OCPSyn, needed to express the native (non-tagged) *Synechocystis ocp* (*srl963*) gene in *E. coli* cells, the nucleotides coding for the N-terminal his-tag in the plasmid pCDF-NtagOCPSyn were excised by mutagenesis using F-ocpSynNative (5’-ATAAGGAGATATACCATGCCATTCACCATTGACTCT-3’) and R-Duet (5’- CATGGTATATCTCCTTATTAAAGTTAAACAAAATTA-3’) as forward and reverse primers, respectively. To construct the plasmid containing the *Planktothrix agardhii* PCC *7805 ocp* (*PLAM_2315*) gene, a PCR was performed using the genomic DNA of *Planktothrix agardhii str. 7805* as a template and F-OCPPlank EcoR1 (5’- CGATGCGAATTCTTCATTTACAGTCGATTCAGCCC-3’) and R-OCPPlank Not1 (5’- CATTATGCGGCCGCCTTCCCCCTTAAATCACAAG-3’), as forward and reverse primers, respectively. Then, the PCR products were cloned into a pCDFDuet-1 plasmid featuring the nucleotides encoding for six-histidine tag at the N-terminus, which had previously been digested by EcoRI and NotI. This plasmid, named pCDF-NtagOCPPx, enabled expression of *Planktothrix* OCP with a his-tag at the N-terminus.

To synthesize non-tagged *Planktothrix* OCP, the nucleotides coding the six-histidine tag in the pCDF-NtagOCPPx plasmid were suppressed using F-OCPPxNoTag (5’- TTAATAAGGAGATATACCATGTCATTTACAGTCGATTCAGCCCGTGGG-3’) and R-Duet (see above) as forward and reverse primers, respectively, enabling to create the plasmid pCDF-OCPPx suited to express the native (non-tagged) *Planktothrix* OCP. Then, a nucleotide sequence encoding 6 histidine residues was added at the 3’end by mutagenesis using the pCDF-OCPPx plasmid as template and F-DuetOCPPx Ctag (5’- CACCACCACCACCACCACTAATTAATAAACGAATCTAATTTGATATAGC-3’) and R-DuetOCPPx Ctag (5’- GTGGTGGTGGTGGTGGTGACGAACTAAATTTAACAACTCTTTAGGTG-3’) as forward and reverse primers, respectively. Thereby, we obtained the plasmid pCDF-CtagOCPPx which was used to express *Planktothrix* OCP his-tagged at the C-terminus.

### Construction of plasmids for *frp* gene expression in *E. coli* cells

The plasmid used to express the *Synechocystis frp* gene in *E. coli* was described in [10]. To express the *Planktothrix agardhii* PCC *7805 frp* (*PLAM_2314*) gene in *E. coli*, the *frp* gene was amplified by PCR using forward (F-FRPPlank EcoR1: 5’- CGATGCGAATTCTCAAGTAAATGAGATTGAATG-3’) and reverse (R-FRPPlank Not1: 5’- TGCTTAGCGGCCGCAACTCAAATTGTTTTAAGAATCCCCG-3’) primers. The resulting fragment was cloned between the EcoRI and NotI sites of the pCDFDuet-1 plasmid which contains the nucleotides encoding for a six-histidine tag.

### Holo-OCP and FRP production, isolation and purification

The production and isolation of holo-OCP were previously described [33]. FRP isolation was described in [10]. To isolate non-tagged OCP, *E. coli* cells were resuspended in lysis buffer (40 mM Tris-HCl pH 8, 1 mM EDTA, 1 mM PMSF, 1 mM caproic acid, 1 mM benzamidic acid, 50 μg mL^-1^ DNAse) and broken using a French Press. Membranes were pelleted and the supernatant was loaded on a Whatman DE-52 cellulose column. The OCP was eluted using a gradient of 60-80 mM NaCl in 40 mM Tris-HCl pH 8. A second purification step was performed by hydrophobic interaction chromatography (HiTrap Phenyl HP column, GE Healthcare), where OCP was eluted in 40 mM Tris-HCl pH 8, 0.5M NaCl. The eluate was dialyzed overnight against 40 mM Tris-HCl pH 8 at 4°C (2 Liters).

### Absorbance measurements and kinetic analysis

Absorbance spectra of PBS and OCP and kinetics of OCP photoactivation (illumination with 5 000 µmol photons.m-^2^. s^−1^ of white light) and dark recovery were measured with a Specord S600 spectrophotometer (Analytic Jena) using 1 cm path-length cuvette. Experiments were performed at an OCP concentration of ∼4.8 µM, corresponding to an OD of 0.3 at 496 nm (calculated using an epsilon equal to 63 000 M^-1^.cm^-1^ [42]). Spectra were acquired in the 250–700 nm range for each time point. OCP^O^ photoactivation and OCP^R^ recovery were both monitored by measuring the increase and decrease in absorbance at 550 nm, respectively.

### Isolation of PBS and fluorescence measurements

The purification of PBS from *Synechocystis* PCC 6803 and *Planktothrix agardhii* PCC 7805 was performed as previously described [43]. The PBS-fluorescence quenching yield was monitored using a pulse amplitude-modulated fluorimeter (101/102/103-PAM, Walz). All measurements were made in a 1 cm path-length stirred cuvette. PBS quenching induced by holo-OCP was measured in 0.5 potassium phosphate buffer (pH 7.5) at 23 °C, under strong blue–green light exposure (900 µmol photons m^−2^ s^−1^). The PBS concentration used was 0.012 µM and the ratio of OCP to PBS was 40:1. OCP samples were pre-illuminated with 5 000 µmol photons m^−2^ s^−1^ of white light at 4°C. Samples in Pasteur pipettes were quickly frozen by immersion in liquid nitrogen. Fluorescence emission spectra at 77K were recorded using a CARY Eclipse fluorescence spectrophotometer fluorometer (Varian).

### Protein Separation

Proteins were analyzed by SDS-PAGE on 15% polyacrylamide gels in a Tris-MES buffer [44]. PBS samples were concentrated by precipitation with 10% (v/v) TCA prior to loading (equal protein quantities in each lane). 10 µL at a PBS concentration of 0.5 µM were deposited per well. The gels were stained by Coomassie Brilliant Blue.

### Femtosecond transient absorption spectroscopy (0 – 1 ns timescale)

Transient absorption measurements were performed using a Helios system from Ultrafast Systems. A short-pulse titanium sapphire oscillator (Mai Tai, Spectra Physics, 70 fs pulse length) followed by a high-energy titanium sapphire regenerative amplifier (Spitfire Ace, Spectra Physics, 100 fs, 1 kHz) provided the 800 nm beam, which was further split to generate: (1) a 532 nm pump pulse in the optical parametric amplifier (Topas Prime with a NirUVVis frequency mixer) and (2) a white light continuum (WLC) probe pulse in a sapphire crystal (430–780 nm). The remaining photons of the 800 nm probe-pulse were filtered just after white light continuum generation. The instrument response function (IRF) was estimated to be around 110 fs (FWHM). The pump-pulse diameter (FWHM) at the sample position was approximately 250 µm, for a pulse energy of 0.8 µJ. Absorbance over a 2 mm optical path was close to 0.7 at the excitation wavelength, corresponding to a concentration of 55 µM (or 1.9 mg/mL). The sample solution was stirred to keep the OCP solution fresh in the probed volume. The pump beam was depolarized to avoid anisotropy effects. To ensure that datasets are comparable to each other, they were all measured under identical conditions (at 22°C) on similarly-aged proteins during a single experimental session. The transient absorption data were corrected for the chirp of white light continuum by aligning kinetics according to a delay introduced by given amount of the material between the OCP sample and the sapphire crystal (used for WLC generation of the probe). For all datasets, the difference absorbance (ΔA) value obtained at the bleaching extremum (in both spectral and temporal dimension) was normalized to −1. Transient spectra were projected onto a 5 nm-spaced grid to get kinetic traces. The comparison of pre-exponential factors at 490 nm (bleaching band) allowed us to estimate the quantum yield of the formation of the various intermediates. We used for interpretation the most recent proposed model [32], whereby S_1_, ICT and S* are formed from S_2_ in parallel paths and decay mainly to S_0_ without any interconversion and with only small contribution of excited-state absorption at 490 nm. For P_1_ formation quantum yield, the positive absorbance contribution was taken into account [32]. Time constants determination and calculation of Decay Associated Difference Spectra (DADS or DAS) were achieved by global analysis using our custom-Python package [32] and the Glotaran software (https://glotaran.org). For the global analyses, a sum of four exponentials (S_2_, ICT, S_1_, S*) convolved by a Gaussian IRF (fixed to 110 fs) and an offset (representing long-lived photo-products with lifetime > 50 ns, namely P_1_) was used. Fitted results are reported in Table 1.

**Table 1:**
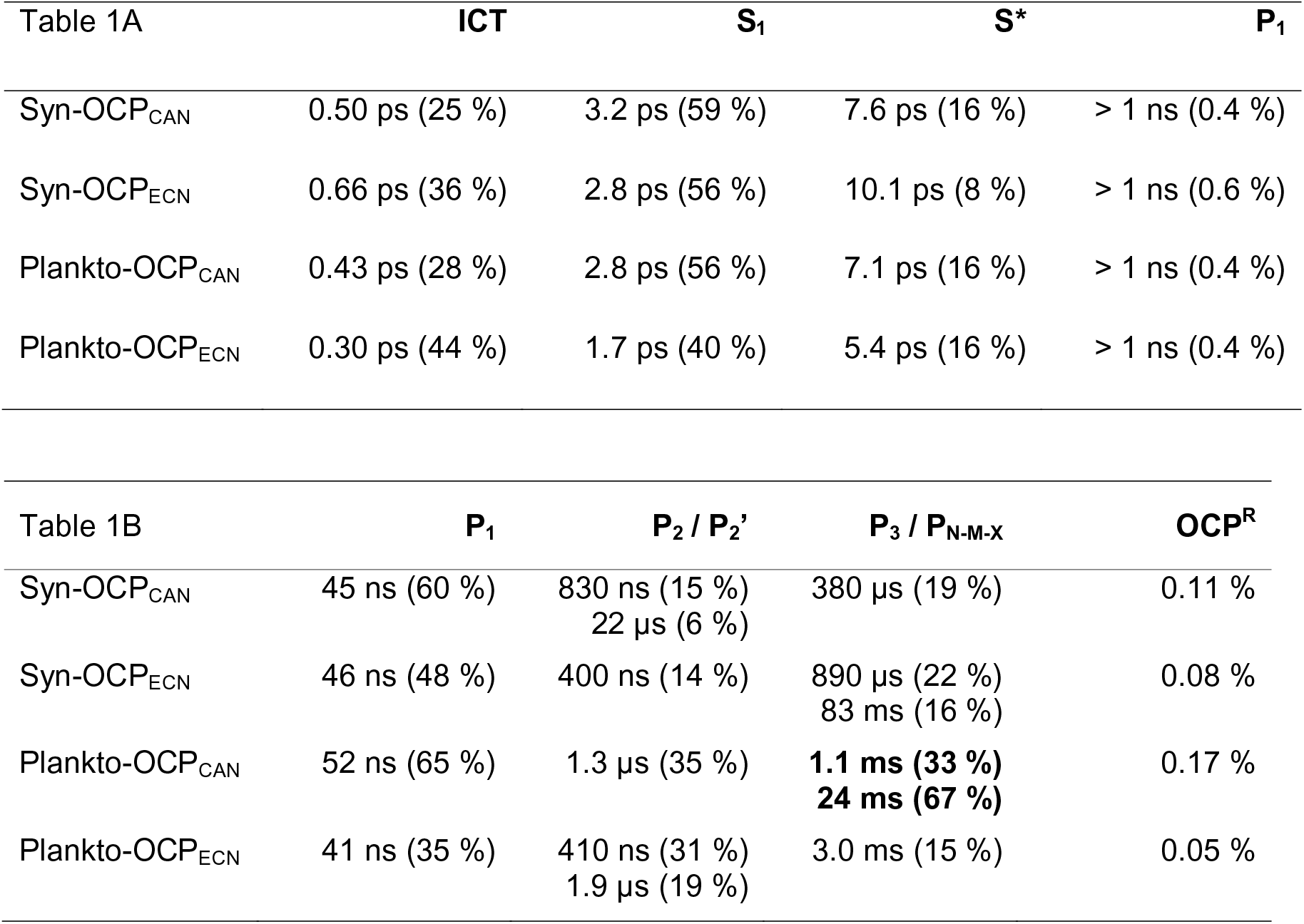
Lifetime, time constants and quantum yields obtained from femtosecond and nanosecond transient absorption experiments. (A) Lifetimes of ICT/S_1_/S*/P_1_ states derived from femtosecond transient absorption measurements (from data in Figure 5 and Figure S2), and estimated formation quantum yields derived from preexponential factors at 490 nm (the magnitude of the bleaching band of each DAS was extracted from the fitting procedure; for more details, see the Materials and Methods section and [32]). The standard error is 10%. P_1_ yields are corrected for its positive absorbance contribution at 500 nm. (B) Time constants and yields (percentages in brackets) derived from ns-s transient absorption measurements for P_1_, P_2_/P_2’_ and P_3_/P_N-M-X_. Only in the case of P_1_ are the value in brackets actual formation quantum yields; for P_2_/P_2’_ and P_3_/P_N-M-X_, they rather indicate the relative contribution of the corresponding exponential terms to the overall decay and growth components (indicated in bold). Percentages in the OCP^R^ column are formation quantum yields of the red absorbing intermediates remaining at 10 ms (determined by a comparative actinometry method with ruthenium complex; for details, see Material and Methods section and [46]).

### Nanosecond transient absorption spectroscopy (50 ns – 1 s timescale)

Measurements were performed with our custom apparatus described previously [45]. 532 nm nanosecond excitation pump pulses of 5 mJ energy were used (one pulse every 20 s for 100 ms kinetics, one pulse every 40 s for kinetics over 100 ms). The probe light from the Xenon lamp was filtered using a 550 nm interference filter (10 nm FWHM) placed before the sample. Scattered pump light was removed by a notch filter placed after the sample. For each experiment, a solution of OCP with absorbance close to 0.7 at the excitation wavelength (1 cm path-length, 11 µM or 0.39 mg/mL) was placed in a 10×10mm cuvette and thermalized at 22°C. No stirring of the sample was applied, enabling to stretch the time window covered by the experiments. Each set of measurements included 100 replicates of each of the five time-windows, together covering the 50 ns – 1 s time range. Recorded data were merged and projected onto a logarithmic grid. Stability of the protein was checked by its steady-state absorbance after each experiment. The formation quantum yield of OCP^R^ was determined using ruthenium as an actinometer [46]. The difference molar absorption coefficient at 550 nm for OCP^R^ was estimated using the molar absorption coefficient (ε) of OCP^O^ at 490 nm = 63000 cm^-1^.M^-1^) [42] and that determined for OCP^R^ after 100 % photo-conversion at 8°C (ε_550 nm_ ≈ 48000, 41 000, 47 000 and 40 000 cm^-1^.M^-1^ for Syn-OCP^R^_CAN_, Syn-OCP^R^_ECN_, Plankto-OCP^R^_CAN_ and Plankto-OCP^R^_ECN_, respectively). For each sample, data were fit globally over the five time-windows using a three-exponential model accounting for three different intermediates states and an offset. The latter is attributed to OCP^R^ and therefore can be used to estimate the yield. Fitted results are shown in Table 1.

### Crystallization

The N-tagged ECN-functionalized OCP from *Planktothrix aghardii* (Plankto-OCP_ECN_), purified on Ni-NTA and phenyl-sepharose columns, was further subjected to size exclusion chromatography under dim red light (HiLoad 16/600 Superdex 75 pg, GE Healthcare). The protein eluted as a unique peak in 50 mM Tris-HCl buffer pH 7.4, 150 mM NaCl. Crystallization conditions were screened manually, using as starting conditions those that afforded crystallization of Syn-OCP [17] and *Arthrospira maxima* OCP [8]. The gel-filtrated Plankto-OCP_ECN_ sample was concentrated to 5.2, 6.6 and 7.0 mg/mL (the protein concentration was determined based on the absorption at 495 nm, using an extinction coefficient of 63 000 M^-1^ cm^-1^ [42]) and crystallization trials were performed in 24-well Limbro plates using the vapor diffusion method in the hanging-drop geometry. Crystallization drops were set at 4°C by mixing 1 µL from the well solution with 1 µL of protein solution. The well solution, of 1 mL volume, was composed of 0.1 M or 0.2 M Bis-Tris pH 5.5 and increasing PEG 3550 concentrations (from 18% to 25%) were tested. Crystals appeared within 3 to 5 months in 0.2 M Bis-Tris pH 5.5, 20%-25% PEG 3550. Following this success, crystallization trials were optimized enabling growth of crystals in a few hours to few days at room-temperature (∼20 °C). Crystallization of Plankto-OCP_CAN_ was achieved using a protein concentration of 1.5 – 2 mg/ml in 50 mM Tris-HCl pH 7.4, 150 mM NaCl, a well solution composed of 0.1 M sodium acetate, pH 5 and 20-25% PEG4000, and by mixing these at 1:1 ratio in the crystallization drops.

### X-ray data collection and processing and structure determination

X-ray data were collected at 100 K from crystals cryoprotected by a short soak in the mother liquor complemented with 18-20 % glycerol and directly frozen in the nitrogen gas stream at the European Synchrotron Radiation Facility (ESRF, Grenoble, France), at the Swiss Light Source (SLS, Villigen, Switzerland) or on the MicroMax-007 HF diffractometer (Rigaku) installed at the Max Planck Institute in Heidelberg (MPI-HD). Specifically, we used : (i) ESRF-ID23EH1, for collection of the Plankto-OCP_ECN_ structure in the *P*2_1_ space group (1.7 Å resolution; λ=0.979 Å; beamsize: 30 (h) x 30 (v) µm^2^); (ii) ESRF-ID29, for collection of the Plankto-OCP_ECN_ in the *C*2 space group (1.7 Å resolution; λ=0.976 Å; beamsize: 30 (h) x 30 (v) µm^2^); (iii) SLS-X10SA (PXII), for collection of the Plankto-OCP_CAN_ structures in the *P*2_1_ space group (1.4 resolution; λ=0.99 Å; beamsize: 50 (h) x 10 (v) µm^2^); and (iii) the MPI-HD diffractometer, for collection of the Plankto-OCP_CAN_ structures in the *P*2_1_ and C2 space group (1.85 Å resolution, respectively; λ=0.99 Å; beamsize: 50 (h) x 10 (v) µm^2^). Data were collected with an oscillation range of 0.1 (ID29 and ID23-EH1), 0.2 (SLS-X10SA (PXII) or 0.25 degree (MPI-HD). All data were indexed using XDS [47], and scaled and merged using XSCALE [48].

### Molecular replacement and structure refinement

Phaser [49] was used to phase by molecular replacement (MR) the data collected on crystalline Plankto-OCP_ECN_ in the *C*2 space group, using as a starting model chain A from the Syn-OCP_ECN_ structure (PDB id: 3mg1, [17]). We used the CCP4 [50] buccaneer pipeline, based on Buccaneer [51], Parrot [52] and Refmac5 [53] for the initial *in silico* rebuilding of the *C*2 Plankto-OCP_ECN_ structure (76.2% and 92.8% identity and similarity with respect to Syn-OCP), resulting in a model characterized by Rfree and Rwork values of 29.63 and 26.17, respectively, and wherein 306 residues had been placed in sequence in two fragments corresponding to the NTD and CTD. Examination of this model revealed imperfections in loop building, which were corrected manually using the molecular graphics program Coot [54]. The *C*2 Plankto-OCP_ECN_ structure was thereafter refined by iterative cycles of reciprocal-space refinement using Refmac5 and manual model-building in real-space using Coot. Therefrom, phasing of the *C*2 Plankto-OCP_CAN_ data was achieved by rigid-body refinement with Refmac5, while that of the *P*2_1_ Plankto-OCP_ECN_ and Plankto-OCP_CAN_ structures was achieved by molecular replacement with Phaser, using as a starting model the refined *C*2 Plankto-OCP_ECN_ structure devoid of the carotenoid, waters and protein alternate conformations. Refinement again consisted of iterative cycles of reciprocal-space refinement using Refmac5 and manual model-building in real-space using Coot. Using *C*2 Plankto-OCP_CAN_ as the reference structure, Cα-Cα distance difference matrices were prepared using a custom-written script, and the hinge motions of domains within monomers, and of monomers within dimers were evaluated using the hinge_find.py script available at https://github.com/gawells/hingefind, inspired from the hinge_find.tcl script [55] available at http://biomachina.org/disseminate/hingefind/hingefind.html. The presence of tunnels in OCP, and notably the extent and volume of the carotenoid binding tunnel, was examined using Caver3 [56] and the dedicated PyMOL [57] plugin available at https://pymolwiki.org/index.php/Caver3). Porcupine plots, showing for each structure the direction and distance travelled by Cα atoms with respect to the *C*2 Plankto-OCP_CAN_ structure, were prepared using the modevectors.py PyMOL script available at https://pymolwiki.org/index.php/Modevectors. Figures were prepared with PyMOL unless stated otherwise. Data processing and refinement statistics are shown in Table 2. *Planktothrix* OCP structures have been deposited in the wwPDB under accession codes 7qd0, 7qcZ, 7qd1 and 7qd2.

**Table 2.**
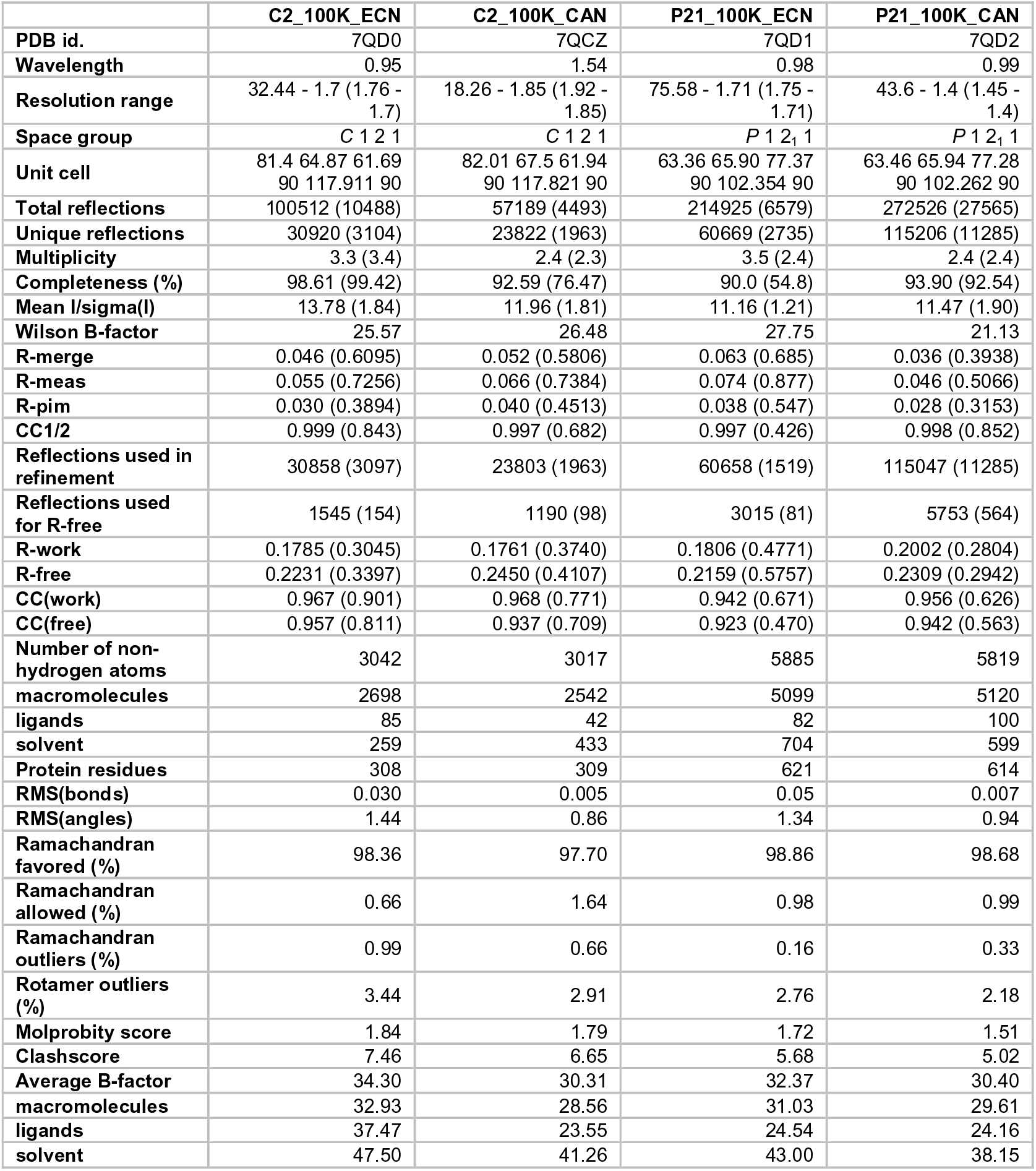
Data collection and refinement statistics. Statistics for the highest-resolution shell are shown in parentheses.

## Results

### Comparison of native (non-tagged) OCP from *Planktothrix aghardii* and *Synechocystis* PCC6803

The *Synechocystis* and *Planktothrix* species share the feature that only one *ocp* gene, classified in the OCP1 clade [16], and one *frp* gene are found in their genomes. The two OCP genes display 76.2% and 92.8% identity and similarity, respectively, with nearly all residues lining the carotenoid tunnel being conserved. We investigated whether or not the functionalizing carotenoid and his-tagging have an effect on the photoactivation and recovery kinetics of these two OCP, by expressing recombinantly the native (non-tagged), N-tagged and C-tagged *Planktothrix* and *Synechocystis ocp* genes in CAN or ECN producing *E. coli* cells (for details on constructs and on their expression and purification, see Materials and Methods section).

We first compared the spectral properties at 9.5 °C of the native (non-tagged) CAN-functionalized Syn-OCP and Plankto-OCP (Figure 2), viz. Syn-OCP_CAN_ and Plankto-OCP_CAN_. The two proteins display identical absorption spectra in the dark-adapted (orange) inactive state (OCP^O^), however slight differences are seen for the light-adapted (red) active state (OCP^R^) (Figures 2A, D and Supplementary Figure 1). Both OCP^R^_CAN_ spectra present an absorption maximum at 530 nm with a shoulder at 560 nm, which is slightly more pronounced in Syn-OCP^R^_CAN_ (Figures 2A and 2D). A difference positive absorbance maximum at 560 nm is observed in both OCP (Figures 2B and 2E). The normalization of the difference spectra time series on the 470 nm peak shows that the ΔA (560 nm) to ΔA (470 nm) ratio is higher in Syn-OCP^R^_CAN_ than in Plankto-OCP^R^_CAN_ (Figures 2C and 2F), pointing to a stronger absorption of the Syn-OCP^R^ state. This normalization also evidences a red shift in the spectra of the two OCP^R^ (from 545 - 550 to 560 nm) as they accumulate (Figures 2C and 2F), suggesting that a conformational change could occur that stabilizes the OCP^R^ state upon increase of its concentration by prolonged illumination.

**Figure 2:**
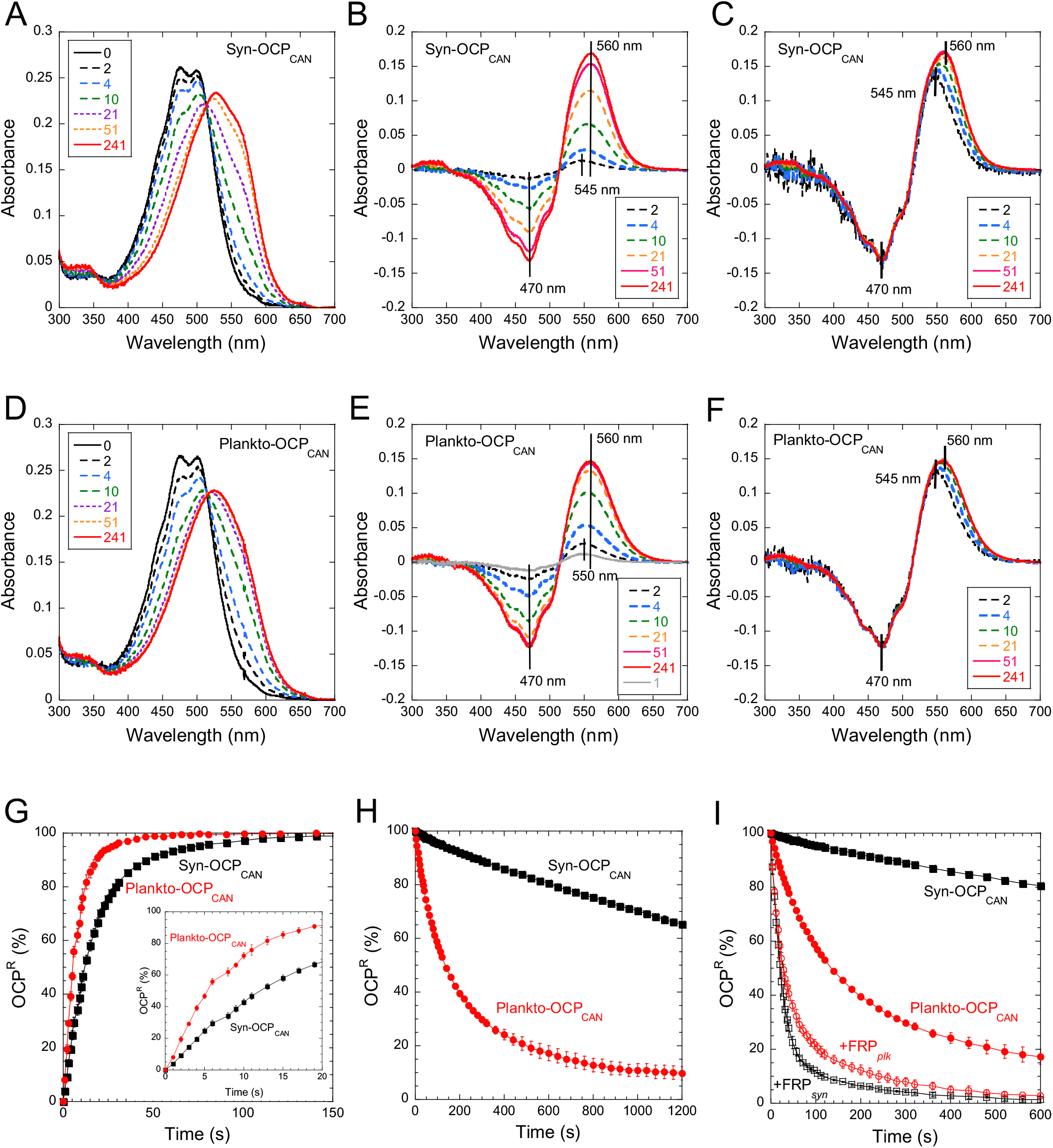
Photoactivation and recovery of native (non-tagged) CAN-functionalized *Synechocystis* and *Planktothrix* OCP. (A and D) Absorbance spectra of Syn-OCP_CAN_ (A) and Plankto-OCP_CAN_ (D) at different times of illumination. (B and E) Difference absorbance spectra derived from A and D respectively. (C and F) Difference absorbance spectra normalized at 470 nm derived from A and D. (G-I) Kinetics of photoactivation (G) and recovery (H-I) of CAN-functionalized Syn-OCP (black) and Plankto-OCP (red). The inset in G shows a close-up view on the first 30 s of illumination. In (I), the effect of the presence of FRP is shown. The FRP to OCP ratio was 1:1. The accumulation of OCP^R^ and its thermal deactivation were followed by increase and decrease of absorbance at 550 nm under illumination and in the dark, respectively. Illumination was performed with white light (5 000 µmol photons m^-2^. s^-1^) at 9.5°C. Error bars: standard deviation. Each curve represents the mean of three independent measurements, respectively.

We then assayed photoactivation and thermal recovery kinetics of Syn-OCP_CAN_ and Plankto-OCP_CAN_ (Figure 2G and H). Experiments were performed at 9.5°C, to minimize the negative contribution of thermal recovery to the steady-state accumulation of OCP^R^ – *i.e.*, to maximize OCP^R^ concentration in the photostationary equilibrium. Accumulation of OCP^R^ and recovery of the OCP^O^ state were monitored by following the rise and decrease in absorbance at 550 nm upon intense white-light illumination and subsequent transfer to darkness, respectively. Results in Figure 2 show that Plankto-OCP^R^_CAN_ not only accumulates faster than Syn-OCP^R^_CAN_ (initial slope is twice as high), but it also recovers faster the dark OCP^O^ state. Indeed, Plankto-OCP^R^_CAN_ recovers fully within the 20 min lapse of our experiment (Figure 2H), whereas only 30 % of Syn-OCP^R^ has reconverted to OCP^O^. The slow recovery kinetics at low temperature of Syn-OCP^R^_CAN_ is known, and shared by other OCP1s from *Arthrospira* and *Tolypothrix* [15,16,43], whereas the fast recovery kinetics of Plankto-OCP^R^_CAN_ is unprecedented for a member of the OCP1 clade and reminisces those displayed by members of the OCP2 [14,16] and OCPX [16] clades. In these clades, the faster OCP^R^-to-OCP^O^ recovery rate coincides with the inability to interact with FRP [15,16]. Therefore, we challenged a possible misclassification of Plankto-OCP as a member of the OCP1 clade by investigating whether or not its recovery is accelerated by the presence of FRP. For this purpose, *Synechocystis* and *Planktothrix* FRPs were expressed and purified, and assayed for their species-specific effect on the CAN-functionalized native versions of OCP. Figure 2I shows that the presence of FRP accelerates the recovery rate of Plankto-OCP_CAN_ (∼2.5 fold increase in the initial rate) although the observed acceleration is smaller than for Syn-OCP_CAN_ in presence of Syn-FRP (∼20 fold increase in the initial rate). These results confirm the correct assignment of Plankto-OCP to the OCP1 clade. They also show that the species-specific acceleration of OCP recovery by FRP is independent of the rate of the reaction in the absence of FRP.

### Influence of the his-tag on OCP photoactivation and recovery kinetics

We have recently shown that the location of the his-tag influences the photoactivation speed (initial rate) in *Synechocystis* OCP [32]. In particular, by using intermediary light intensity (∼100 µmol photons.m^-2^. s^-1^) we observed a more efficient accumulation of OCP^R^ in Ntag-Syn-OCP_ECN_ than in Ctag-Syn-OCP_ECN_ despite a comparable P1 formation quantum yield. Hence, we here asked whether or not presence of a his-tag, and its location at the N- or C-terminus, would influence the photoactivation and thermal recovery of Plankto-OCP and Syn-OCP. We compared results obtained from the N-tagged (Ntag-Syn-OCP_CAN_ and Ntag-Plankto-OCP_CAN_) and C-tagged (Ctag-Syn-OCP_CAN_ and Ctag-Plankto-OCP_CAN_) variants of these OCP to those of the native counterparts. It was consistently observed that the native proteins photoactivate faster than the his-tagged proteins, and that N-tagged OCP photoactivate faster than their C-tagged counterparts (Figure 3A and 3B). Nevertheless, when triggered with 5 000 µmol photons m^-2^. s^-1^ white light, the effect of his-tagging on the photoactivation rate was not severe Focusing next on the thermal OCP^R^ to OCP^O^ recovery, we found that it is delayed by presence of a his-tag in all tested OCP (Figures 3C and 3D), although the effect is more pronounced in Plankto-OCP_CAN_, which recovers faster than Syn-OCP_CAN_. Introduction of a his-tag at the N-terminus hardly affects the recovery of Plankto-OCP, whereas that at the C-terminus slows down the recovery by a factor of ∼6 (Figure 3D). In contrast, in Syn-OCP, his-tagging only has a slight effect on recovery.

**Figure 3:**
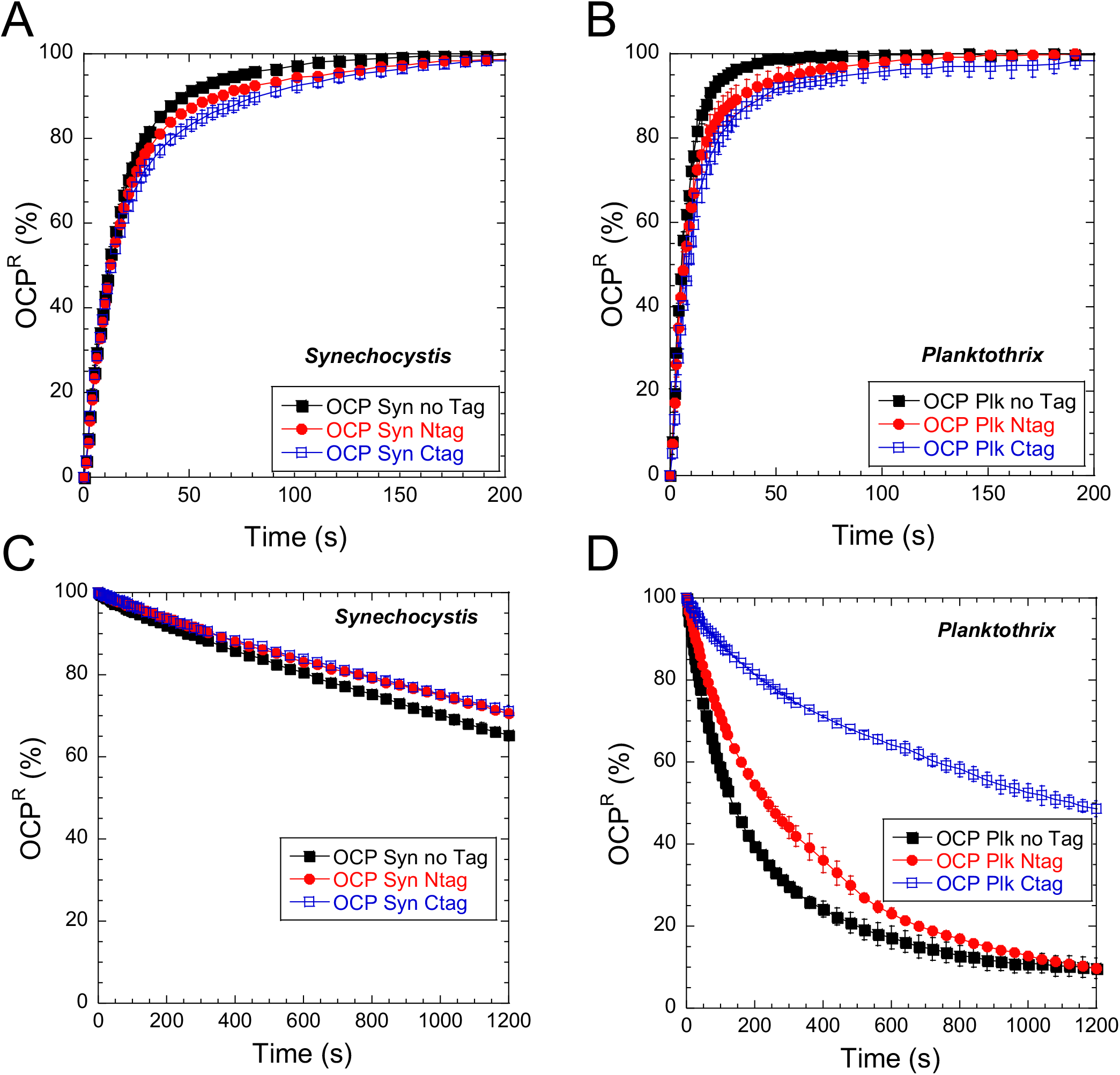
Presence of a his-tag has an impact on photoactivation and recovery. Effect of presence and position of his-tag on photoactivation (A-B) and recovery (C-D) of CAN-functionalized *Synechocystis* and *Planktothrix* OCP. The accumulation of OCP^R^ and its thermal deactivation were followed by increase and decrease of absorbance at 550 nm. The OCP were illuminated with white light (5 000 µmol photons m^-2^. s^-1^) at 9.5°C. Error bars: standard deviation. Each curve represents the mean of three independent measurements.

### Influence of the functionalizing carotenoid on OCP photoactivation and recovery kinetics

We then investigated the extent to which photoactivation and thermal recovery are affected by the type of functionalizing carotenoid. As it was found above that N-tagged and native OCP are the most similar in terms of photoactivation and recovery rates, we used N-tagged Syn-OCP and Plankto-OCP in the following assays. Irrespective of the species, OCP^R^ accumulation was found to be slower in ECN-functionalized than CAN-functionalized OCP (Figures 4A and 4B), while recovery is faster (Figure 4C and 4D). Indeed, the initial rate of OCP^R^ accumulation are twice as high in Ntag-Syn-OCP_CAN_ and Ntag-Plankto-OCP_CAN_ than in their ECN-functionalized counterparts (Figures 4A and 4B). The most straightforward explanation for these observations is that CAN stabilizes OCP^R^ and/or facilitate the carotenoid translocation during photoactivation. Alike their CAN-functionalized counterparts (Fig. 2A, D), the photoactivated Syn-OCP^R^_ECN_ and Plankto-OCP^R^ are spectrally similar, both presenting a maximum absorption at 510 nm (Figures 4E and H). In difference absorption spectra, the positive maximum is yet at 550.5 nm, *i.e.,* slightly blue shifted compared to the OCP_CAN_ counterparts (Figure 2 and Figures 4F, I), due to a reduced shoulder at 560 nm. It is notable that an increase in the ΔA (550 nm) to ΔA (470 nm) ratio is observed in the first 10 seconds of illumination of Plankto-OCP_ECN_ (Figure 4J); such an effect is not seen with Syn-OCP_ECN_ (Figure 4G).

**Figure 4:**
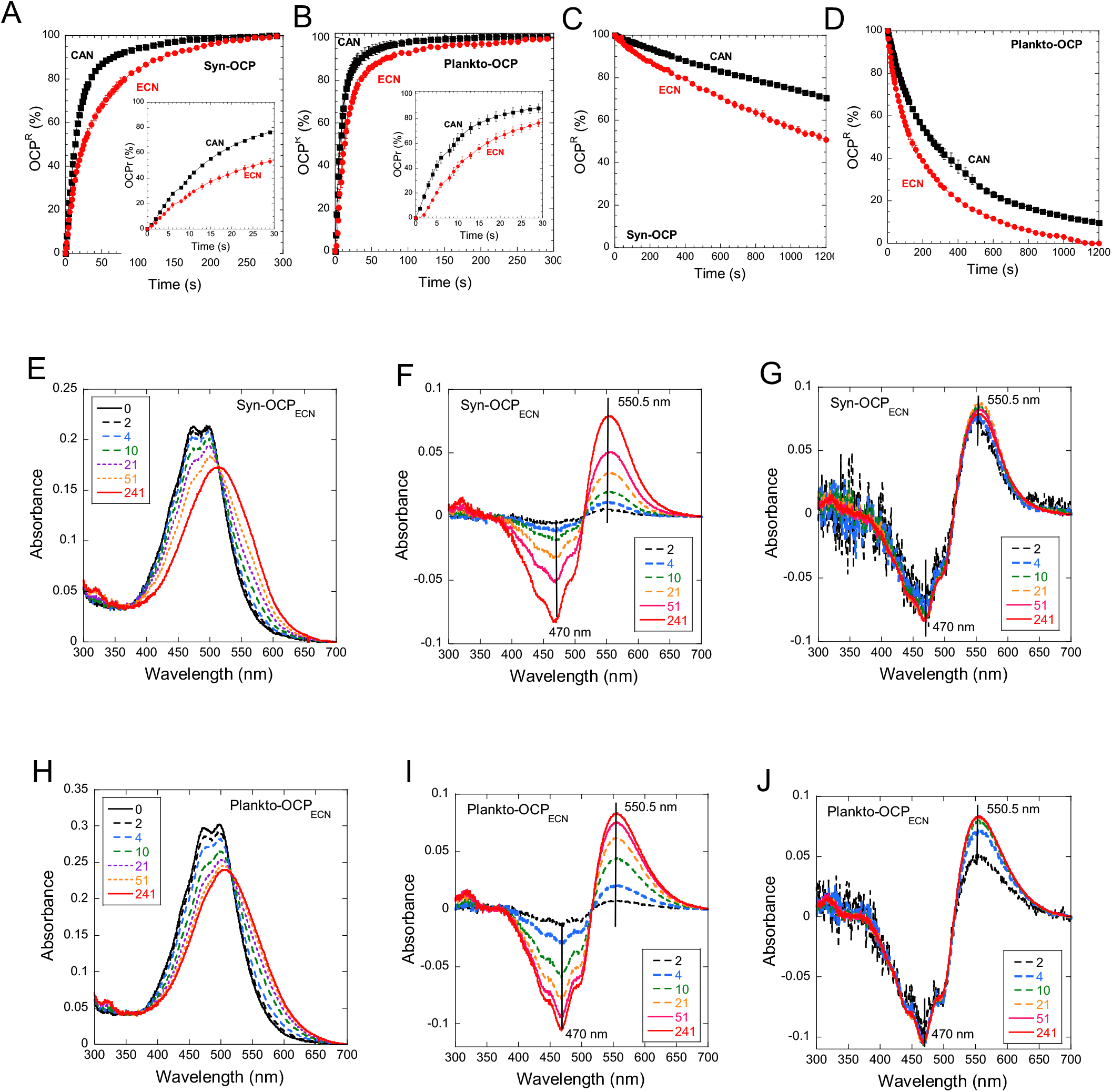
Effect of the nature of the functionalizing ketocarotenoid on photoactivation and recovery of N-terminally his-tagged *Synechocystis* and *Planktothrix* OCP. (A-B) Kinetics of photoactivation of Syn-OCP (A) and Plankto-OCP (B) functionalized with ECN (red) or CAN (black). The insets in A and B show a close-up view on the first 30 s of illumination. The OCP were illuminated with white light (5 000 µmol photons m^-2^. s^-1^) at 9.5°C. (C and D) Thermal recovery in darkness of ECN- (red) and CAN-functionalized Syn-OCP^R^ (C) and Plankto-OCP^R^ (D). The accumulation of OCP^R^ and its thermal recovery were monitored by following the increase and decrease of absorbance at 550 nm under illumination and in the dark, respectively. Error bars: standard deviation. Each curve represents the mean of three independent measurements, respectively. (E and H) Absorbance spectra of ECN-functionalized Syn-OCP (E) and Plankto-OCP (H) at different times of illumination. (F and I) Raw difference absorbance spectra derived from E and H, respectively. (G and J) Difference absorbance spectra derived from E and H, respectively, after normalization on the 470 nm band.

We inquired whether or not the increased photoactivation rate of CAN-functionalized Plankto- and Syn-OCP stems from changes in the carotenoid excited state dynamics. By carrying out fs-ns timescale transient absorption spectroscopy on the four OCP, we could estimate the primary quantum yields for the formation of the five intermediates occurring during the initial 100 fs-100 ps of the photoactivation cascade, *i.e.,* the S_2_, S_1_ and ICT excited-states, the S* state, and the first photoproduct P_1_ (see Material and Methods section and [32]). The formation and decay of these states in CAN- and ECN-functionalized Plankto-OCP (Figure 5) and Syn-OCP (Supplementary Figure 2) were monitored by recording and globally-fitting femtosecond transient spectra collected at different time delays following a 110-fs pulse excitation at 532 nm (see the Materials and Methods section for further details).

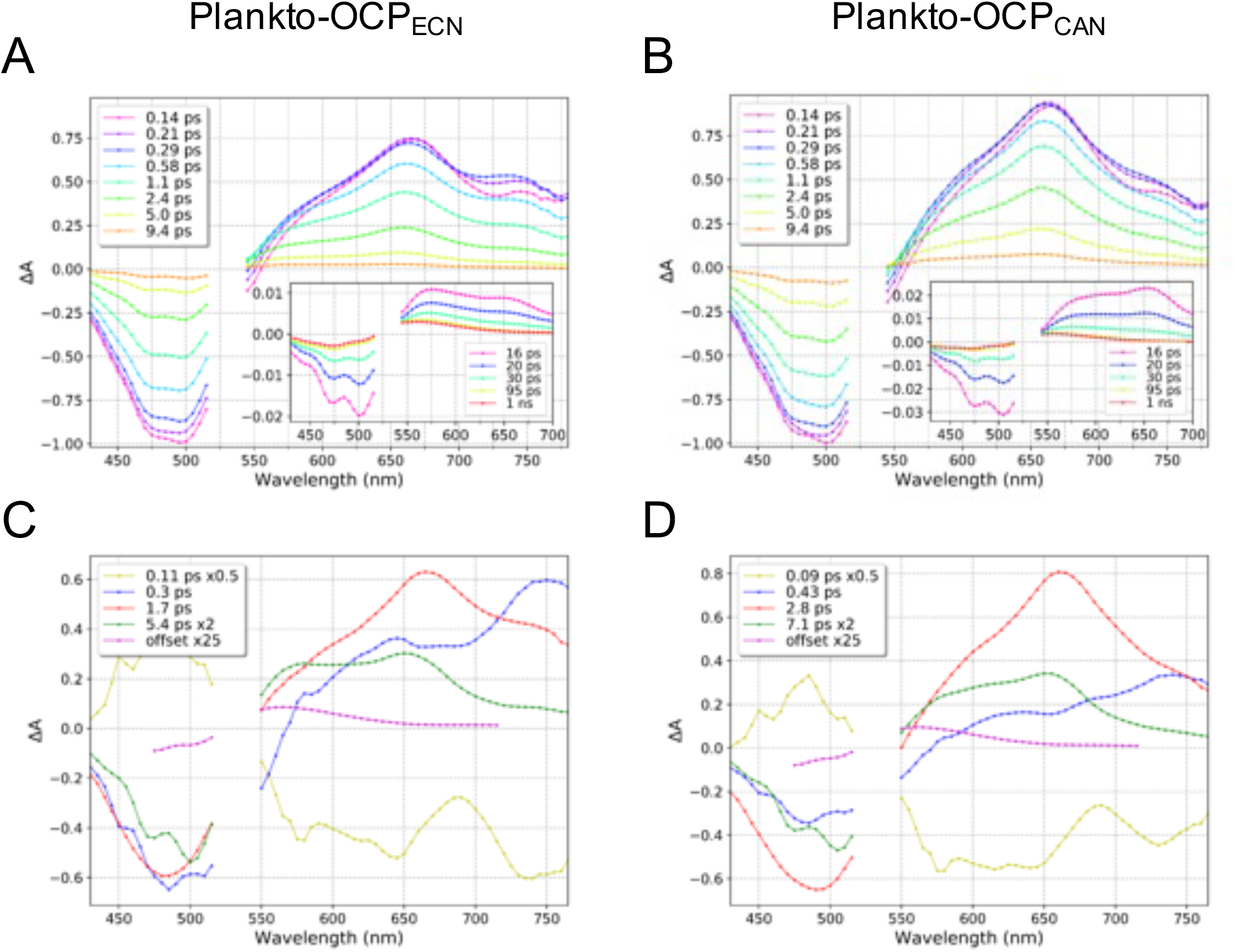

In agreement with previous reports, we found that 0.15 ps after excitation, the S_2_ excited state has already started to decay. Transient absorption spectra are characterized by a ground state bleaching (GSB) negative band with an extremum at ≈ 500 nm, indicative of OCP^O^ depopulation, and by positive absorption (ESA) bands for the S_1_, ICT and S* states peaking at ≈ 660 nm, ≈ 750 nm and ≈ 575 nm (shoulder), respectively. All excited- and vibrationally-hot ground-states have decayed by the 30 ps time-delay, and only a broad positive band centered at ≈ 560 nm, previously assigned to the photoproduct P_1_ [19,32], can be seen at the 95 ps time delay. P_1_ shows no spectral evolution in our experimental time window, *i.e.,* up to 1 ns time delay.

Four exponential components (S_2_, ICT, S_1_ and S*) and an offset (P_1_) were accounted for in the global decay analysis, enabling to extract the decay associated spectra (DAS) and lifetimes of the S_2_, ICT, S_1_, S* and P_1_ states in ECN- and CAN-functionalized Plankto-OCP (Figure 5C and 5D) and Syn-OCP (Supplementary Figure 2C and D). Note that as our resolution is about 110 fs, the DAS associated with the shortest time-constants is estimated from a convolution of the S_2_ decay with the rise of the other excited-state signals. The difference spectrum obtained ≈ 30 ps post-excitation (offset value in the sum of exponentials) is attributed to the P_1_ intermediate state. Using the GSB kinetics, and assuming that the S_1_, ICT and S* states all form from S_2_ and parallelly decay to the ground state S_0_, we could estimate the quantum yields of each state, including the P_1_ state. In agreement with previous reports, the ps excited-state dynamics and yields are similar for the four tested OCP, with similar DAS observed for the ICT (blue), S_1_ (red), S* (green) and P_1_ (magenta) states, respectively (Figure 5C and 5D). However, a careful comparison of the DAS of CAN- and ECN-functionalized OCP reveals that the absorbance amplitudes above 700 nm of the S_1_ and ICT states are higher in the ECN-functionalized OCP, reflecting a more pronounced ICT character in these states. The lifetimes derived from the global fitting of our data are in agreement with previous reports [58], viz. ∼0.10ps (± 0.01ps), ∼0.5ps (± 0.2ps), ∼2.6ps (± 0.9ps) and ∼7.5ps (± 2.5ps) for the S2, ICT, S1 and S* states of the four OCP, respectively (Figure 5 and Table 1). We note that differences are seen for the lifetimes of the S_1_ and S* states in the ECN and CAN-functionalized OCP, which are most pronounced in Plankto-OCP. As expected from the DAS and the literature [40], and irrespective of the considered OCP variants, the formation QY for the ICT and S* states are lower and higher in CAN-functionalized OCP, respectively. The observed P_1_ yield is yet similar in the four tested OCP (≈ 0.5 ± 0. 1 %), in agreement with recent results from us [32], and others [19,23].

The above-described fs-ns transient spectroscopy data exclude the hypothesis that the type of carotenoid or OCP scaffold significantly influences the yield of P_1_. Hence, we next examined the ns-s dynamics by performing nanosecond transient absorption experiments, whereby a nanosecond laser pulse is used to trigger excitation and the photoactivation outcome is probed in the 50 ns – 1 s time window by monitoring changes in the absorbance at 550 nm (Figure 6). Indeed, on these timescales, the maximum difference-absorbance varies between 563 nm, characteristic of the P_1_ state, and 550 nm, signing for the nascent OCP^R^ state. Intermediate states were assigned on the basis of earlier reports, with the P_1_, P_2_-P_2_’ and P_3_(P_N_)/P_M_/P_X_ states displaying lifetimes of ∼ 50 ns, ∼ 0.5-10 µs, and ∼ 1-100 ms, respectively. Recall that these states were proposed to be associated with (i) rupture of H-bonds between the carotenoid and the protein scaffold (P_1_); (ii) rearrangement of the carotenoid at the NTD/CTD interface (P_2_/P_2_’) and translocation of the carotenoid from the NTD/CTD interface into the NTD (P_3_); and (iii) NTE and CTT detachment (P_M_) followed by dissociation of the two domains (P_X_), respectively. A last conformational change thence occurs, yielding from P_X_ the metastable OCP^R^ (Figure 1). This last step could correspond to the repositioning of the CTT on the CTD-half of the carotenoid tunnel [34]. Note that partial recovery of the OCP^O^ state occurs at all steps (see Figure 1), explaining the decrease in absorbance at 550 nm over the probed time window.

**Figure 6:**
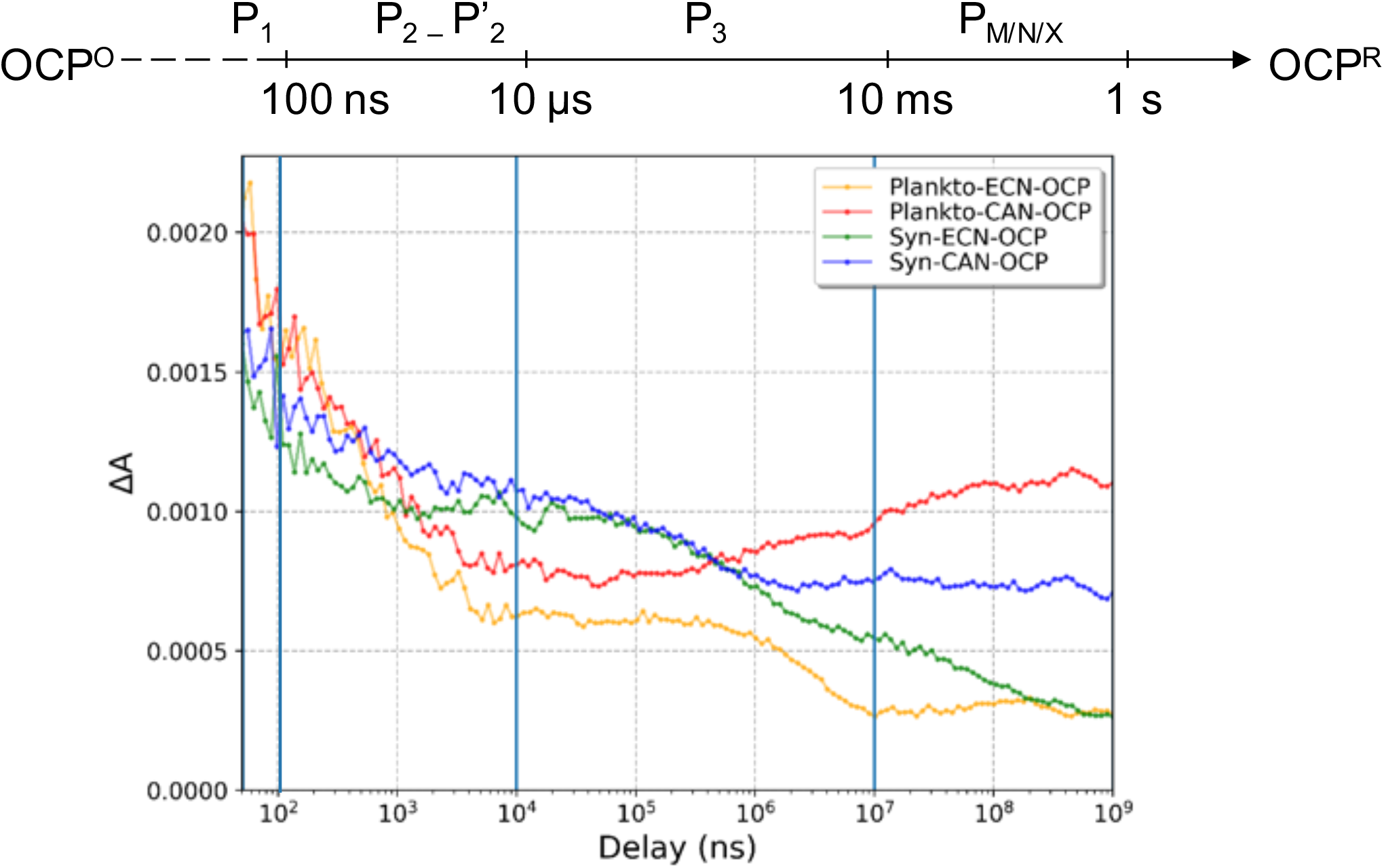
Nanosecond-second dynamics in CAN- and ECN-functionalized Plankto-OCP and Syn-OCP. Time evolution (50 ns to 1 s) of the difference absorption signals at 550 nm recorded after excitation by a 532 nm ns-pulse on Plankto-OCP_ECN_ (yellow), Plankto-OCP_CAN_ (red), Syn-OCP_ECN_ (green) and Syn-OCP_CAN_ (blue). The intermediate states proposed [12,19,23] to underlie the observed absorption changes are indicated at the top of the figure. Measurements were carried out at 22°. Each set of measurements included 100 replicates of each of the five merged time-windows, together covering the 50 ns – 1 s time range.

Clear differences are seen between the four tested OCP in these experiments. First, irrespective of the carotenoid, the starting difference absorbance signal (at 50 ns) is higher for Plankto-OCP (∼0.002) than for Syn-OCP (∼0.0015), suggesting a higher P_1_ yield (Figure 6). This observation contradicts the above assumption that the P_1_ yield is the same for all investigated OCP, but can be rationalized by recalling that (i) in the fs-ns experiments, the GSB band at 490 nm is used to estimate the P_1_ yield, instead of the characteristic positive absorption band at 563 nm, in the ns-s experiments; and most importantly (ii) a large fraction of P_1_ reverts to the dark-adapted OCP^O^ state. Thus, the higher P_1_ yield observed for Plankto-OCP at the start of ns-s transient absorption experiments (50 ns) could be related to a reduced recovery from P_1_ of the Plankto-OCP^O^ state, rather than from an increased P_1_ yield. Also irrespective of the functionalizing carotenoid, the difference absorptions of Plankto-OCP and Syn-OCP drop by about 60 % and 40 % on the ns - µs time scale, respectively, with a larger drop in Plankto-OCP signal that could underlie sub-optimal translocation of the carotenoid into the Plankto-NTD, compared to the Syn-NTD. The most important differences between Plankto-OCP and Syn-OCP are visible in the µs to ms timescale, whereas differences between CAN-and ECN-functionalized OCP concentrate in the ms-s time window. Thus, both carotenoid translocation and NTE/CTT detachments seem to be affected by the change in protein scaffold, however, it is domain dissociation that is most affected by a change in the functionalizing carotenoid. This step appears to be faster and more efficient in CAN-functionalized OCP, with a slight increase in ΔA (550 nm) being visible after 1 ms, whereas recovery to the initial state is higher for ECN-functionalized OCP, as evidenced by the decrease of ΔA (550nm). The observation of this decline being present only in ECN-functionalized OCP is in line with the observation that CAN-functionalized OCP can be photoactivated more efficiently (Figure 4). In the case of Plankto-OCP_CAN_, we observe faster domain separation and no recovery to OCP^O^ on the µs-s timescale, suggesting 100% conversion efficiency from P_3_ to final OCP^R^. The larger difference absorption signals observed for Plankto-OCP (most notably Plankto-OCP_CAN_) on the 10 ms - 1 s time window mirror the differences in photoactivation efficiency observed in stationary irradiation experiments.

Multiscale exponential fitting was carried out to extract the lifetimes of the ns-s intermediate states (Table 1). The lifetime of the P_1_ state was found to be almost independent of the OCP variant, *i.e.,* ∼50 ns. For the P_2_-P_2’_ states (*i.e.,* on the 50 ns to 10 µs time scale), data were fit by one or two exponentials, depending on the case, yielding lifetimes in the order of 0.5-2 µs. Likewise, for the P_3_/P_N_/P_X_ states (*i.e.,* on the 0.1-10 ms time scale), either one or two exponentials were required to fit the data, yielding lifetimes in the order of 0.1-1 ms. Note that these lifetimes are in good agreement with those reported earlier based on experiments carried out on the C-tagged Syn-OCP_ECN_ [19,23]. Our data establish that the OCP^R^ yield is higher for CAN-functionalized OCP, and that Plankto-OCP photoactivates more efficiently. Thus, the differences observed in the photoactivation efficiency of CAN- and ECN-functionalized Plankto- and Syn-OCP stem from evolutions observed during the (comparatively-slow) carotenoid translocation, NTE/CTT detachment and domain dissociation steps – but not from changes in the excited-state dynamics of the OCP-embedded carotenoid. The experimental setup did not allow to evaluate a lifetime for OCP^R^, but previous reports have pointed to the second timescale [23].

### Interaction between OCP and the phycobilisome

We inquired the energy-quenching performance of the various ECN- and CAN-functionalized OCP, focusing on OCP and PBS from *Synechocystis* and *Planktothrix.* The PBS from *Synechocystis* has been well characterized in several laboratories, including ours [43,59], yet no information was available regarding the *Planktothrix* PBS. Hence, a prerequisite was to characterize it biochemically and spectroscopically (Supplementary Figure 3). We found that similar to *Synechocystis* PBS, *Planktothrix* PBS consists of a core (formed by three cylinders containing four allophycocyanin (APC) trimers) from which radiate six rods constituted of three phycocyanin (PC) hexamers.

With this characterization in hand, we pursued the investigation of native Syn-OCP and Plankto-OCP energy-quenching activities, focusing first on the effect of the functionalizing-carotenoid (Figure 7) in species-specific OCP/PBS complexes. The decrease of PBS fluorescence was followed using a PAM fluorimeter during incubation of PBS under strong blue-green light and in presence of pre-photoactivated OCP. Under these conditions, a faster and larger decrease of fluorescence is suggestive of a stronger OCP-PBS interaction. We found that native Syn-OCP_CAN_ and Syn-OCP_ECN_ induce similar Syn-PBS fluorescence quenching (∼75 % of PBS fluorescence is quenched after 300 s), despite a slightly lower initial rate for native-Syn-OCP_CAN_ suggesting a weaker binding to the PBS (Figure 7A). This hypothesis was confirmed by the eight times faster PBS fluorescence recovery observed for native-Syn-OCP_CAN_, compared to native-Syn-OCP_ECN_ (Figure 7B). The nature of the functionalizing-carotenoid also had an effect on native-Plankto-OCP quenching efficiency and on the Plankto-PBS fluorescence recovery rate. PBS fluorescence quenching was twice more efficient with native-Plankto-OCP_ECN_ than with native-Plankto-OCP_CAN_ (75 and 40 % of fluorescence quenching after 300 s incubation, respectively) and fluorescence recovery was nearly three times faster for native-Plankto-OCP_CAN_ than native-Plankto-OCP_ECN_. These results support the hypothesis that CAN-functionalized OCP display a reduced affinity for the PBS.

**Figure 7:**
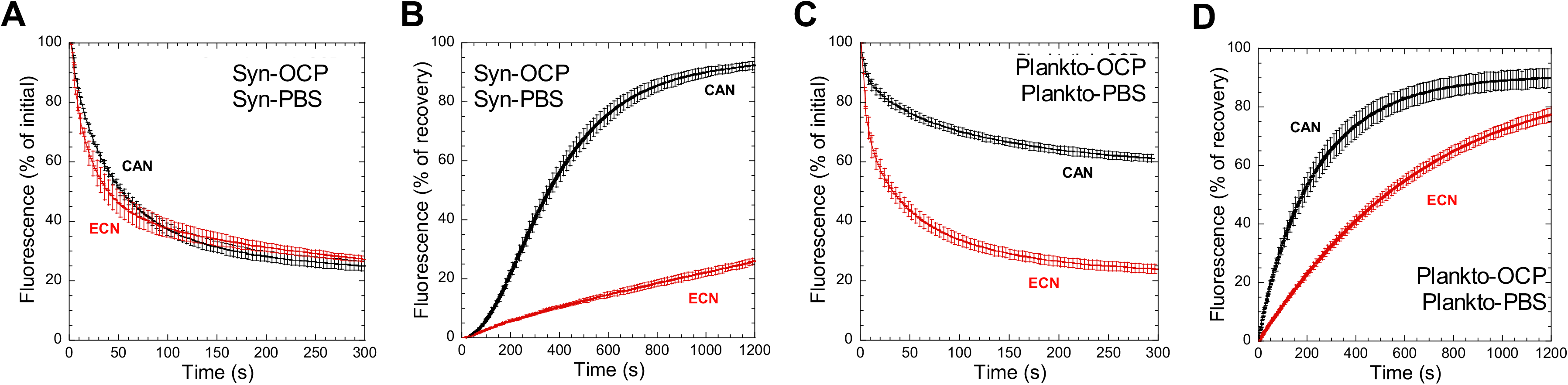
Effect of ketocarotenoid on OCP-PBS interaction. (A, C) Syn-PBS (A) and Plankto-PBS (C) fluorescence-quenching induced at 23° C in 0.5 M phosphate buffer by non-tagged (native) CAN- (black) and ECN- (red) functionalized Syn-OCP (A) and Plankto-OCP (C). For these experiments, OCP was pre-photoactivated by illumination with a strong white light (5 000 µmol photons m^-2^. s^-1^) at 4°C, then mixed with Syn-PBS or Plankto-PBS at a 40:1 ratio, and kept under illumination by strong blue-green light (900 µmol photons m^-2^. s^-1^) during the lapse of the measurement. (B, D) Dark recovery of fluorescence in *Synechocystis* (B) and *Planktothrix* (D) phycobilisomes. 100% of fluorescence in all graphs is the initial fluorescence of phycobilisomes without quenching. Error bars: standard deviation. Each curve represents the mean of three independent measurements.

We also investigated the effect of a his-tag, present either at the N- or C-terminus, on the species-specific PBS-fluorescence quenching efficiency and recovery rate (Figure 8). We found that irrespective of the species (*i.e.,* for both Syn-OCP + Syn-PBS and Plankto-OCP + Plankto-PBS), his-tagged OCP are more efficient at inducing PBS-fluorescence quenching, with C-tagged OCP further surpassing the N-tagged variants (Figure 8A and 8B). This effect is most clear when considering the native, N-tagged and C-tagged Plankto-OCP/PBS complexes. A possible rationalization could come from the higher stabilization of the OCP^R^ state upon his-tagging at the C-terminus. However, this stabilization is not sufficient to explain the observed drastic difference in PBS fluorescence-recovery rates. Indeed, only up to 20 % of the initial PBS fluorescence is recovered after 20 min incubation in the dark with the C-tagged Plankto-OCP (Figure 8D, E), which is at variance with the full recoveries observed when native or N-tagged Plankto-OCP and Syn-OCP are used. Thus, we favor the hypothesis that atop of stabilizing OCP^R^, the C-terminal his-tag also stabilizes the OCP^R^/PBS complex, explaining the faster quenching and slower recovery.

**Figure 8:**
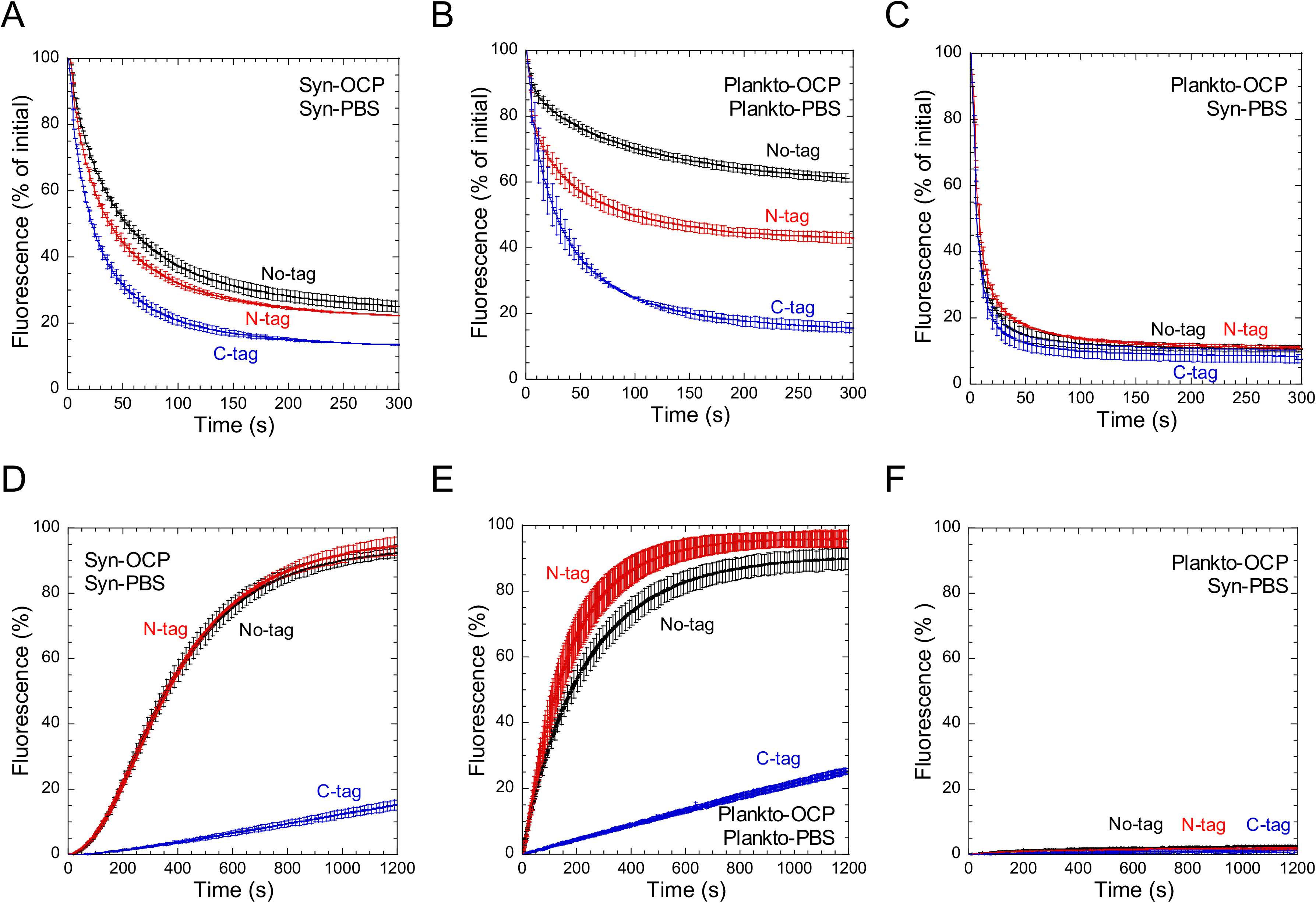
Effect of the presence and position of a his-tag on OCP-PBS interaction. (A-C) *Synechocystis* (A, C) and *Planktothrix* (B) PBS fluorescence quenching induced at 23°C in 0.5 M phosphate buffer by native (non-tagged; black), N-tagged (red) and C-tagged (blue) CAN-functionalized *Synechocystis* (A) and *Planktothrix* (B, C) OCP. For these experiments, OCP was pre-photoactivated by illumination with a strong white light (5 000 µmol photons m^-2^. s^-1^) at 4°C, then mixed with Syn-PBS or Plankto-PBS at a 40:1 ratio, and kept under illumination by strong blue-green light (900 µmol photons m^-2^. s^-1^) during the lapse of the measurement. (D-F) Dark recovery of fluorescence in *Synechocystis* (D, F) and *Planktothrix* (E) PBS. 100 % of fluorescence in all graphs is the initial fluorescence of phycobilisomes without quenching. Error bars: standard deviation. Each curve represents the mean of three independent measurements.

In the past, we demonstrated that *Synechocystis*, *Arthrospira* and *Anabaena* OCP bind more strongly to the *Synechocystis* PBS than to the *Anabaena* or *Arthrospira* PBS [43], with *Synechocystis* OCP binding comparatively weakly to the PBS from other species [43]. Hence, we asked whether or not these conclusions hold for *Planktothrix* OCP and PBS as well. We found that irrespective of the presence and position of the his-tag, *Planktothrix* OCP binds more strongly to *Synechocystis* PBS than to *Planktothrix* PBS, with all Plankto-OCP inducing a faster and more efficient fluorescence quenching of Syn-PBS (Figure 8B) compared to Plankto-PBS (Figure 8C). Furthermore, the *Synechocystis* PBS fluorescence recovery was virtually null, indicating that the complex formed by Plankto-OCP and Synechocystis-PBS is highly-stable at 0.5 M phosphate (Figure 8F). Contrastingly, Syn-OCP was unable to induce *Planktothrix* PBS fluorescence even at higher phosphate concentrations (till 1.4 M phosphate), suggesting that it is able to interact only with *Synechocystis* PBS.

### Structures of CAN- and ECN-functionalized *Planktothrix* OCP reveal new features

The crystal structures of *Anabaena* PCC 7120 (PDB id: 5hgr; also referred to as *Nostoc*) and *Tolypothrix* PCC 7601 (PDB id: 6pq1; also referred to as *Fremyella diplosiphon*) OCP have been solved in the CAN-functionalized states, while that of *Arthrospira maxima* (PDB id: 5ui2; also referred to as *Limnospira*) was determined in hECN functionalized states. Only the structure of *Synechocystis* PCC 6803 OCP was solved in both the CAN- and the ECN-functionalized states (PDB ids: 4xb5 and 3mg1, respectively), offering a first basis to rationalize the more efficient photoactivation observed for CAN-functionalized OCP in transient absorption (Figure 6) and photostationary experiments (Figure 4). Structural information on *Planktothrix* OCP was yet absent, preventing identification of the structural features that could underlie its functional characteristics – notably, the thwarted translocation of carotenoids into the NTD, revealed by transient absorption spectroscopy, and the increased photoactivation rate and lower stability of its OCP^R^ state, revealed by photo-stationary experiments (Figures 4 and 6).

Hence, we set to characterize the structure of Ntag-Plankto-OCP in the CAN- and ECN-functionalized states. The structure was solved, for each of these, in two space groups, viz. *C*2 and *P*2_1_, revealing a remarkable conservation of the secondary structure (Figure 9A). The *C*2 structures were solved at slightly higher resolution (1.4 and 1.7 Å resolution for Ntag-Plankto-OCP_CAN_ and Ntag-Plankto-OCP_ECN_, respectively) than the *P*2_1_ structures (1.8 Å resolution for both OCP) (Table 2). In the *P*2_1_ space group, the asymmetric unit features the dimer that has been observed in previous structures [8,17,21,30,58,60,61]. Note that in the case of Syn-OCP_CAN_ (4xb5), Anabaena-OCP_CAN_ (5hgr) and *Tolypothrix-*OCP_CAN_ (6pq1), the dimer is crystallographic, *i.e.,* the dimerization interface perfectly aligns with a crystallographic axis; hence, the asymmetric unit confusingly features a monomer and application of symmetry operations are needed to reveal the dimer. In Syn-OCP_ECN_ and Arthrospira-OCP_ECN_, however, the asymmetric unit features a dimer. In the case of Syn-OCP, this dimer was recently shown to naturally occur *in vitro* [62,63] with a dissociation constant in the order of 14-17 µM, depending on reports [16,63]. The large dimerization interface (1046.7 and 1084.7 Å^2^ of buried surface area (BSA) in the *P*2_1_ structures of Plankto-OCP_CAN_ and Plankto-OCP_ECN_) is mainly supported by N-terminal helices αA (NTE) and αB, which contribute ∼70 % of the BSA, with minor contributions from helix αH (∼ 20 % of the BSA) and the αE-αF and β2-β3 loops (∼ 10 % of the BSA) (Figure 9B, C and Supplementary Table S1). Interestingly, the relative contributions of αA and αB to the BSA at the dimerization interface vary depending on the functionalizing carotenoid, amounting to 37 and 33 % in Plankto-OCP_CAN_, and 27 and 40 %, in Plankto-OCP_ECN_, respectively. Additionally, two H-bonds fixing αA and αB from facing monomers in the Plankto-OCP_CAN_ dimer (viz. R9(NH2)–Q30(O) and R9(NH1)–L31(O)) are suppressed in the Plankto-OCP_ECN_ dimer (Figure 9D and Supplementary Table S1). Thus, the NTE (αA) is less constrained (BSA decreases by 25%) at the dimerization interface in the *P*2_1_ Plankto-OCP_ECN_ structure than in the *P*2_1_ Plankto-OCP_CAN_ structure, but the dimer is more tightly packed, with 25, 41 and 52 % increase in the BSA contributed by αB (largest contributor to the dimerization interface in all structures), and the αE-αF and β2-β3 loops, respectively (Supplementary Table S1). The rest of the H-bonding network at the dimerization interface is overall conserved in the *P*2_1_ structures, with 3 H-bonds between αB and the facing αH (N14(OD1)–A133(N), T15(O)–N134(ND2), T17(OG1)–N134(ND2)), one H-bond between αA and the facing αEF loop (either D6(OD2)– T90(OG1) or D6(OD2)–N88(ND2) in the Plankto-OCP_CAN_ and Plankto-OCP_ECN_ structures, respectively) and a salt-bridge between facing αB residues (D19(OD2)-R27(NE)) (Supplementary Figures 4 and 5).

**Figure 9.**
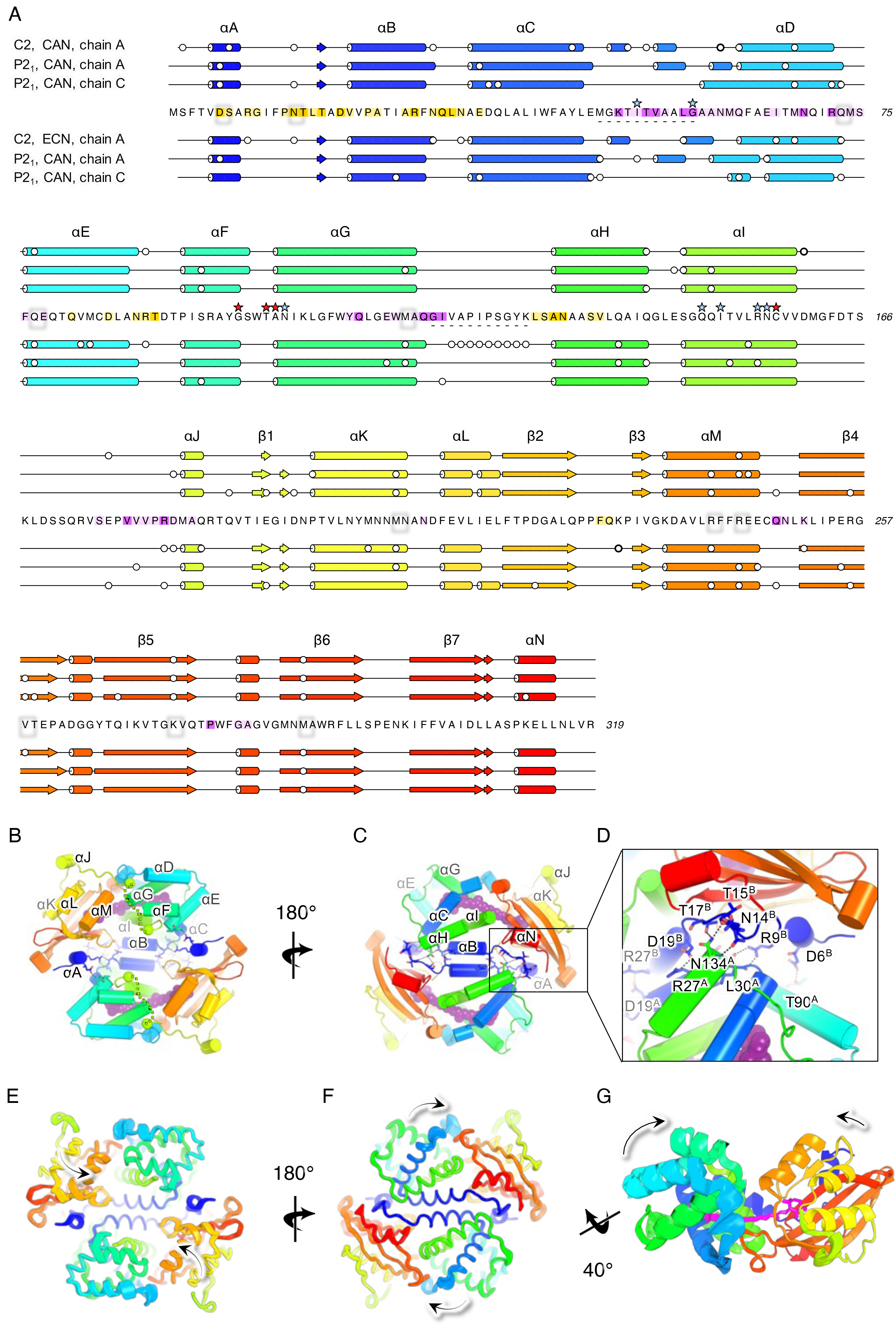
All OCP structures feature a dimer, including those of Plankto-OCP. (A) The secondary structure of OCP is overall well conserved amongst Plankto-OCP structures obtained in different space groups and with different pigments. Residues involved in the dimerization are highlighted in yellow, whereas residues involved in the formation of interface X are highlighted in purple. Dark and light colouring indicate residues involved in polar and van der Walls interactions, respectively. White dots highlight residues which are observed in alternate conformations. Blue stars indicate residues which have been shown to play a role in the OCP-PBS interaction, and red stars point to residues that could be at the origin of the stronger attachment of Plankto-OCP to the Syn-PBS. (B-C). The asymmetric unit of the *P*2_1_ crystals features a dimer (here shown as a ribbon with the two facing monomers colored sequence-wise, from cold (N-ter) to hot (C-ter) colors), whereas in *C*2 crystals, the dimer is crystallographic, hence only a single monomer is found in the asymmetric unit. (D) Polar contacts at the dimerization interface involve a conserved salt-bridge between D19 and R27, as well as conserved H-bonds between facing D6 and T90, and between facing N134 and N14, T15, and T17. (E, F, G) The *C*2 structures display a more compact conformation than the *P*2_1_ structures, at both the dimer (E, F) and the monomer levels (G). The figure illustrates the trajectory followed by Plankto-OCP Cα atoms as we interpolate from the *C*2-CAN to the *P*2_1_-CAN structure, highlighting the secondary structure elements which diverge most upon packing in the two crystal types. Arrows show the overall direction travelled by domains as we interpolate from the *C*2-CAN structure to the *P*2_1_-CAN Plankto-OCP chain A structure, revealing compaction of the OCP monomer.

In the *C*2 structures, the dimerization interface is crystallographic (-x, y, -z) and at the origin of the two-fold symmetry of the crystals. The BSA at the dimerization interface amounts to 1061 and 1023 Å^2^ in the Plankto-OCP_CAN_ and Plankto-OCP_ECN_ structures, respectively (Supplementary Table S1). We note that in the *C*2 structures, the relative contributions of αA and αB to the BSA at the dimerization interface hardly vary depending on the functionalizing carotenoid, amounting to respectively 32 and 40 % in Plankto-OCP_CAN_, and 30 and 42 %, in Plankto-OCP_ECN_ (Supplementary Table S1). Nonetheless, alike in the *P*2_1_ structures, the two H-bonds affixing αA and αB from facing monomers in the *C*2 Plankto-OCP_CAN_ dimer, *i.e.,* R9(NH2)-Q30(O) and R9(NH1)-L31(O), are absent in the *C*2 Plankto-OCP_ECN_ dimer. Thus, these H-bonds are present in both the *P*2_1_ and *C*2 Plankto-OCP_CAN_ structures, but in neither of the Plankto-OCP_ECN_ structures. The rest of the H-bonding network at the dimerization interface is yet overall preserved in the two *C*2 structures, with two to three H-bonds between αA and the facing αH (T15(O)-N134(ND2) and T17(OG1)-N134(ND2) in the two structures, and N14(OD1) to A133(N) in the Plankto-OCP_CAN_ structure only), one H-bond between αA and the facing αE-αF loop (D6(OD2)-T90(OG1) and a salt-bridge between facing αB residues (D19(OD2)-R27(NE)) (Figure 9D). The latter salt-bridge is preserved among all OCP structures reported thus far, suggesting that it is a defining interaction in the naturally-occurring OCP dimer. Indeed, it was shown that mutation into a leucine of the highly-conserved R27 yields a constitutively monomeric OCP [16]. The extent of the biological dimerization interface is accordingly overall preserved among all available OCP structures including ours, with a mean BSA of 1051 ± 90 Å^2^ (viz. 6pq1, 5ui2, 5hgr, 3mg1, 4xb5, and 7qd0, 7qcZ, 7qd1 and 7qd2). In this context, two structures stand out, viz. the *Tolypothrix* OCP_CAN_ and *Arthrospira* OCP_hECN_ structures, with BSA of 857.9 and 1186.1 Å^2^, respectively (Supplementary Tables S1 and S2).

Further analysis of the crystalline interfaces reveals that differences between the *P*2_1_ and *C*2 crystals originate at a second large interface, absent in previously characterized OCP structures (Figure 10). This additional interface, coined interface X, corresponds to the dimerization interface in the *P*2_1_ crystals (BSA of 1001.6 and 1054.6 Å^2^ in the Ntag-Plankto-OCP_CAN_ and Ntag-Plankto-OCP_ECN_ structures, respectively) but largely exceeds it in the *C*2 crystals (BSA of 1650.3 and 1548.7 Å^2^ in the Plankto-OCP_CAN_ and Plankto-OCP_ECN_ structures) (Supplementary Table S3). Interface X involves multiple secondary structure elements, including the αC-αD loop (∼ 20 and 36 % of the overall BSA in *C*2 and *P*2_1_ crystals, respectively), αD (∼ 20 and 10 %, respectively), αE (∼ 10 and 5 %, respectively), αG and the αG-αH loop (20 and 30 %, respectively), the linker (20 and 10 %), and the αM-β4 (∼ 5 and 4 %, respectively) and β5-β6 (∼ 2 and 5 %, respectively) loops (Supplementary Table S3). Only the first two structural elements contribute H-bonds at interface X in the *P*2_1_ crystals, whereas all of them do in the *C*2 crystals (Supplementary Table S3). Conformational changes in αG and in the αC-αD and αG-αH loops, and results in changes in the BSA contributed by the αC-αD loop (+8 and +24 % in the Ntag-Plankto-OCP_CAN_ and Ntag-Plankto-OCP_ECN_ structures, respectively), αD (-64 and -68 %, respectively), αE (-70 and -66 %, respectively) and the linker (-74 and -68 %, respectively), explaining the shrunken interface X in the *P*2_1_ crystals, compared to the *C*2 crystals (Supplementary Table S3). These changes in packing translate to changes in the opening-angles at the dimerization (Figure 9E, F, G) and NTD/CTD interfaces (Figure 11), offering a glimpse into the molecular breathing motions that animate OCP, at both the monomer and the dimer levels (Figure 10). Briefly, monomers come closer to one another in the *P*2_1_ (asymmetric unit) dimer than in the *C*2 (crystallographic) dimer (change in opening-angle and distance between chains: -5.6° and +1.1 Å for Ntag-Plankto-OCP_CAN_; - 6.8° and + 1.2 Å for Ntag-Plankto-OCP_ECN_).

**Figure 10:**
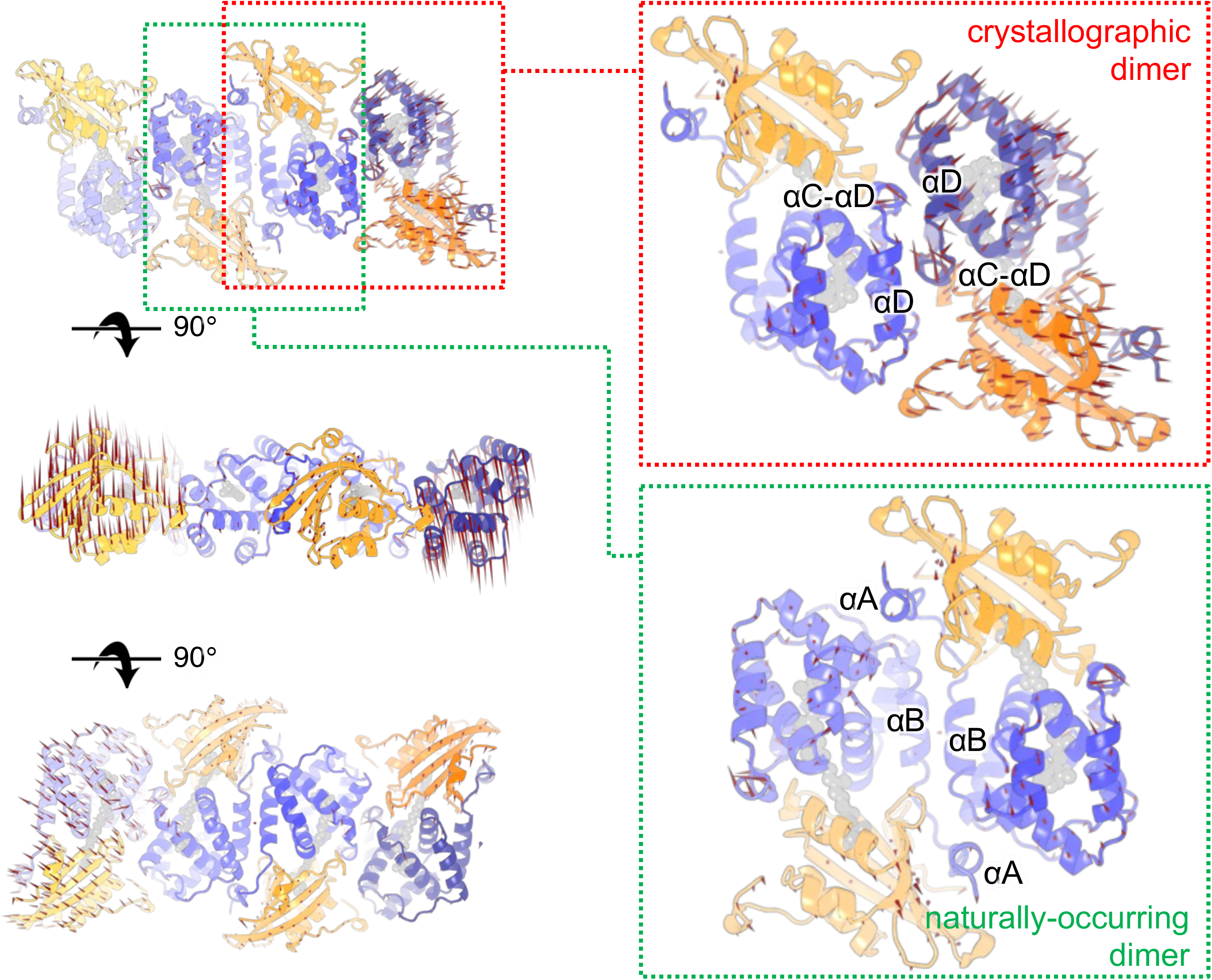
The *Planktothrix* OCP features two similarly large packing interfaces in crystals. In Plankto-OCP crystals, a large interface, additional to the dimerization interface (Table S1), is found which we coined interface X (Table S3). This interface, mainly contributed by helices αD, αE and αG, and by the αC-αD and αG-αH loops, matches the dimerization interface, in *P*2_1_ crystals (BSA of ∼1050 Å^2^), but largely exceeds it, in the *C*2 crystals (BSA of ∼1600 Å^2^). Changes in the extent of interface X result in a reorientation of domains in each monomer forming the naturally-occurring dimer, and in an increase in the opening angle between monomers, in the dimer. Arrows indicate the direction and distance along which Cα atoms travel as we interpolate from the *P*2_1_ to the *C*2 crystals.

**Figure 11:**
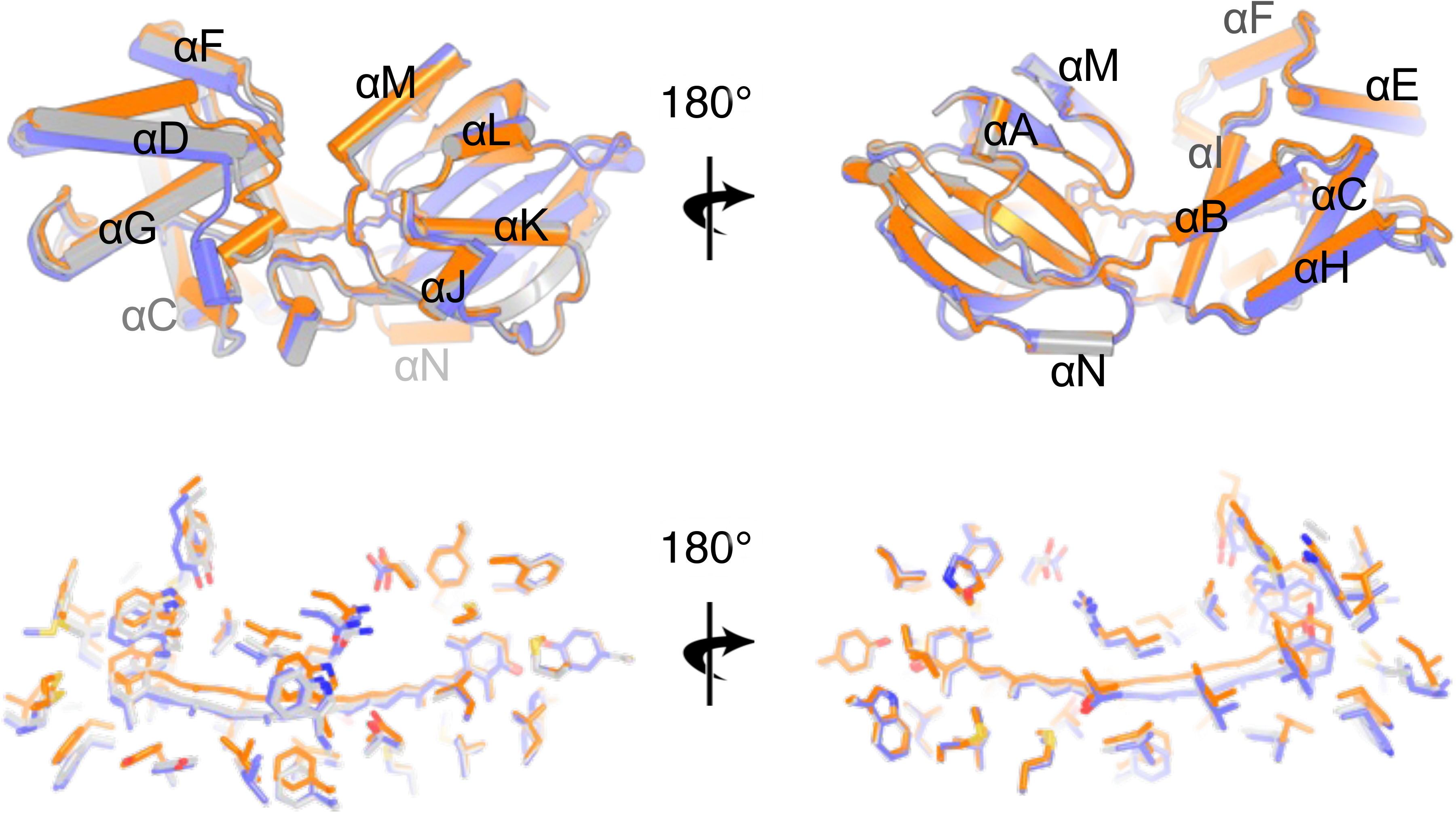
Plankto-OCP structures differ in the compaction of the monomer as well as in the internal structure of the NTD. (A) The *C*2 (orange), and *P*2_1_ chain A (slate) and chain B (grey) structures are overlaid as ribbons. Structural alignment, performed using the CTD atoms, highlight the change in opening angle between the CTD and the NTD. Indeed, the CTD structure is highly conserved with hardly no conformational changes observed amongst structures. Large differences are yet seen in the NTD, notably in the αC-αD and αG-αH loops, but as well in the relative positioning of the αC and αD helices. Here, the Plankto-OCP-CAN structures are shown, but the same observations can be made by comparing the Plankto-OCP-ECN structures. (B) Close up view on CAN and on the residues lining the carotenoid tunnel. A change in the orientation of the carotenoid is seen upon compaction of the OCP structure due to change in space group.

Yet, the *P*2_1_ monomers are characterized by an increased opening angle between the NTD and the CTD (Figures 9G, 10 and 11). Chain A shows a larger deviation with respect to the unique chain in *C*2 crystals (change in opening-angle and distance between the NTD and CTD: +3.1° and +1.0 Å for Ntag-Plankto-OCP_CAN_; +4.6° and +1.0 Å for Ntag-Plankto-OCP_ECN_) than chain B (change in opening-angle and distance between the NTD and CTD: +1.3° and +0.4 Å for Ntag-Plankto-OCP_CAN_; +1.4° and +0.3 Å for Ntag-Plankto-OCP_ECN_) (Figure 9). As a result, the predicted radii of gyration (Rg) for the *P*2_1_ asymmetric unit dimers (26.16 and 26.12 Å for Ntag-Plankto-OCP_CAN_ and Ntag-Plankto-OCP_ECN_, respectively) are larger than those predicted for the *C*2 crystallographic dimers (26.01 and 25.90 Å for Ntag-Plankto-OCP_CAN_ and Ntag-Plankto-OCP_ECN_, respectively) (Supplementary Table S2). Consistently, *P*2_1_ chain B features a structure that is intermediate between *P*2_1_ chain A and the unique C2 chain (change in opening-angle and distance between the NTD and CTD in *P*2_1_ chain A with respect to *P*2_1_ chain B: +3.3° and +0.9 Å for Ntag-Plankto-OCP_CAN_; +4.0° and +0.7 Å for Ntag-Plankto-OCP_ECN_) (Figure 11A). This is visible also in the predicted Rg for the various chains (20.81, 20.67 and 20.52 Å for Ntag-*P*2_1_ chain A, *P*2_1_ chain B and *C*2 Plankto-OCP_CAN_, respectively; and 20.73, 20.50 and 20.48 Å for the Ntag-Plankto-OCP_ECN_ counterparts, respectively) (Supplementary Table S2). Altogether, these changes affect the positioning of the carotenoid which tilts towards Y44 in the *P*2_1_ structures, despite preservation of H-bonds from its carbonyl oxygen to Y203(OH) (2.5-2.8 Å distance between non-hydrogen atoms) and W290(NH1) (2.8-3.0 Å distance between non-hydrogen atoms) and a quasi-perfect alignment of its β1-ring in the CTD (Figure 11B). With respect to the *C*2 Ntag-Plankto-OCP_CAN_ structure, the tilt angles of the carotenoid are 3.8 and 1.8° in chains A and B of the *P*2_1_ Ntag-Plankto-OCP_ECN_ structure, 3.2 and 2.9° in chains A and B of the *P*2_1_ Ntag-Plankto-OCP_CAN_ structure, and 0.8° in the *C*2 Ntag-Plankto-OCP_ECN_ structure. Thus, crystal packing traps different conformations of the Ntag-Plankto-OCP monomers, which differ in (i) the positioning of the carotenoid (Figures 9-12); (ii) the conformation displayed, at interface X, by αG and the αC-αD and αG-αH loops, (iii) the opening angle between domains, at the NTD/CTD interface; and (iv) in the opening angle between monomers, in the biological dimer. It is of important note that despite these changes, and the presence of two alternate side-chain conformations for R155 in the Ntag-Plankto-OCP_ECN_ structures, the R155-E246 salt-bridge and N104-W279 H-bond, which support the NTD / CTD interface, are preserved in all structures (distance between non-hydrogen atoms 2.8-3.2 Å) (Supplementary Figures 4 and 5). We also note that previously determined structures of *Synechocystis*, *Anabaena*, *Tolypothrix* and *Arthrospira* OCP align best with the *C*2 Ntag-Plankto-OCP_CAN_ and Ntag-Plankto-OCP_ECN_ structures, suggesting that these should be used for comparisons, rather than the *P*2_1_ structures (Supplementary Table S2). In this context, it must be stated that differences in the opening angle between the NTD and the CTD can also be seen from the comparison of the two chains constituting the asymmetric unit dimer in the *Anabaena* OCP structure (PDB id: 5hgr), with chain A aligning best with the *C*2 Ntag-Plankto-OCP conformers. We last note that ECN-functionalized OCP monomers and dimers are more compact than the CAN-functionalized counterparts (Figure 11 and Supplementary Table S2), and that a similar trend is visible in the comparison of the Ntag-Syn-OCP structures functionalized by CAN (PDB id: 4xb5; one chain with a predicted radius of gyration of 20.49 Å) and ECN (PDB id: 3mg1; two chains with predicted radii of gyration of 20.34 and 20.33 Å). In the case of the Plankto-OCP structure, conformational changes again concentrate in the NTD (Figure 12).

**Figure 12:**
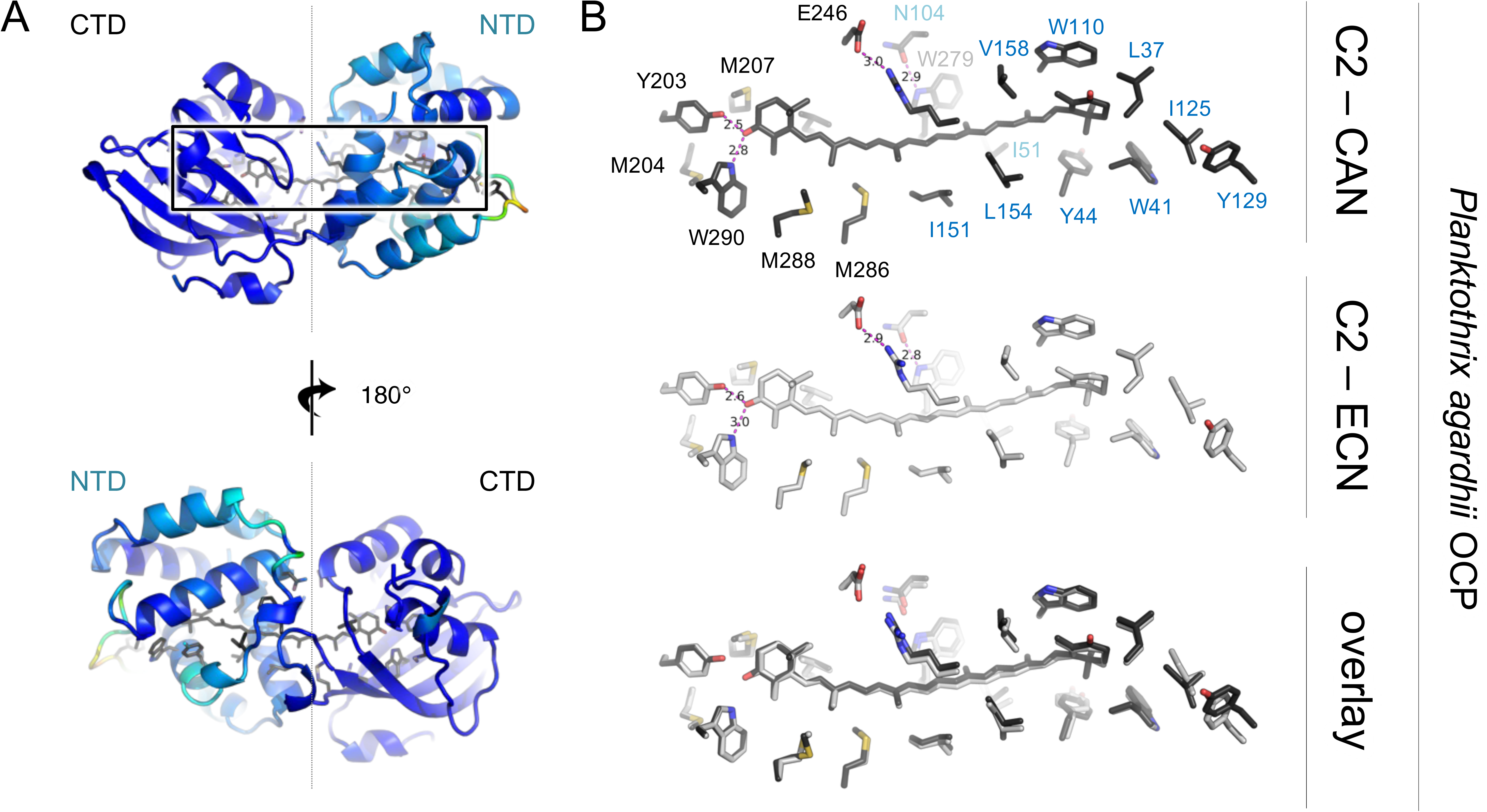
Conformational changes observed upon change of the functionalizing carotenoid are limited to the NTD. (A) The structure of Plankto-OCP-CAN is shown as a ribbon colored from cold to hot colors as a function of the RMSD to the Plankto-OCP-ECN structure. (B) Close-up view on the carotenoid and residues lining the homonymous tunnel. The orientation in (B) is similar to that in the upper panel of (A).

Conformational changes at the Cα level are best visualized in Cα-Cα distance difference matrices (DDM). Calculated using the *C*2 Ntag-Plankto-OCP_CAN_ structure as a reference, these confirm that the main difference between the *P*2_1_ chain A, *P*2_1_ chain B and *C*2 structures is a change in the orientation of the NTD vs. the CTD, accompanied (or triggered) by a change in the conformation of αA and the αC-αD and αG-αH loops. As a result, the two loops draw away from the CTD, pulling with them αC, αD, αE, αF and αG (Figure 13A). Conformational changes are far less pronounced in the CTD, where only β5 slightly changes position, moving away from αJ, β2 and αM (Figure 13A). In this context, it should be recalled that aside αA, the NTD consists of two 4-helix bundles contributed by helices αB-αC-αH-αI and αD-αE-αF-αG, respectively, which appose one onto the other leaving a central void that constitutes the carotenoid tunnel in the NTD. Our structural comparison indicates that regions most affected by the change in space group are those linking the two bundles, with conformational changes affecting the position – but not the internal structure – of the second bundle with respect to the CTD. Interestingly, the αC-αD and αG-αH loops are also the secondary structure elements most affected by the change in functionalizing carotenoid. From the comparison of the *C*2 structures, it is visible that these changes result in a modification of the distance between helices αD and αF, on the one hand, and the CTD, on the other hand. Thus, the structural dynamics at the basis of the change in space group and those resulting from the change in functionalizing carotenoid are localized in the same regions of the protein. This suggests that they could reflect a functional role. In line with this hypothesis, calculation of a DDM between the isolated NTD of Syn-OCP_CAN_, considered as a surrogate for the structure of the NTD in OCP^R^, and the NTD in the dark-adapted Syn-OCP_CAN_^O^ reveals major changes in the αC-αD and αG-αH loops (Figure 13B), which result in a dramatic rearrangement of the first αB-αC-αH-αI bundle whilst leaving unperturbed the internal conformation of the αD-αE-αF-αG bundle. Thus, like changes in space group and functionalizing carotenoid in Ntag-Plankto-OCP, photoactivation affects the internal structure and relative positioning of the Syn-OCP αB-αC-αH-αI bundle through conformational changes in the αC-αD and αG-αH loops, while the second αD-αE-αF-αG bundle appears stable and acts as a base. This observation is surprising given that the interface between the first bundle and the CTD is larger (BSA of 643.2 Å^2^; four H-bonds and a salt bridge) than that between the second bundle and the CTD (BSA of 189.3 Å^2^; 1 H-bond). Regardless, our results show that the two most mobile regions across the NTD – and therefore, across the whole Plankto-OCP – are the αC-αD and αG-αH loops, making these the first candidates to explain the increased photoactivation efficiency of Plankto-OCP.

**Figure 13:**
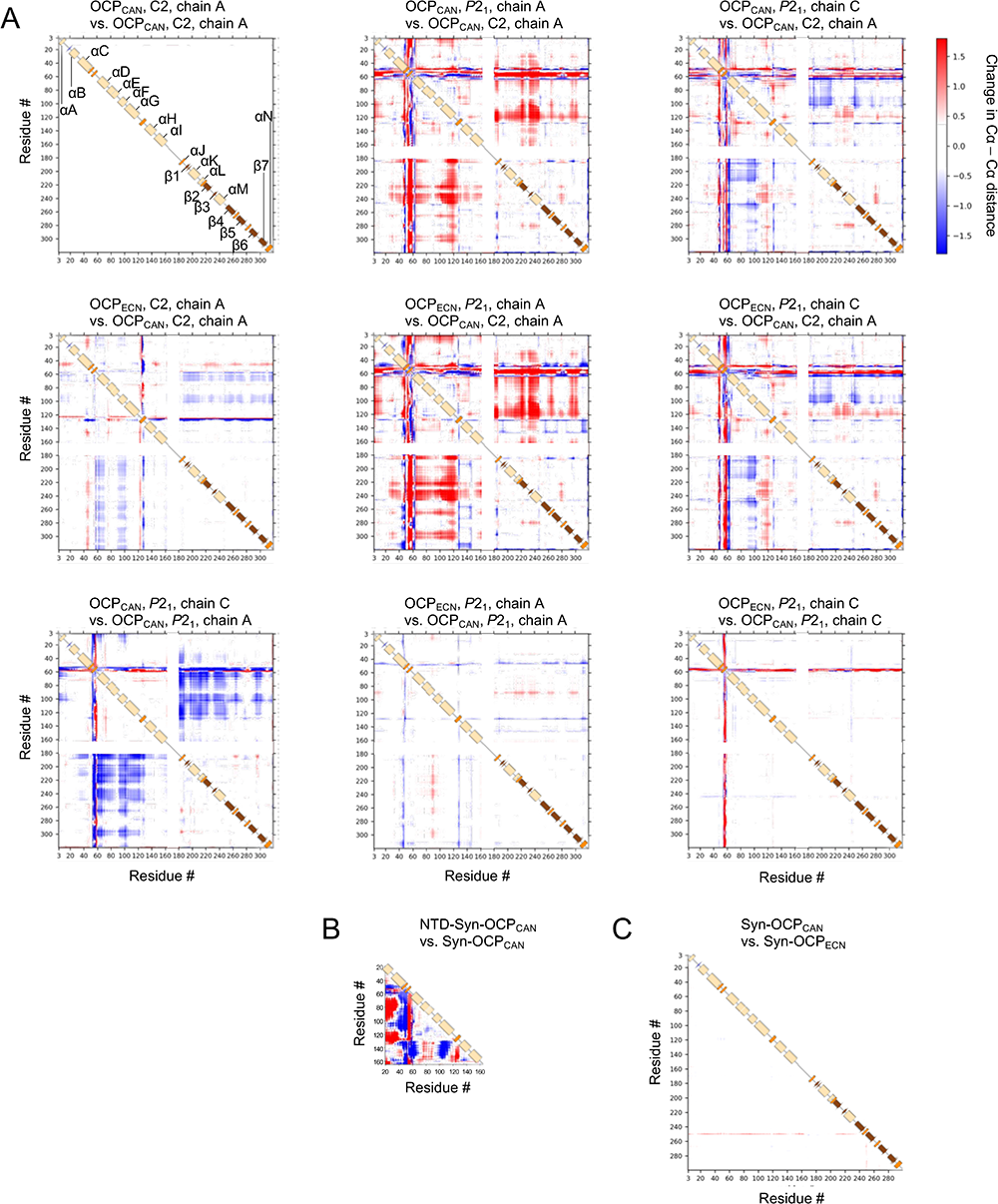
Crystal packing traps different conformations of the Ntag-Plankto-OCP monomers. (A) Changes in Cα-Cα distances in the various ECN- and CAN-functionalized *C*2 and *P*2_1_ chains were monitored by computing Cα-Cα difference-distance matrices (DDM). As a reference structure, we used the *C*2 Plankto-OCP-CAN (2 first rows), or either chain A or C of the *P*2_1_ PlanktoOCP-CAN structure (third line). In each DDM, the lower and upper panels (separated by a sketch of the secondary structure) show the changes in Cα-Cα distances for alternate conformers A and B with respect to the reference structure. Indeed, alternate conformations are seen in all Plankto-OCP structures. The overall similarity between upper and lower panels indicates that alternate conformations hardly affect the protein backbone. (Upper row) In the *P*2_1_ crystals, two chains are found, which either display a more expanded (chain A) or a more compact (chain B) structure, due to changes in the opening angle between the NTD and the CTD. The DDM further indicates that these changes stem from helix αD, and the αC-αD and αG-αH loops, either drawing away (chain A) or coming closer (chain B) to the CTD, respectively. (Middle row) Comparison of the *C*2-CAN and *C*2-ECN structures suggests that the presence of CAN results in a more compact protein, but support to this hypothesis could not be obtained from comparison of the *P*2_1_ chain A or *P*2_1_ chain B structures. (Lower row) Nonetheless, we observe a similar trend when comparing, either in the CAN-functionalized or the ECN-functionalized states, *P*2_1_ chain C and *P*2_1_ chain A. Thus, changes in functionalizing carotenoid have a lesser influence on the OCP conformation than changes in space group. (B) DDM calculated for the isolated NTD of Syn-OCP-CAN vs. Syn-OCP-CAN. This DDM suggests that upon photoactivation, large scale conformational changes occur in the NTD that mainly result in helix αB drawing away from helices αD to αG, while the αC-αD loop comes closer to these and to αH to αI. Also, αG edges closer to helices αH to αI, while the αG-αH loop draws farther. (C) DDM calculated for Syn-OCP_CAN_ vs. Syn-OCP_ECN_. Hardly no change in the Syn-OCP structure is seen upon change in the functionalizing carotenoid.

The αC-αD loop conformation differs in the *P*2_1_ chain A, *P*2_1_ chain B and *C*2 structures, but is overall preserved in each pair of ECN- and CAN-functionalized structures – notwithstanding the presence of multiple residues in alternate conformations in the C2 Ntag-Plankto-OCP_ECN_ structure (Figure 11 and Supplementary Table S2). The αG-αH loop conformation is overall preserved among *P*2_1_ structures, with only a slight displacement of αH observed between the *P*2_1_ chains A and B, as a result of changes in the αC conformation concomitant to those in the αC-αD loop (Figures 11 and 13). The αG-αH loop conformation is yet more divergent among the *C*2 structures, with the *C*2 Ntag-Plankto-OCP_ECN_ structure displaying two alternate main-chain conformations for the highly-conserved residues 122-VAPIPSGYKL-130, of which none overlaps with the conformations observed in *C*2 Ntag-Plankto-OCP_CAN_ and in the *P*2_1_ chains (Supplementary Figure 6 and Supplementary Table S2). Interestingly, these changes in the conformations of the αC-αD and αG-αH loops result in the opening/closing of water-channels from the protein surface to the carotenoid tunnel in the NTD. Briefly, four main channels (#1, #2, #3 and #4) can be identified on the basis of our structures (Supplementary Figures 7, 8 and 9). The first one (#1), which is colinear with the carotenoid axis and adjacent to the β2-ring, features at its bottleneck the fully conserved L37, M83, M117 and I125 (Supplementary Figures 7, 8 and 9) and encompasses the binding site of the carotenoid in the structure of the isolated NTD of Syn-OCP_CAN_ (Supplementary Tables S2 and S4). Channel #1 is opened in all Ntag-Plankto-OCP structures, and as well in all previously determined OCP structures. In the *C*2 Ntag-Plankto-OCP_ECN_ structure, however, the presence of two alternate main-chain conformations for αG-αH loop residues 120-GIAPIPSGYKL-131 (Supplementary Figure S6) results in two configurations of the channel (see Supplementary Table S4) characterized by either a wider or narrower opening (Supplementary Figures 6, 8 and 9). The enlargement of channel #1 is compulsory for the carotenoid to translocate fully across the NTD, as illustrated by the observation that the αG-αH loop displays a large conformational change in the structure of the isolated NTD of Syn-OCP_CAN_, compared to the NTD in the dark-adapted Syn-OCP^O^_CAN_ (Figure 13B) [21]. Hence the two *C*2 Ntag-Plankto-OCP_ECN_ conformers offer an illustration of how structural dynamics in the αG-αH loop may participate in the regulation of photoactivation. We yet note that none of the conformations observed in our various Ntag-Plankto-OCP structures, or in the previously determined OCP structures, matches that found in the structure of the isolated NTD of Syn-OCP_CAN_.

The second (#2), third (#3) and fourth channel (#4) are all perpendicular to the carotenoid tunnel, and open on either side of Y44 side chain, ending up just above the β2-ring of the carotenoid (#2) or its second terpene unit (#3, #4) (Supplementary Figures 7 and 8). Channel #2 features W41 and Y44 at its bottleneck, whereas channel #3 is lined by Y44, M47, I51, I151 and F280, and channel #4 by Y44, Y111 and αG (Supplementary Figures 7 and 8). Residues W41, M47, I151 and M280 are strictly conserved, whereas V53 and I53 fill the structural position of Plankto-OCP I51 in *Anabaena* OCP and Syn-OCP, respectively, and Y44 and Y111 are replaced by a phenylalanine (F44) and an asparagine (N111), respectively, in *Anabaena* OCP. We find that channel #2 is opened toward the bulk in all *P*2_1_ structures, albeit a much wider opening is observed in chain A of the *P*2_1_ Ntag-Plankto-OCP_ECN_ structure, due to a large swing of Y44 towards Y111 (Supplementary Figure 8) – *i.e.,* away from W41. Of note, Y44 is found in alternate side-chain conformations in the *C*2 Ntag-Plankto-OCP_CAN_ structure, and as well in the previously determined Syn-OCP_ECN_ structure (Supplementary Figure 8). Additionally, channel #3 is opened in chain A of the *P*2_1_ Ntag-Plankto-OCP_CAN_ and Ntag-Plankto-OCP_ECN_ structures, although in only one alternate conformer due to alternate conformations in M47. In both *C*2 structures, channel #2 and #3 are closed. Nonetheless, in one of the two alternate conformers of the C2 Ntag-Plankto-OCP_CAN_ structure, channel #4 is opened (Supplementary Figure 7). We note that all previously determined OCP structures feature similar openings of the carotenoid tunnel towards the bulk that are perpendicular to the carotenoid axis. For example, channels #3 and #4 are opened in the Syn-OCP structures (PDB ids: 3mg1 and 4xb5), while channels #2 and channel #3 are opened in the *Tolypothrix* OCP structure (PDB id: 6pq1) and in the two chains of the asymmetric unit dimer from the *Arthrospira* structure (PDB id: 5ui2; also referred to as *Limnospira*). In *Tolypothrix* OCP, a supplementary channel (#5) is observed, ∼ 180° apart from channel #2, which is lined by the highly conserved F28, I40, S157 and M161 (Supplementary Figure 10). The opening of channel #5, which alike channel #2 ends up above the β2-ring of the carotenoid albeit from the other side of the tunnel, mainly depends on the side chain conformation of M161 in the *Tolypothrix* OCP structure. Channel #5 is also opened in the two chains of the asymmetric unit dimer from the *Anabaena* OCP structure (PDB id: 5hgr), where its opening if facilitated by the substitution of I40 (in Plankto-OCP, Syn-OCP, *Arthrospira* and *Tolypothrix* OCP) for a valine – atop the conformational change in M161. Channel #3 is additionally opened in chain B of *Anabaena* OCP, but closed in chain A due to change in the conformation of M47. Thus, the Ntag-Plankto-OCP structures recapitulate most previously observed conformational dynamics in the carotenoid residues and illustrate that the opening of channel #1-4 mainly depends on the conformations displayed by residues in αC (A33-E46) and in the αC-αD (M47-G57) and αG-αH (G120-L131) loops. The *Tolypothrix* and *Anabaena* OCP structures additionally reveal the existence of another channel (#5), which ends up on the side of the carotenoid β2 ring where the methyl or carbonyl oxygen atom of ECN and CAN are exposed, respectively. Given that the *Tolypothrix* OCP features the same residue distribution as Plankto-OCP, Syn-OCP and *Arthrospira* OCP along the five channels, it is possible that channel #5 would open in these as well.

We noted earlier that the positioning of the carotenoid differs in the various Ntag-Plankto- OCP structures, due to changes in the opening angle between domains, *i.e.* at the NTD/CTD interface. Besides this point, the most notable dissimilarity in the carotenoid tunnel of Plankto-OCP is the presence of a methionine at position 207, instead of a leucine in other OCP (L205 in Syn-OCP, and L207 in *Anabaena, Arthrospira* and *Tolypothrix* OCP) (Supplementary Figures 5, 6, 10). At position 288, a methionine is found in Plankto-OCP, Ana-OCP (PDB id: 5hgr) and *Tolypothrix* (PDB id: 6pq1)) OCP (PDB id: 4xb5), which is substituted for an isoleucine in *Synechocystis* (I286) and *Arthrospira* OCP (I288) (Supplementary Figures 5, 6, 10). Thus, Plankto-OCP is particularly enriched in methionine residues. Of these two additional methionines, we speculate that only the former may be related to the higher photoactivation rate of Plankto-OCP, given that the slower *Tolypothrix* and *Anabaena* OCP also feature a methionine at position 288. The only other structural difference between the carotenoid tunnels of Plankto-OCP and Syn-OCP is the placement of Plankto-OCP I51 at the position occupied by Syn-OCP I53, due to changes in the sequence of the αC-αD loop (47-MGKTIT<u>V</u>AA<u>L</u>GAA-59 in Plankto-OCP vs. 47-MGKT<u>L</u>TIAA<u>P</u>GAA-59 in Syn-OCP). Again, we speculate that these changes could be involved in the higher photoactivation and recovery rates of Plankto-OCP, given that the αC-αD loop exhibits varying conformations in the various Ntag-Plankto-OCP structures (Supplementary Figures 10 and 12) but adopts the same conformation in the *Synechocystis* (Syn-OCP), *Anabaena*, *Arthrospira* and *Tolypothrix* OCP structures. Since the αC-αD loop in Ana-OCP (47-MGKTIT<u>V</u>AA<u>P</u>GAA-59) differs from that in Plankto-OCP only by the substitution of L56 for a proline – found as well in *Synechocystis* (Syn-OCP), *Arthrospira* and *Tolypothrix* OCP – it is most probable that the increased dynamics of this loop in Plankto-OCP originate from the replacement of this residue. P56 fits into a groove at the surface of the NTD (formed by S60, M61, G108, Y111 and W277 in Syn-OCP; S60, M61, G108, N111 and W279 in *Anabaena* OCP; and N60, M61, G108, Y111 and W279 in *Arthrospira* and *Tolypothrix* OCP) in all previously described OCP structures but is solvent exposed in Plankto-OCP (Supplementary Figure 10). This unfavorable feature could be at the basis of the increased conformational variability of the αC-αD loop in Plankto-OCP.

## Discussion

Here, we reported results from a comparative analysis of structure-function-dynamics relationships in various OCP1 from different cyanobacterial species, including the previously uncharacterized Plankto-OCP and the most studied Syn-OCP. All OCP-related functional properties – photoactivation, thermal recovery, interaction of the OCP^R^ state with the PBS and consequential fluorescence quenching and recovery – were examined for their dependence on the functionalizing carotenoid (ECN vs. CAN). We also investigated the influence of his-tagging at either the N or C-terminus, which afforded information on the role of the NTE (αA) and the CTT (αN) in the various molecular processes. In an attempt to rationalize functional observations, we solved the *hitherto* uncharacterized structures of Plankto-OCP in the ECN- and CAN-functionalized states, each in two space groups, and compared them to the available structures of ECN- and CAN-functionalized Syn-OCP [17,21]. When useful, we also included in our structural comparison the structures of CAN-functionalized *Anabaena* OCP [61], hECN-functionalized *Arthrospira* OCP [8] and of CAN-functionalized *Tolypothrix* OCP [30].

Initially, our interest in Plankto-OCP was sparked by the recent observation that the *in vivo* OCP-related NPQ-mechanism is more efficient in *Planktothrix* than in *Synechocystis* cells [41]. The functional characterization of Plankto-OCP properties afforded rationalization of this phenotype, showing that it not only photoactivates faster than Syn-OCP, but also recovers faster (especially at 9°C). Such a fast thermal recovery had thus far been observed only for OCP2 and OCPX variants, but not for members of the OCP1 clade. Indeed, *Synechocystis*, *Arthrospira* and *Tolypothrix* OCP1 are all characterized by a slow recovery at 9°C. The recovery of Plankto-OCP can be further accelerated by addition of FRP, confirming its belonging to the OCP1 clade, but the degree of acceleration is reduced by at least 8-fold. In previous studies, it was found that slow recovery and the ability to bind to FRP coincide in the OCP1 clade, as opposed to OCP2 and OCPX [16] which recover faster thermally, but are unable to bind FRP. Based on sequence alignments, it was further suggested that the defining feature of the OCP1 clade is the presence of residues R229 and D262 (*Synechocystis* number), which are absent in OCP2 and OCPX [16]. Our data confirm that presence of R229 and D262 correlates with the ability to bind FRP, but indicate that the slow recovery rate of *Synechocystis*, *Arthrospira* and *Tolypothrix* OCP1 is unrelated to their presence.

The faster photoactivation and recovery of Plankto-OCP could stem from its higher protein flexibility, illustrated by our Plankto-OCP crystalline structures. Indeed, crystal packing traps different conformations of the Ntag-Plankto-OCP monomers, which differ in (i) in the positioning of the carotenoid (Figures 9-13); (ii) the conformation displayed by αG and the αC-αD and αG-αH loops; (iii) the opening angle between domains at the NTD/CTD interface; and (iv) the opening angle between monomers in the naturally-occurring dark-adapted dimer. Hence, the Plankto-OCP structures offer a peek into the molecular breathing motions that animate OCP, at the monomer and the dimer levels. The *P*2_1_ structures demonstrate that the two monomers in a dimer can adopt slightly different structures, differing in the opening angle at the NTD/CTD interface, and consequently, in the orientation of the carotenoid in the tunnel. These differences could account for the spectroscopic observation that two states of dark-adapted OCP coexist *in vitro* [40]. The conformations present in the *C*2 structures match best those displayed by chain B in the *P*2_1_ structures but they are not equivalent, differing in the opening angle at the NTD/CTD interface, in the exact positioning of helices around the carotenoid, and in the conformation displayed by the αC-αD and αG-αH loops. These two loops show the highest diversity among the various structures reported herein, whereas they adopt the same conformation in all previously-determined *Synechocystis* (Syn-OCP), *Anabaena*, *Arthrospira* and *Tolypothrix* OCP structures. Hence, conformational diversity in these loops could be at the basis of the peculiar functional properties of Plankto-OCP. Below, we further detail how our observations could be linked to function.

The *P*2_1_ chain A and *P*2_1_ chain B structures feature a “porous wall”, traversed by channels perpendicular to the carotenoid tunnel, whereas the *C*2 structures feature a carotenoid tunnel that is insulated from the bulk, except at its extremities (Supplementary Figures 7, 8 and 9, and Supplementary Tables S2 and S4). Together, the six Plankto-OCP chains illustrate that the opening of channels #1-4 mainly depends on the conformations displayed by residues in αC (A33-E46; channels 1-2) and in the αC-αD (M47-G57; channels 2-4) and αG-αH (G120-L131; channel 1) loops, and question the possible role of bulk-water access to the carotenoid tunnel in the photoactivation mechanism. The enlargement of channel #1 is compulsory for the carotenoid to fully translocate across the NTD, and accordingly, a large conformational change was seen in the αG-αH loop in the structure of the isolated NTD of Syn-OCP_CAN_, compared to the NTD in the dark-adapted Syn-OCP^O^_CAN_ structure [21] (Figure 13). Two alternate conformations are observed for this loop in the *C*2 Ntag-Plankto-OCP_ECN_ structure, offering further illustration of how structural dynamics could participate in the regulation of photoactivation. We yet note that none of the conformations observed in our various Ntag-Plankto-OCP structures, nor in any of the previously determined OCP structures, matches that found in the structure of the isolated NTD of Syn-OCP_CAN_ [21], indicating that full translocation of the carotenoid into the NTD must occur for this conformation of the αG-αH loop to become favorable.

We also investigated if the presence of multiple alternate conformations could underlie the higher photoactivity of Plankto-OCP. Plotting these against the secondary (Figure 9A and Supplementary Table S2) or tertiary structure, we identify the αC-αD (M47-G57) loop, αM and β4 as clusters. Indeed, these secondary-structure elements or epitopes appear prone to display alternate conformations, irrespective of the space group or functionalizing carotenoid (Figure 9A). For example, W288, at the CTD end of the carotenoid tunnel, is present in alternate conformations in all structures. Additionally, D6, N14, Q73, Q77, M117, M207, R241, R244, V258, K275 are present in alternate conformations in three of the six structurally-analyzed Plankto-OCP chains. In conclusion, we propose that the structural traits which explain the higher photoactivation rate of Plankto-OCP are (i) the increased flexibility in the αC-αD (M47-G57) and αG-αH (G120-L131) loops, which results in opening of water channels to the carotenoid tunnel; and (ii) the increased molecular breathing motions at the levels of the dimer, the NTD/CTD interface, and the NTD helix core. As to the increased recovery rate, a possible hypothesis is that it originates from a decreased stabilization of the Plankto-OCP^R^ state. Comparing residues distribution along the carotenoid tunnel in the NTD, we found that the only structural difference between the NTD tunnels of Plankto-OCP and Syn-OCP is the placement of Plankto-OCP I51 at the position occupied by Syn-OCP I53, due to changes in the sequence of the αC-αD loop (47-MGKTIT<u>V</u>AA<u>L</u>GAA-59 in Plankto-OCP *vs.* 47-MGKT<u>L</u>TIAA<u>P</u>GAA-59 in Syn-OCP) (Supplementary Figures 5, 6 and 10), suggesting a role for this residue in the stabilization of OCP^R^. Another, more hypothetical means by which the Plankto-OCP^R^ state would be rendered less stable could be through a decrease in the affinity of the CTT for the empty CTD tunnel. Syn-OCP is indeed unique among the variants discussed herein in that it features a 314-NFAR-317 sequence at its C-terminus instead of 316-NLVR-319 in the other OCP (Supplementary Figure 10). The combined substitution of a leucine for a phenylalanine, and of a valine for an alanine, could account for a higher affinity of the Syn-OCP CTT for the empty CTD tunnel. Reversely, the replacement of Syn-OCP I286 by a methionine in Plankto-OCP (M288) could decrease the affinity of the CTT for the empty CTD tunnel.

We also examined the effect of his-tagging at the N- or C-terminus on the various functional properties of OCP (Figures 3 and 8). Indeed, both the NTE and the CTT have been proposed to play important roles in the OCP photoactivation and recovery mechanisms. We found that irrespective of the variant, the N-tagged protein displays the best match to the native protein, both in terms of photoactivation rate and thermal recovery rate. Native Syn-OCP and Plankto-OCP display the fastest photoactivation rate, indicating that increased disorder in their NTE and/or a weaker interaction between the NTE and the CTD does not benefit photoactivation as was previously suggested [64]. Recovery was also found to be faster in native Syn-OCP and Plankto-OCP than in their his-tagged counterparts. Notably, a dramatic drop in the recovery rate was observed for the C-tagged Plankto-OCP, which could stem either from a higher stability of the OCP^R^ state in presence of a C-terminal his-tag, or from a frustrated recovery of the OCP^O^ state due to hindered rebinding of the CTT to the CTD β-sheet. Regardless, these results point to an important role played by the CTT in the stabilization of OCP^R^ and thermal recovery of the OCP^O^ state. Little is known regarding the configuration of the CTT residues in the OCP^R^ state and its role in OCP photoactivation and recovery. However, it was shown that in the *Synechocystis* C-terminal domain homologue (CTDH), which belongs to a family of carotenoid-transporting proteins structurally homologous to the OCP CTD [36,37], the CTT can exchange between a closed conformation, whereby it covers the empty carotenoid tunnel, and an opened conformation, similar to that observed in OCP^O^. By analogy, it was therefore proposed that the last step of OCP photoactivation is the repositioning of the CTT on the empty CTD-half of the carotenoid tunnel, preventing exposure to the bulk of its highly hydrophobic residues. Presence of a six-histidine tag in the CTT could hinder this movement, leading to slower kinetics of photoactivation and recovery. The position of the his-tag also influences the PBS quenching, with again a more pronounced effect of C-terminal his-tagging. It was earlier proposed that the presence of a six-histidine tag upstream the NTE could destabilize the interaction with the PBS [64], yet the present data invalidate the hypothesis, demonstrating that N-tagged and native OCP detach as swiftly from the PBS. Contrastingly, introduction of the six-histidine tag downstream the CTT results in increased PBS-fluorescence quenching and slower fluorescence recovery. This result can be rationalized by envisioning that the C-terminal his-tag strongly stabilizes the PBS-bound OCP^R^ structure, possibly through interaction of the tag with PBS amino acids. In conclusion, our investigation of the effect of his-tagging on the functional properties of OCP suggest that rebinding of the CTT and (to a lesser extent) the NTE to the CTD β-sheet are rate limiting steps in the thermal recovery of the OCP^O^ state. Completion of these steps is inhibited by presence of a his-tag at either the C- or N-terminal extremities but C-terminal his-tagging furthermore stabilizes the OCP^R^/PBS complex.

A cross-species characterization of OCP/PBS complexes was also conducted, which revealed that irrespective of his-tagging, Plankto-OCP binds stronger to Syn-PBS than does Syn-OCP, and binds stronger to Syn-PBS than to Plankto-PBS. These findings echo the previous observation that both *Arthrospira* and *Anabaena* OCP bind stronger to Syn-PBS than to *Arthrospira* and *Anabaena* PBS, respectively [43]. Recent studies on Syn-OCP have shown that besides R155, early demonstrated as compulsory for binding to the PBS [65], residues L51, P56, G57, A58, N104, I151, and N156 play important roles in the OCP/PBS interaction [66]. In all OCP, N104 and R155 are involved in the stabilization of the OCP^O^ state by contributing H-bonds to the NTD/CTD interface (N104(OD1) to W277(Syn)/W279 (Plankto)(NE1); R155(NH2) to E244 (Syn)/E246 (Plankto)(OE1)) and in addition, N104 contributes to the stabilization of the linker by establishing a H-bond to E174(OE1) in all structures but the Plankto-OCP structures. As G57, N104, I151, R155 and N156 are structurally conserved in Plankto-OCP, one can eliminate the hypothesis that the observed differences in Syn-PBS quenching would stem from these. As to Syn-OCP L51, it is replaced by an isoleucine in the *Anabaena*, *Limnospira* and *Tolypothrix* proteins, but the side chain of this residue occupies the same position in all structures, fitting in a groove contributed by M47, I151, F280 and V284, at the NTD-CTD interface. In Plankto-OCP, L51 is conserved but it is found at the position occupied by I53 in all previously-determined OCP structures due to changes in the sequence of the αC-αD loop (47-MGKTIT<u>V</u>AA<u>L</u>GAA-59 in Plankto-OCP vs. 47-MGKT<u>L</u>TIAA<u>P</u>GAA-59 in Syn-OCP). Contrastingly Syn-OCP P56 is replaced by a leucine in Plankto-OCP, while being conserved in other OCP. Hence, the difference in PBS binding-affinity and fluorescence-quenching observed between Syn-OCP and Plankto-OCP could stem from this specific change in the sequence and structure of the αC-αD loop. Within 4 Å of R155, supposedly central to the interaction, we find four additional candidate positions at which residue substitutions could explain the higher affinity of Plankto-OCP as compared to Syn-OCP: G99, T102, A103 and C157, respectively. These residues are substituted by A99, S102, P103 and A157 in Syn-OCP, and by A99, S102, P103 and S157 in Ana-OCP. Future mutagenesis work concentrated on these residues could unveil the molecular basis for this unexpected cross-species preference of all tested OCP for the Syn-PBS. Specific to the lower affinity of Plankto-OCP for Plankto-PBS than for Syn-PBS, it must be recalled that OCP binding to the PBS is very sensitive to the structural intactness of the PBS core. Although 77K fluorescence spectra of both isolated PBS suggested an as efficient energy transfer from PC to the last core emitters, we cannot discard the hypothesis that a slightly different interaction between the APC trimers could be at the origin of the weaker OCP-PBS binding observed for *Arthrospira* and *Planktothrix* species-specific complexes. As it was also observed that the Plankto-Lcm protein is more sensitive to proteolysis than Syn-Lcm (Supplementary Figure 3), another (not necessarily exclusive) hypothesis could be that the reduced affinity of Plankto-OCP for the Plankto-PBS originates in the partial degradation of the Lcm component.

We last examined the influence of the functionalizing carotenoid on OCP excited and intermediate states dynamics, following the observation that CAN-functionalized OCP photoactivates faster and recovers slower than ECN-functionalized OCP. Using fs-ns and ns-s transient absorption spectroscopy on ECN- and CAN-functionalized Plankto-OCP and Syn-OCP, we inquired the time scale(s) on which the gain in photoactivation efficiency occurs for the CAN-functionalized proteins. We observed differences in the respective yields of the S_1_, ICT and S* states, but nearly no change in their characteristic lifetimes nor in the P_1_ yield (Table 1). Hence, the difference in photoactivation rate of ECN- and CAN-functionalized OCP does not stem from changes in their excited state dynamics nor in the P1 formation quantum yield. Interestingly the increased S* yield observed for CAN-functionalized Plankto-OCP and Syn-OCP is not mirrored by an increase in the P_1_ yield, suggesting that the former is not its only precursor. Contrastingly, clear differences between the four tested OCP were observed in the ns-s time scale. The most striking differences between Plankto-OCP and Syn-OCP are visible in the ns-ms time window, while those between CAN- and ECN-functionalized OCP concentrate in the ms-s timescale. Thus, our results suggest that both carotenoid translocation (ns-µs) and NTE/CTT detachments (µs-ms) are affected by the change in protein scaffold, whereas it is domain dissociation that is most affected by a change in the functionalizing carotenoid (ms-s). Our data support that domain dissociation is faster and more efficient in CAN-functionalized OCP, with no decline observed in the difference absorption at 550 nm. Specific to Plankto-OCP_CAN_, a faster domain separation is observed with virtually no recovery to OCP^O^ on the µs-s timescale (100% efficiency from P_3_ to final OCP^R^). The increased photoactivation rate of Plankto-CAN is likely grounded in this property. Thus, the differences observed in the photoactivation speed of CAN- and ECN-functionalized Plankto-OCP and Syn-OCP stem from changes in the (comparatively-slow) carotenoid translocation, NTE/CTT detachment and domain dissociation steps – rather than from not changes in the excited-state dynamics or P1 formation quantum yield. We yet must note that a larger drop in difference absorption at 563 nm is seen on the ns-µs time scale, which could sign for a higher barrier for carotenoid translocation into the Plankto-NTD than the Syn-NTD.

It remains unclear whether or not canthaxanthin is used as a functionalizing carotenoid for OCP in the natural context. When expressed in their parent strain, *Arthrospira* and *Synechocystis* OCP [8,11,17] bind hECN, while *Tolypothrix* OCP1 binds CAN [30]. Recombinant expression of *Arthrospira* and *Anabaena* OCP in *Synechocystis* cells also yield an hECN-functionalized proteins [33,43]. Yet, when overexpressed and isolated from CAN-producing *E. coli* cells, all these OCP bind CAN, suggesting that OCP may alternatively bind hECN or CAN depending on the carotenoid presents in the cells. We earlier reported the partial inability of *Anabaena* and *Tolypothrix* OCP to fully convert to OCP^R^, when functionalized by ECN [8,11,17]. Here, thermal recovery kinetics were found to be slower in the CAN-functionalized Syn- and Plankto-OCP, possibly due to a reduced stabilization of the OCP^R^ state by ECN, as compared to CAN. Assuming that the β-rings of the carotenoid are exposed to the bulk in the OCP^R^ state, as suggested by the structure of the CAN-functionalized isolated NTD of Syn-OCP (PDB id: 4xb4; [21]), the higher stability of the CAN-functionalized OCP^R^ could result from the presence of an oxygen on its β2 ring, favoring interaction with the bulk. A similar stability would thus be expected for hECN, which features a hydroxyl group in the β2-ring. Hence, the possibility remains that all OCP bind hECN in their parent strain.

Supporting this hypothesis is the observation that quenching by *Plankto*-OCP of the *Plankto-*PBS fluorescence is efficient only when ECN is used as the functionalizing carotenoid (or when the protein is tagged at the C-terminus). It is difficult to rationalize the observation that CAN-OCP interaction with the PBS is weaker than that with ECN-OCP (Figures 7 and 8, and Supplementary Figure 11). These results are at variance with the observation that CAN stabilizes the OCP^R^ state, and suggest that the isolated OCP^R^ and PBS-bound-OCP^R^ structures could differ. The sole difference between CAN- and ECN-functionalized OCP^R^ is the presence of a carbonyl oxygen on the β2-ring, but this difference should not affect binding to the PBS, since its epitope has been mapped at the opposite end of the carotenoid tunnel in the NTD, proximate to R155. Thus, it is presumably the β1-ring, identical in CAN and ECN, which will be in contact with the PBS. A possible explanation could be that upon OCP binding to PBS, the carotenoid migrates into to the PBS, enabling a better interaction with the bilin pigments, as proposed earlier [21,67]. If it is the β1-ring that plunges into the PBS – *i.e.,* the carotenoid moves backwards with respect to the OCP^O^ to OCP^R^ transition – then the β2-ring will be repositioned inside the highly hydrophobic carotenoid tunnel, possibly past its original position in the OCP^O^ structure, which would explain the reduced stability of the CAN-functionalized OCP-PBS complexes, due to replacement of a methyl in ECN by a carbonyl oxygen in CAN. Thus, our results support the hypothesis that the isolated and PBS-bound OCP^R^ differ. If true, OCP-related quenching of PBS-fluorescence would be a two-step reaction (at the very least), with first the binding of OCP^R^ to PBS, and then a change in the OCP^R^ structure – as initially proposed by [21,67].

## Conclusion

We here have reported on a comparative structure-function-dynamics study on two OCP from the OCP1 clade, *i.e.,* Syn-OCP and Plankto-OCP. Our structures reveal that Plankto-OCP is more flexible and we speculate that this increase in flexibility explains its faster photoactivation and recovery. Specifically, our Plankto-OCP structures evidence increased structural dynamics in the αC-αD loop, shown to play a central role in the interaction between OCP^R^ and PBS [65–67]. Increased dynamics in this loop could be at the origin of the stronger binding of Plankto-OCP (compared to Syn-OCP) to Syn-PBS. Irrespectively, our data point to more efficient carotenoid translocation and NTE/CTT detachments in Plankto-OCP. We also show that presence of a his-tag influences both the photocycle of OCP [32] and its interaction with the PBS. Most impacting is introduction of the his-tag at the C-terminus, which results both in large damping of the photoactivation and recovery and in stronger binding to the PBS, suggesting an important role for the CTT in these molecular processes. Our work last uncovers the strong influence of the nature of the functionalizing carotenoid on all aspects of OCP function. The mere substitution on ring β2 of a dimethyl in ECN by a carbonyl in CAN results in increased photoactivation efficiency, which our ns-s spectroscopic data suggest to be due to faster domain dissociation. Nonetheless, CAN-functionalized OCP display reduced recovery, reduced binding to PBS and reduced energy-quenching activity. These features could result from thwarted back-migration of CAN, compared to ECN, as required for recovery and, presumably, PBS quenching and stabilization of the OCP-PBS complex. Additionally, the observation that quenching of Plankto-PBS is efficient only when by Plankto-OCP is functionalized by ECN suggests that likely, the carotenoid functionalizing OCP in the *Planktothrix* cells is not CAN.

## Supporting information

All Supplementary Figures and Tables

## Acknowledgments

We are grateful to Giorgio Schirò and Martin Weik for continued support of the project. We thank Sandrine Cot for technical assistance. We thank the ESRF and SLS synchrotron radiation facilities for beamtime allocation under long-term projects MX1992 and MX2329 (IBS BAG at ESRF), and acknowledge financial support by CEA, CNRS, Université Grenoble Alpes and Université Paris-Saclay. IBS acknowledges integration into the Interdisciplinary Research Institute of Grenoble (IRIG, CEA). This work was supported by the Agence Nationale de la Recherche (grants ANR-17-CE11-0018-01 to J.-P.C. and ANR-2018-CE11-0005-02 to the three French laboratories), the Polish National Science Centre (NCN project 2018/31/N/ST4/03983), and used the platforms of the Grenoble Instruct-ERIC center (ISBG; UMS 3518 CNRS-CEA-UGA-EMBL) within the Grenoble Partnership for Structural Biology (PSB). Platform access was supported by FRISBI (ANR-10-INBS-05-02) and GRAL, a project of the University Grenoble Alpes graduate school (Écoles Universitaires de Recherche) CBH-EUR-GS (ANR-17-EURE-0003).

## Data and materials availability

Atomic coordinates and structure factors have been deposited in the Protein Data Bank under the following accession codes: 7qd0, 7qcZ, 7qd1 and 7qd2. All other data are available in the main text or the supplementary materials.

## Supplementary figure captions

**Supplementary Figure 1: OCP absorption spectrum changes upon photoactivation.** (A) Absorbance spectra of CAN-functionalized Syn-OCP (black) and Plankto-OCP (red) in the dark-adapted (OCP^O^, continuous line) and light-adapted (OCP^R^, dashed line). (B) Difference-absorbance spectra (light-adapted minus dark-adapted) derived from (A).

**Supplementary Figure S2: Femtosecond transient absorption data collected on the Ntag-Syn-OCP.** (A, B) Transient absorption spectra measured after excitation at 532 nm are shown for time delays ranging between 0.14 and 1 ns for (A) Ntag-Syn-OCP_ECN_ and (B) Ntag-Syn-OCP_CAN_. All datasets were normalized to -1 at the bleaching minimum (∼500 nm), in both spectral and temporal dimensions. (C, D) Decay Associated Spectra (DAS) obtained from the global fit of the transient absorption data spectra shown in (A) and (B), respectively. Data were fitted using four exponential components convolved by a Gaussian pulse (IRF, 110 fs FWHM) and an offset attributed to P_1_ (lifetime > 10 ns).

**Supplementary Figure 3: *Planktothrix* and *Synechocystis* phycobilisomes.** (A) SDA-PAGE of the proteins present in *Synechocystis* (Syn) and *Planktothrix* (Plankto) phycobilisomes. Both the Syn-PBS and the Plankto-PBS are characterized by a similar phycocyanin (PC) to allophycocyanin (APC) ratio of ∼3 (compared to ∼2 in *Arthrospira* PBS) and feature a comparable protein composition, with an Lcm (ApcE) of about 95 kDa, two rod linkers (L_R33_ and L_R30_) and a rod-core linker (L_RC_). Nevertheless, the principal Lcm band in Plankto-PBS was found to migrate at a lower molecular weight than that of Syn-PBS, and was accompanied by a second band at 75 kDa, suggesting partial proteolytic degradation (B) Absorbance spectra of purified Plankto-PBS (fuchsia), Syn-PBS (black) and *Arthrospira*-PBS (blue). The absorption spectrum of Syn-PBS features a sharp band characterized by a maximum at 620 nm and a shoulder at 650 nm. A broader band with a maximum at 615-618 nm and no 650 nm shoulder is seen for Plankto-PBS and *Arthrospira*-PBS (Jallet et al., 2014). In the case of *Arthrospira*-PBS, the observation was rationalized by the presence of phycobiliviolin instead of phycocyanobilin in some phycocyanin (PC) units of the PBS (Babu et al, 1991). This could also be the case for Plankto-PBS. (C) Florescence spectra at 77K of purified Plankto-PBS (fuchsia) and Syn-PBS (black). The intense peak at 685 nm (last acceptor of PBS) and very low peaks at 645-660 nm (PC and APC) indicate that the energy-transfer from the PC to the final emitters (ApcE and ApcD) is as efficient in Plankto-PBS as it is in Syn-PBS.

**Supplementary Figure 4: Overview of the carotenoid and of residues lining its binding tunnel in the CAN-functionalized Plankto-OCP and Syn-OCP structures.** Residues that differ in the two proteins are labelled in red. H-bonds and salt-bridges are highlighted by pink dashed lines.

**Supplementary Figure 5: Overview of the carotenoid and of residues lining its binding tunnel in the Plankto-OCP-ECN and Syn-OCP-ECN structures.** Residues that differ in the two proteins are labelled in red. H-bonds and salt-bridges are highlighted by pink dashed lines.

**Supplementary Figure 6: Overview of channels affording bulk-solvent access to the carotenoid tunnel in the CAN-functionalized Plankto-OCP and Syn-OCP structures.** Channels were identified using Caver 3.0 and are localized in the NTD. Channel #1 is at the NTD end of the carotenoid tunnel, whereas channels #2, #3 and #4 are perpendicular to the later.

**Supplementary Figure 7: Overview of channels affording bulk-solvent access to the carotenoid tunnel in the ECN-functionalized Plankto-OCP and Syn-OCP structures.** Channels were identified using Caver 3.0 and are localized in the NTD. Channel #1 is at the NTD end of the carotenoid tunnel, whereas channels #2, #3 and #4 are perpendicular to the later.

**Supplementary Figure 8: Distribution of interatomic distances at the bottlenecks of channels #1 to #4 in the *Plankthotrix*, *Synechocystis*, *Limnospira*, *Anabaena* and *Toplipothrix* OCP structures**. (A) Box plot of the distribution of absolute distances. (B) Box plot of the distribution of distances relative to the average amongst the various investigated structures. In (A), structures showing the largest deviation are highlighted.

**Supplementary Figure 9: Sequence and secondary structure alignment of the five OCP variants compared structurally.** (A) β-sheets are shown in marine blue, while canonical (3.6_13_-helix) and 3_10_-helices are shown in red and yellow, respectively. Starbursts indicates residues whose substitution might play a role in the increase photoactivation rate (green) of Plankto-OCP. (B) The phylogenetic tree computed from the sequence-alignement of the five OCP variants reveals that alike *Plankthotrix and Tolypothrix* OCP belongs to a different sub-clade than *Synechocystis*, *Limnospira* and Anabaena OCP.

**Supplementary Figure 10: Close-up view on the FRP and PBS epitopes in the Plankto-OCP and Syn-OCP structures.** The FRP epitope, featuring residues that interact with helix αA in the dark-adapted OCP structures, is very well conserved amongst Plankto-OCP structures. When compared to Syn-OCP, however, slight changes can be seen in the distribution of charged residues at this interface, which could explain the reduced acceleration of Plankto-OCP recovery by FRP. The PBS epitope, featuring NTD residues facing the CTD and within 4 Å of R155 in the dark-adapted OCP structure, is less conserved amongst Plankto-OCP structures, due to increased structural dynamics in helix αC and in the αC-αD loop. The *C*2 Plankto-OCP structure is that most similar to Syn-OCP, in terms of charged-residues distribution at this interface. The functionalizing carotenoid does not seem to have an influence on the structuring of the PBS and FRP epitopes.

**Supplementary Table 1: Recapitulation of the contribution of secondary structure elements to the dimerization interface in terms of buried surface area (Å^2^) and number of H-bonds.**

**Supplementary Table 2: Characteristics of the main crystal packing interfaces, of the carotenoid tunnel, and predicted radii of gyration and number of alternate conformations in the various OCP structures, including ours.**

**Supplementary Table 3: Recapitulation of the contribution of secondary structure elements to interface X in terms of buried surface area (Å^2^).**

**Supplementary Table 4: Distribution of interatomic distances at the bottlenecks of channels #1 to #4 in the *Plankthotrix*, *Synechocystis*, *Limnospira*, *Anabaena* and *Tolypothrix* OCP structures.**

