## Supplementary material for "Structure-function-dynamics relationships in the peculiar *Planktothrix* PCC7805 OCP1: impact of his-tagging and carotenoid type": All Supplementary Figures and Tables

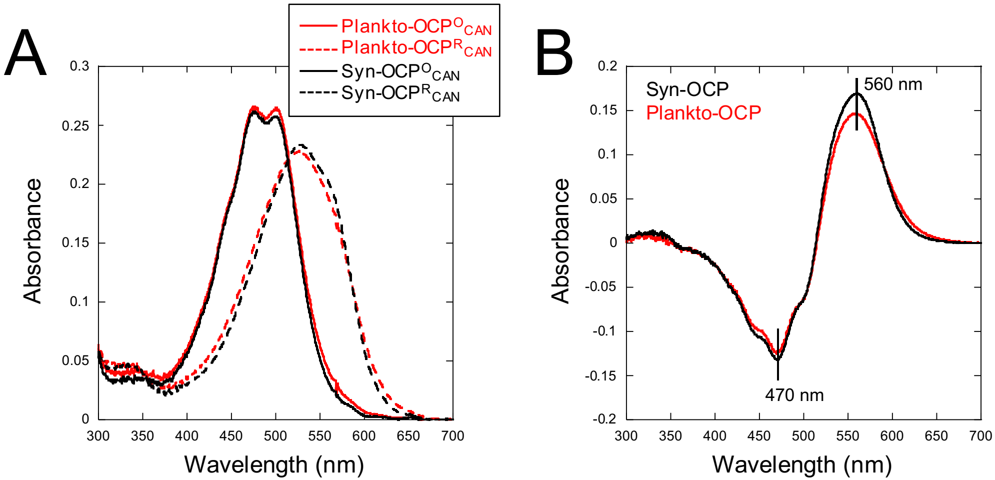
A
B
Plankto-OCPO CAN
Plankto-OCPR CAN
Syn-OCPO CAN
Syn-OCPR CAN
Absorbance
Wavelength (nm)
Syn-OCP
Plankto-OCP
560 nm
470 nm
0.3
0.25
0.2
0.15
0.1
0.05
0
-0.05
-0.1
-0.15
-0.2
300
350
400
450
500
550
600
650
700

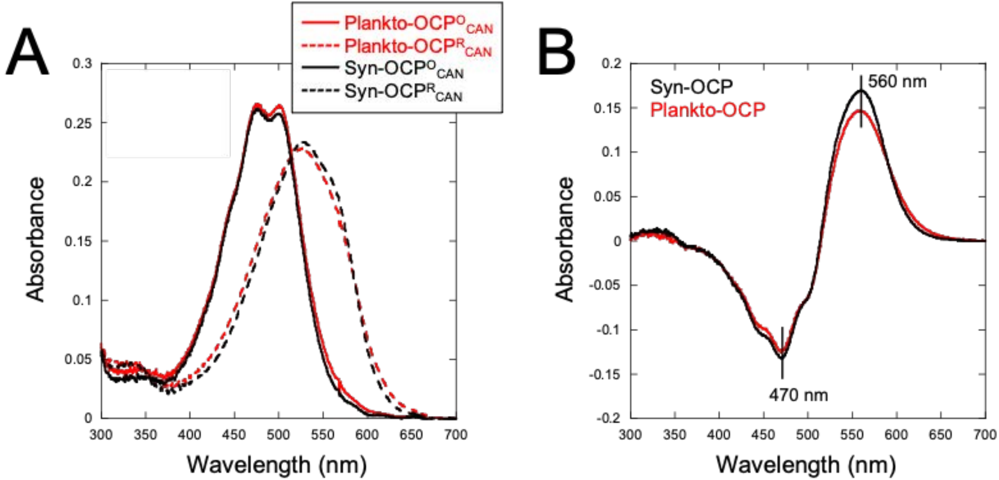
A
Plankto-OCP$^O_{CAN}$
Plankto-OCP$^R_{CAN}$
Syn-OCP$^O_{CAN}$
Syn-OCP$^R_{CAN}$
Absorbance
Wavelength (nm)
B
Syn-OCP
Plankto-OCP
560 nm
470 nm
Absorbance
Wavelength (nm)

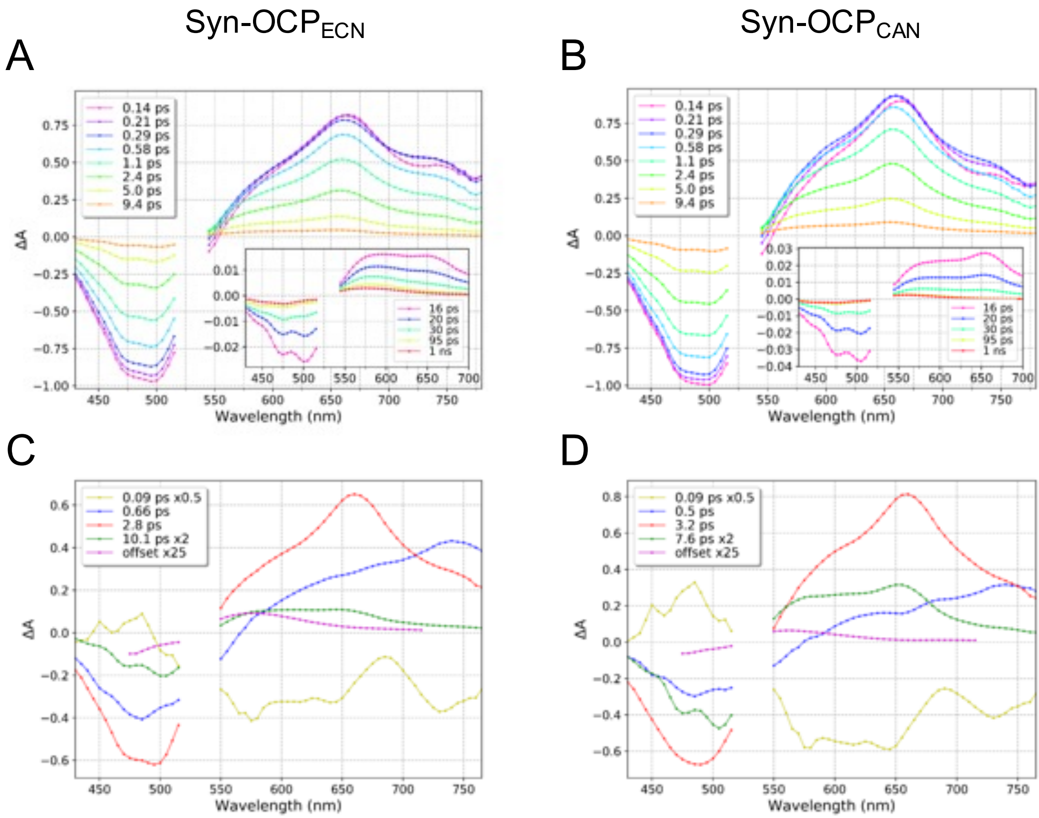

Syn-OCP$_{ECN}$
Syn-OCP$_{CAN}$
A
B
C
D
ΔA
Wavelength (nm)
0.14 ps
0.21 ps
0.29 ps
0.58 ps
1.1 ps
2.4 ps
5.0 ps
9.4 ps
16 ps
20 ps
30 ps
95 ps
1 ns
0.09 ps x0.5
0.66 ps
2.8 ps
10.1 ps x2
offset x25
0.09 ps x0.5
0.5 ps
3.2 ps
7.6 ps x2
offset x25

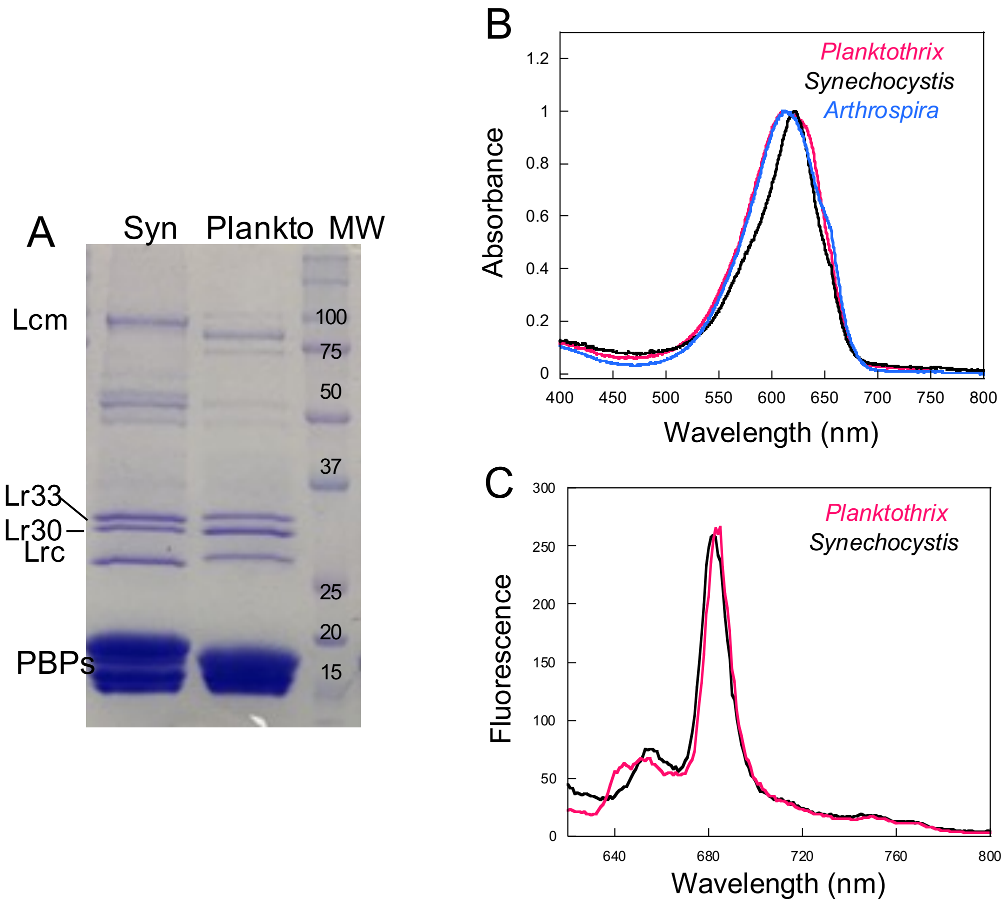
A
Syn
Plankto
MW
Lcm
100
75
50
37
Lr33
Lr30
Lrc
25
20
15
PBPs
B
Planktothrix
Synechocystis
Arthrospira
Absorbance
Wavelength (nm)
C
Planktothrix
Synechocystis
Fluorescence
Wavelength (nm)

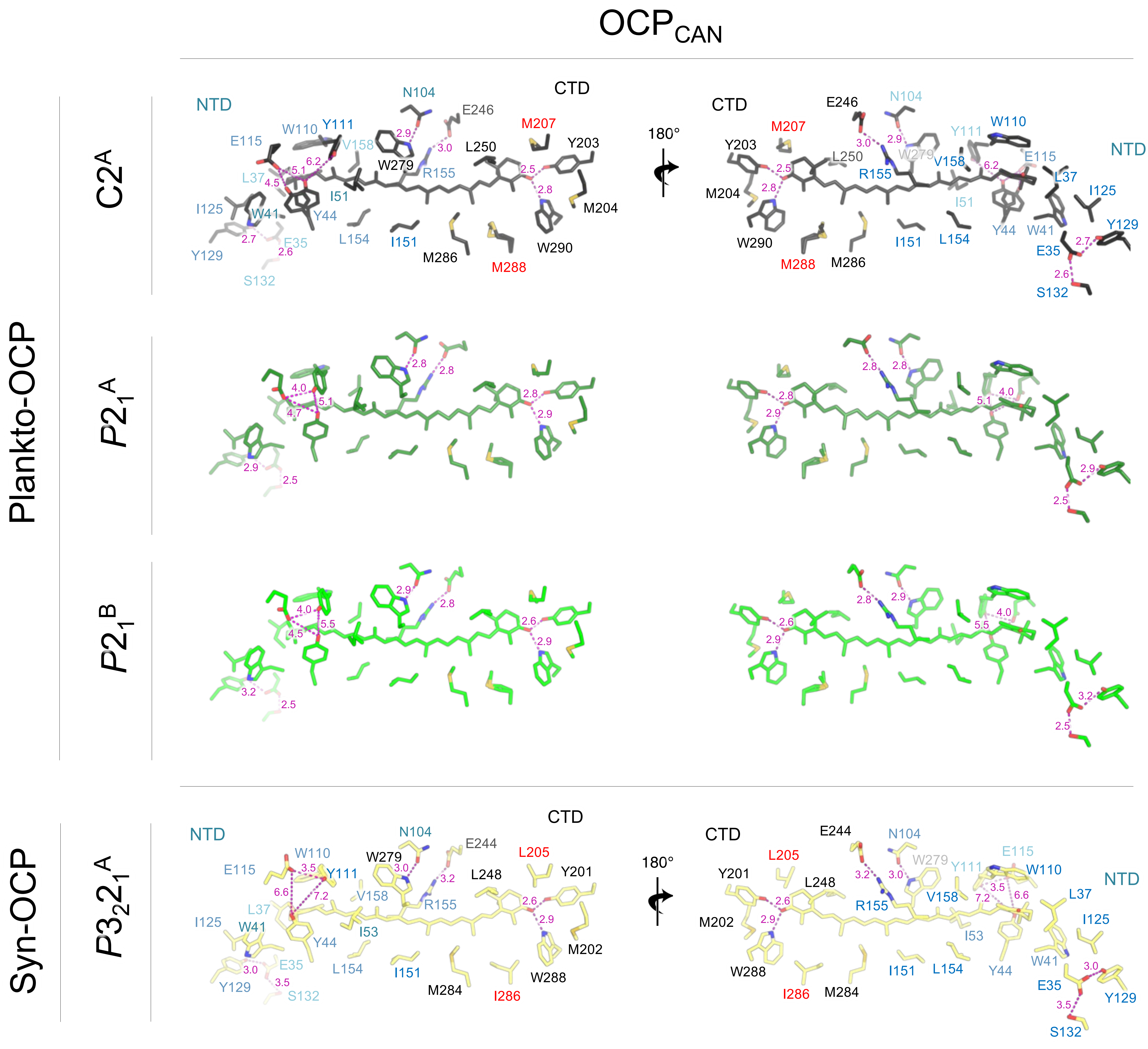
OCPCAN
Plankto-OCP
C2A
P21A
P21B
Syn-OCP
P3221A
NTD
CTD
180°
N104
E246
M207
Y203
L250
W110
Y111
E115
V158
L37
W279
R155
I51
I125
W41
Y44
L154
I151
M286
M288
W290
M204
Y129
E35
S132
E244
L205
Y201
L248
I53
M284
I286
W288
M202

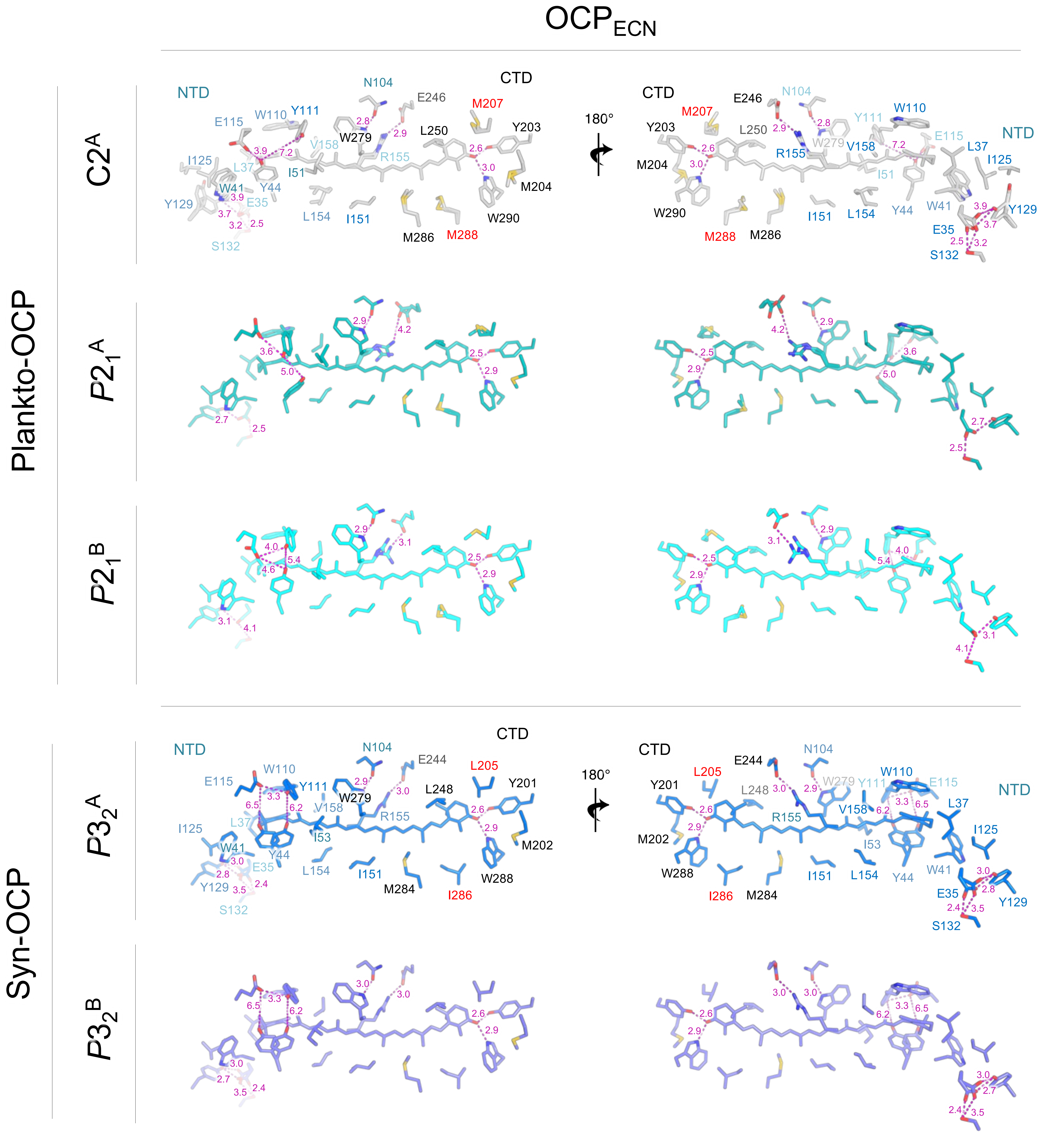

OCP$_{ECN}$
Plankto-OCP
Syn-OCP
C2$^A$
$P2_1^A$
$P2_1^B$
$P3_2^A$
$P3_2^B$
NTD
CTD
180°
N104
E246
M207
Y203
W110
Y111
E115
V158
W279
R155
L250
I125
L37
I51
Y44
W41
E35
Y129
S132
L154
I151
M286
M288
W290
M204
E244
L205
Y201
L248
I53
L154
I151
M284
I286
W288
M202

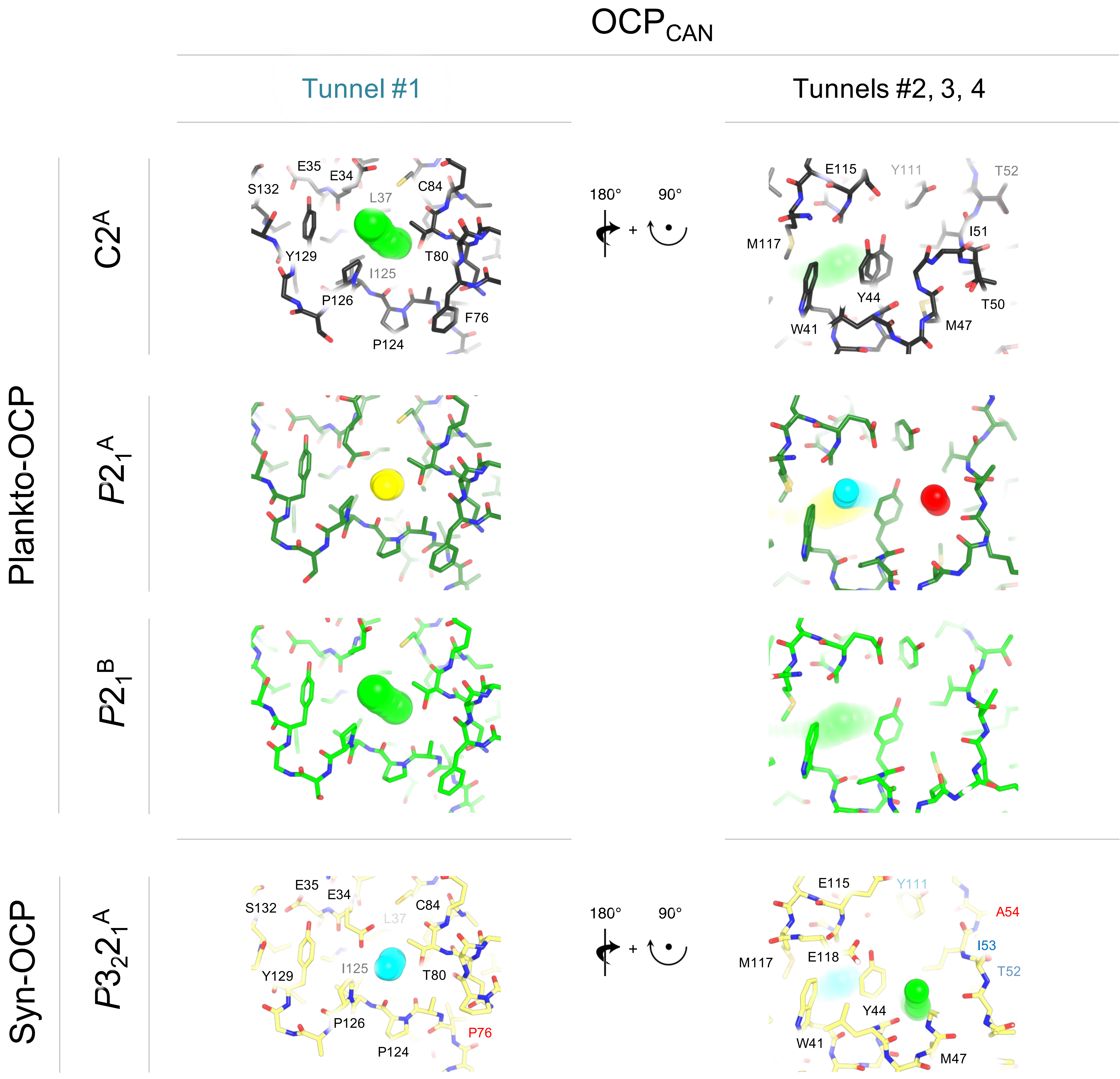
OCP_CAN
Tunnel #1
Tunnels #2, 3, 4
Plankto-OCP
C2^A
P2_1^A
P2_1^B
Syn-OCP
P3_2 2_1^A
180°
90°
+
E35
E34
S132
L37
C84
Y129
T80
I125
P126
F76
P124
P76
E115
Y111
T52
A54
I51
I53
M117
E118
Y44
T50
W41
M47

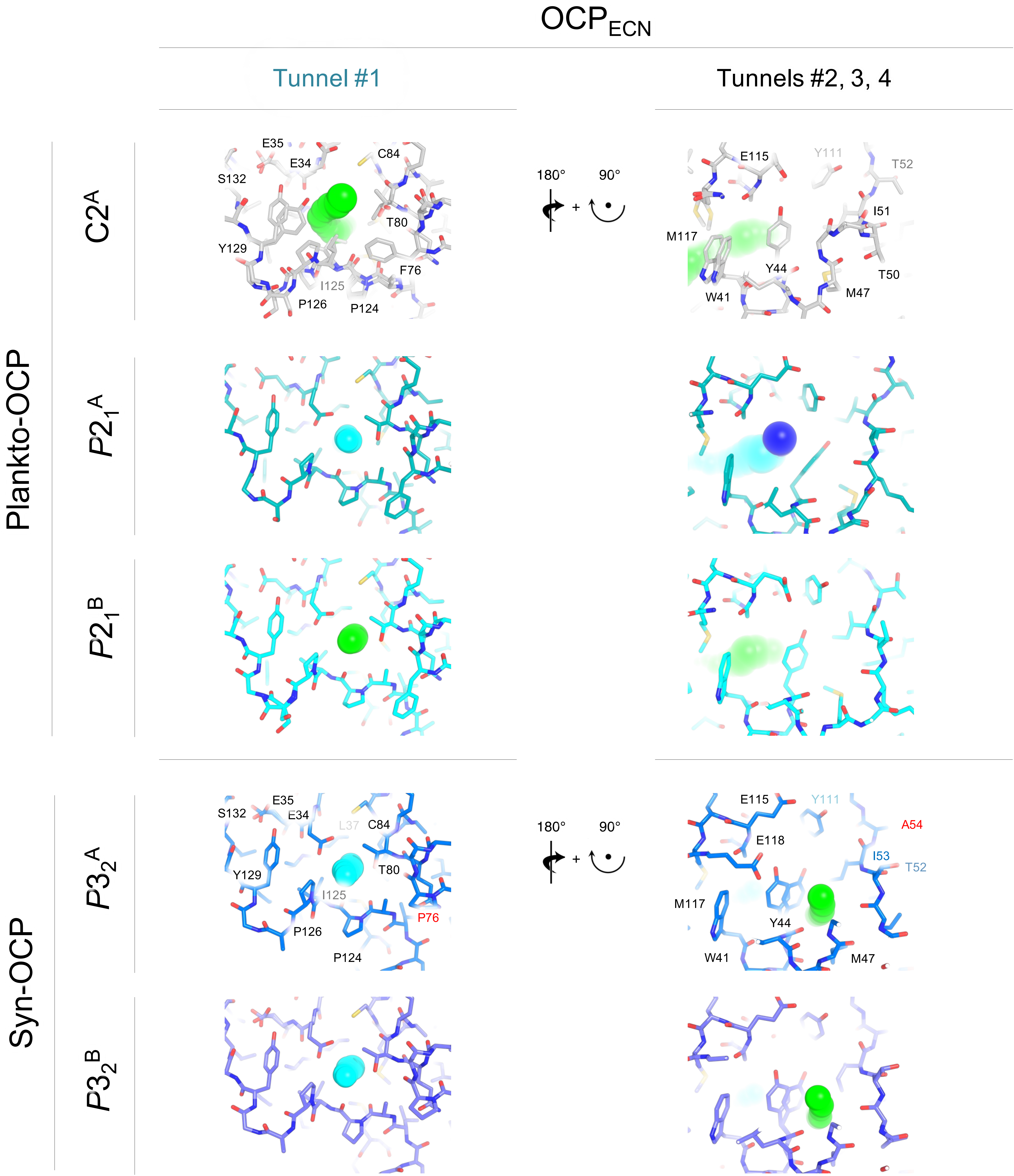

$OCP_{ECN}$
Tunnel #1
Tunnels #2, 3, 4
Plankto-OCP
Syn-OCP
$C2^{A}$
$P2_1^{A}$
$P2_1^{B}$
$P3_2^{A}$
$P3_2^{B}$
180°
90°
+
E35
E34
C84
S132
T80
Y129
F76
I125
P126
P124
E115
Y111
T52
I51
M117
Y44
T50
W41
M47
L37
P76
E118
A54
I53

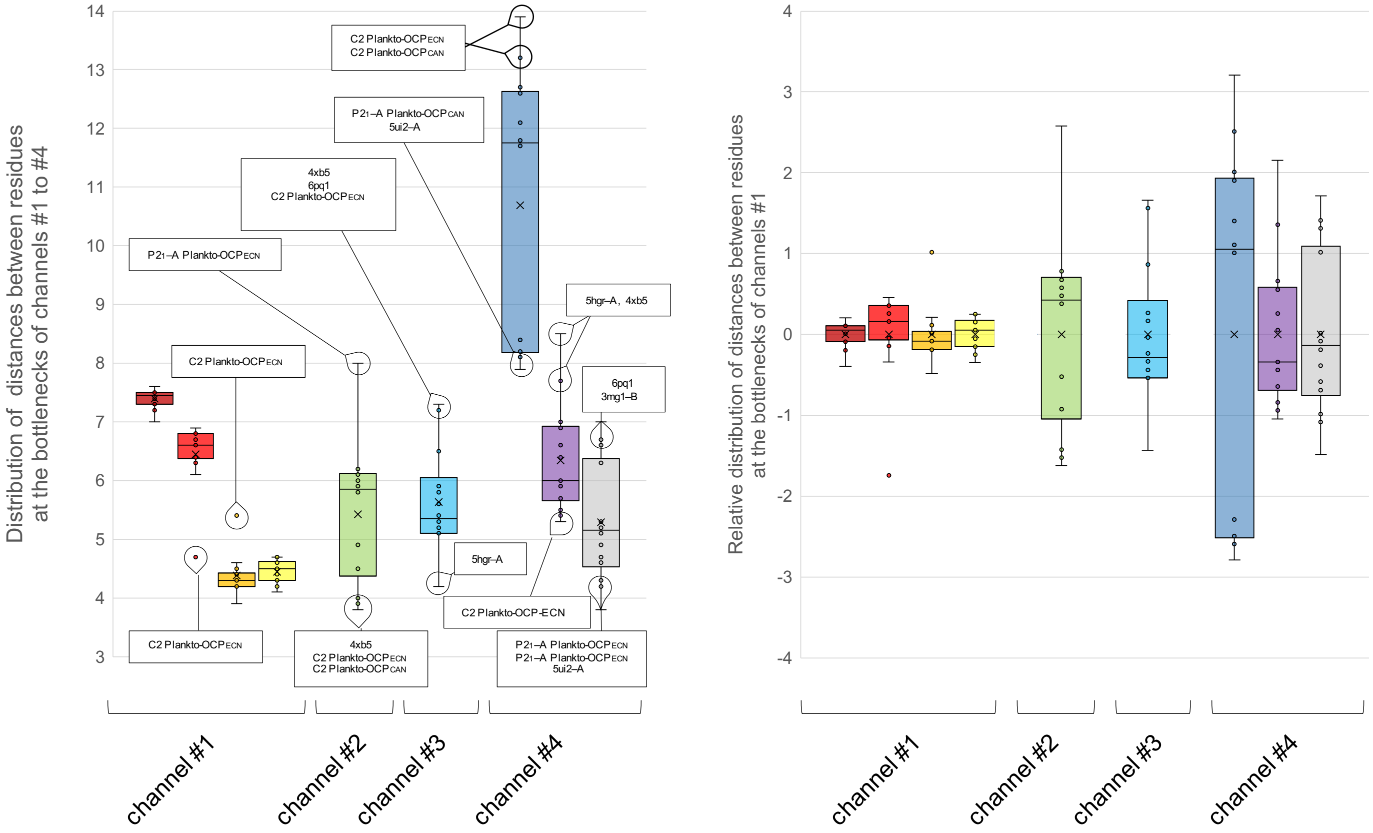
Distribution of distances between residues at the bottlenecks of channels #1 to #4
C2 Plankto-OCPECN
C2 Plankto-OCPCAN
P21–A Plankto-OCPCAN
5ui2–A
4xb5
6pq1
C2 Plankto-OCPECN
P21–A Plankto-OCPECN
5hgr–A, 4xb5
C2 Plankto-OCPECN
6pq1
3mg1–B
5hgr–A
C2 Plankto-OCP-ECN
C2 Plankto-OCPECN
4xb5
C2 Plankto-OCPECN
C2 Plankto-OCPCAN
P21–A Plankto-OCPECN
P21–A Plankto-OCPECN
5ui2–A
channel #1
channel #2
channel #3
channel #4
Relative distribution of distances between residues at the bottlenecks of channels #1
channel #1
channel #2
channel #3
channel #4

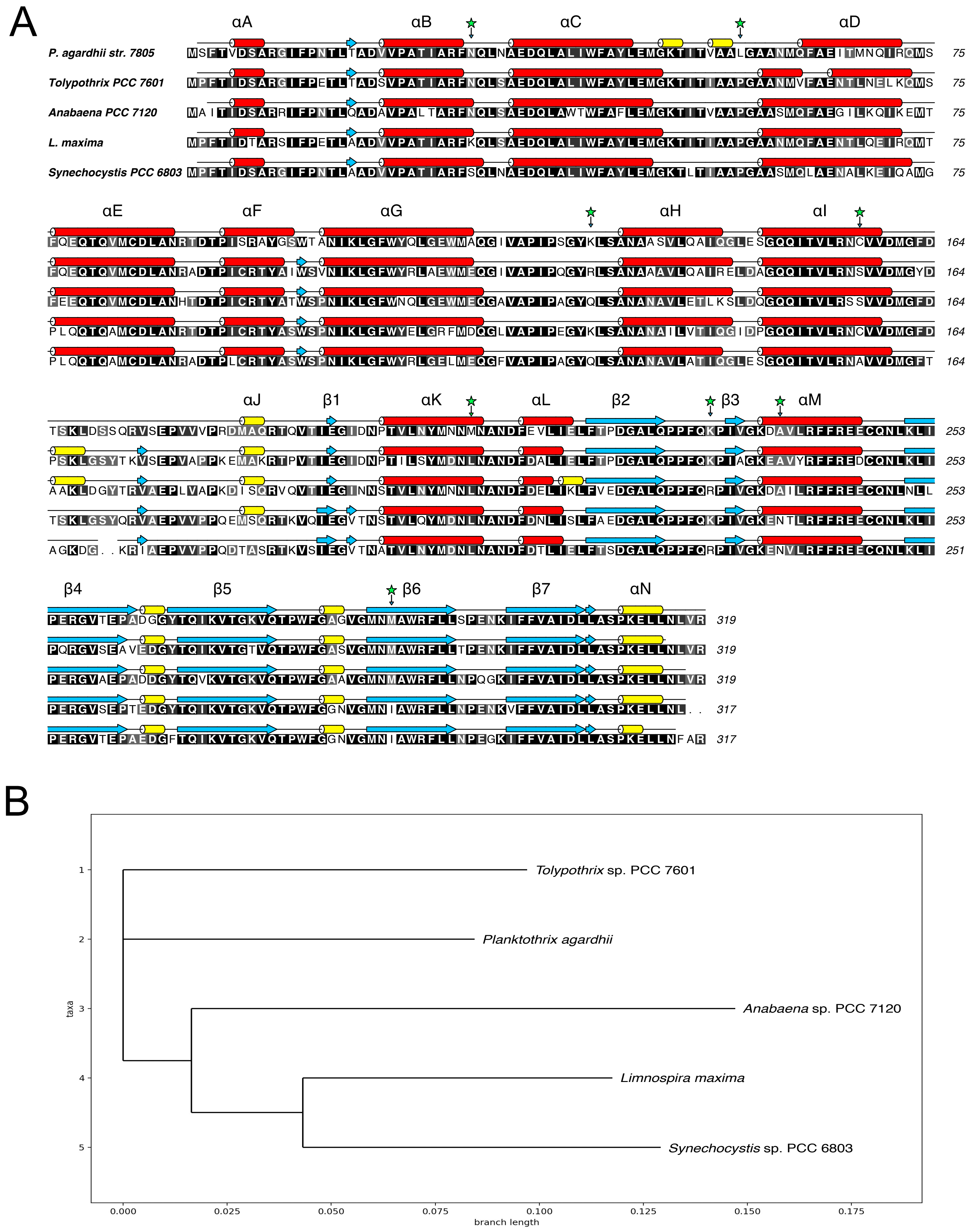
A
αA
αB
αC
αD
P. agardhii str. 7805
Tolypothrix PCC 7601
Anabaena PCC 7120
L. maxima
Synechocystis PCC 6803
75
αE
αF
αG
αH
αI
164
αJ
β1
αK
αL
β2
β3
αM
253
251
β4
β5
β6
β7
αN
319
317
B
Tolypothrix sp. PCC 7601
Planktothrix agardhii
Anabaena sp. PCC 7120
Limnospira maxima
Synechocystis sp. PCC 6803
taxa
1
2
3
4
5
0.000
0.025
0.050
0.075
0.100
0.125
0.150
0.175
branch length

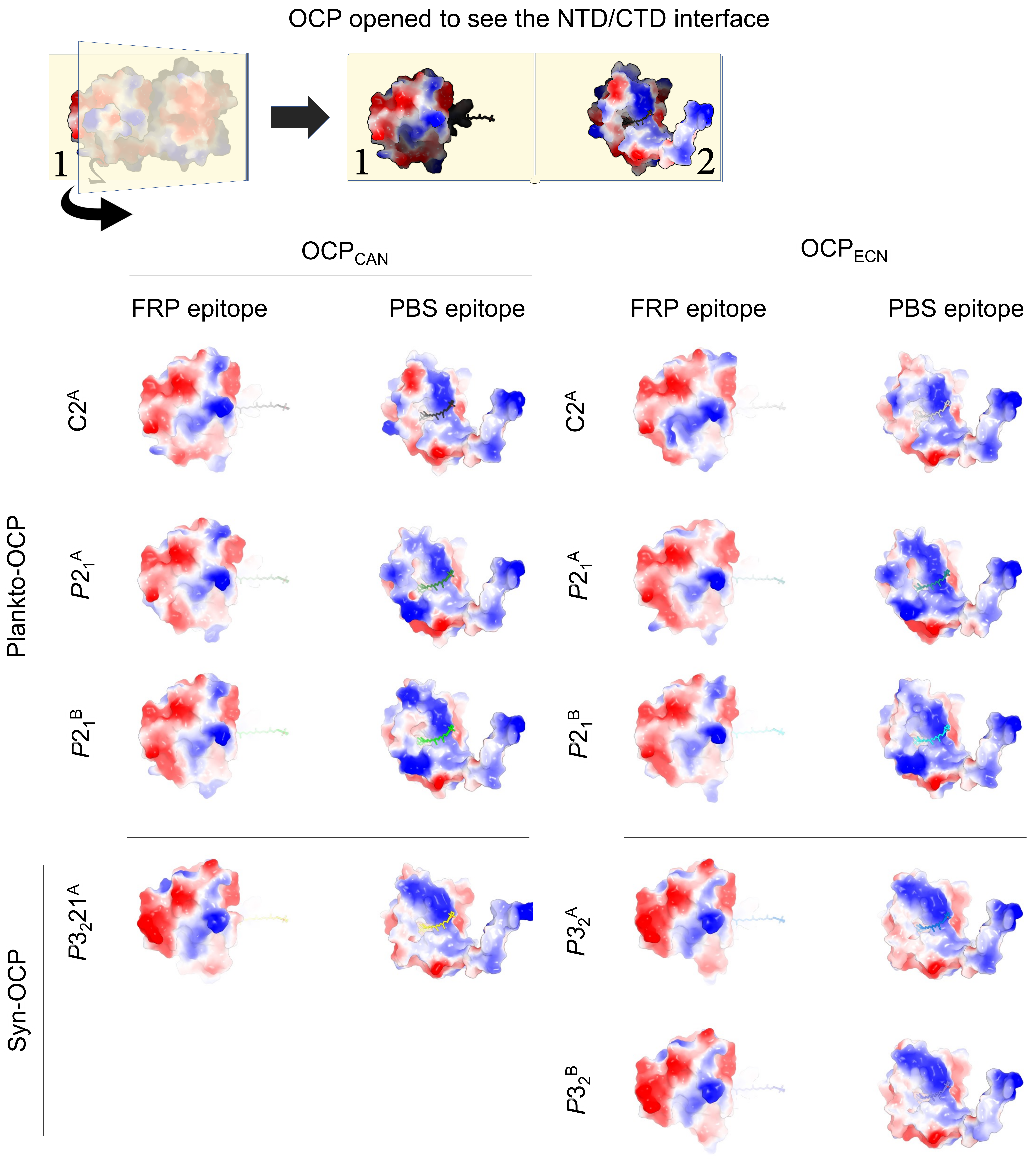

OCP opened to see the NTD/CTD interface
1
1
2
OCP$_{CAN}$
OCP$_{ECN}$
FRP epitope
PBS epitope
FRP epitope
PBS epitope
Plankto-OCP
C2$^A$
$P2_1^A$
$P2_1^B$
Syn-OCP
$P3_221^A$
$P3_2^A$
$P3_2^B$

| Examined structure | Contribution of secondary structure elements to the dimerisation interface | | | | | | | | | | | |
|---|---|---|---|---|---|---|---|---|---|---|---|---|
| | αA | αB | αE-αF loop | αH | β2-β3 loop | all | αA | αB | αE-αF loop | αH | β2-β3 loop | all |
| | buried surface area contribution to the dimerization interface (Å2) | | | | | | number of atoms involved in H-bonds or salt-bridges at the dimerization interface | | | | | |
| *$P2_1$* ECN | 289.83 | 430.49 | 81.25 | 196.64 | 86.29 | 1084.5 | 1 | 5 | 1 | 3 | 0 | 7 |
| *$P2_1$* CAN | 386.62 | 342.4 | 57.46 | 203.68 | 56.54 | 1046.7 | 3 | 5 | 1 | 3 | 0 | 8 |
| | fraction of the total BSA | | | | | | fraction of the total H-bond pool | | | | | |
| *$P2_1$* ECN | 0.27 | 0.40 | 0.07 | 0.18 | 0.08 | 1.00 | 0.10 | 0.50 | 0.10 | 0.30 | 0.00 | 1.00 |
| *$P2_1$* CAN | 0.37 | 0.33 | 0.05 | 0.19 | 0.05 | 1.00 | 0.25 | 0.42 | 0.08 | 0.25 | 0.00 | 1.00 |
| | % change in P21-ECN as compared to P21-CAN | | | | | | % change in H-bond pool of P21-ECN as compared to P21-CAN | | | | | |
| | -25.03 | 25.73 | 41.40 | -3.46 | 52.62 | 3.61 | -66.67 | 0.00 | 0.00 | 0.00 | n/a | -16.67 |
| | buried surface area contribution to the dimerization interface (Å2) | | | | | | number of atoms involved in H-bonds or salt-bridges at the dimerization interface | | | | | |
| C2 ECN | 306.98 | 429.03 | 78.38 | 174.43 | 33.88 | 1022.7 | 1 | 4 | 1 | 2 | 0 | 8 |
| C2 CAN | 336.83 | 423.86 | 73.82 | 193.21 | 33.18 | 1060.9 | 3 | 7 | 1 | 3 | 0 | 14 |
| | fraction of the total BSA | | | | | | fraction of the total H-bond pool | | | | | |
| C2 ECN | 0.30 | 0.42 | 0.08 | 0.17 | 0.03 | 1.00 | 0.13 | 0.50 | 0.13 | 0.25 | 0.00 | 1.00 |
| C2 CAN | 0.32 | 0.40 | 0.07 | 0.18 | 0.03 | 1.00 | 0.21 | 0.50 | 0.07 | 0.21 | 0.00 | 1.00 |
| | % change in C2-ECN as compared to C2-CAN | | | | | | % change in H-bond pool of C2-ECN as compared to P21-CAN | | | | | |
| | -8.86 | 1.22 | 6.18 | -9.72 | 2.11 | -3.60 | -66.67 | -42.86 | 0.00 | -33.33 | n/a | -42.86 |
| | % change in *$P2_1$*-ECN as compared to C2-ECN | | | | | | % change in H-bond pool of *$P2_1$*-ECN as compared to C2-ECN | | | | | |
| | -5.59 | 0.34 | 3.66 | 12.73 | 154.69 | 6.04 | 0.00 | 25.00 | 0.00 | 50.00 | n/a | 25.00 |
| | % change in *$P2_1$*-CAN as compared to C2-CAN | | | | | | % change in H-bond pool of *$P2_1$*-CAN as compared to C2-CAN | | | | | |
| | 14.78 | -19.22 | -22.16 | 5.42 | 70.40 | -1.34 | 0.00 | -28.57 | 0.00 | 0.00 | n/a | -14.29 |

| Examined structure | PDB id | Chain | Biological interface | Additional interface | Carotenoid BSA [A2] | Carotenoid SAA [A2] | Carotenoid BSA [%] | Carotenoid SAA [%] | Predicted radius of gyration | | rmsd to C2 OCP-CAN - monomer | rmsd to C2 OCP-CAN - dimer | rmsd to C2 OCP-CAN - NTD | rmsd to C2 OCP-CAN - all CTD | number of alternate conformations | Residues in alternate conformations |
|---|---|---|---|---|---|---|---|---|---|---|---|---|---|---|---|---|
| *P. aghardii* OCP-CAN, *P*2₁ → *P. aghardii* OCP-CAN, $P2_1$ | | A | 1001.60 | 1001.60 | 966.40 | 39.10 | 0.96 | 0.04 | 20.68 | 26.35 | 0.53 | 1.186 | 0.47 | 0.32 | 17 | D6, E34, T67, S95, M117, E146, S147, Q150, D183, P198, M207, R241, R244, E245, V258, K274, M288 |
| | | B | | | 965.10 | 41.50 | 0.96 | 0.04 | 20.53 | | 0.48 | | 0.42 | 0.27 | 36 | D6, N14, D35, Q36, M68, R72, Q73, S95, Q143, Q150, Q189, E193, D196, M207, R241, Q248, R256, V258, T259, Q268, K274, M288, K312 |
| *P. aghardii* OCP-ECN, $P2_1$ | | A | 1054.30 | 1054.60 | 945.70 | 25.20 | 0.97 | 0.03 | 20.63 | 26.26 | 0.69 | 1.087 | 0.54 | 0.27 | 17 | D6, M47, I51, E65, Q77, E115, M117, Q140, Q143, T152, V179, M207, R244, E246, E255, M288 |
| | | B | | | 940.00 | 36.00 | 0.96 | 0.04 | 20.45 | | 0.46 | | 0.45 | 0.25 | 17 | I25, E34, E46, M47, Q62, Q73, S95, I121, E176, R182, E193, D222, R241, R244, Q248, R256, M288 |
| *P. aghardii* OCP-CAN, C2 | | A | 1060.90 | 1650.30 | 969.40 | 37.80 | 0.96 | 0.04 | 20.52 | 26.12 | 0.00 | 0 | 0.00 | 0.00 | 17 | S2, S7, N14, N29, Y44, T50, T52, N60, Q62, M68, Q77, Q143, S147, D160, E176, K274, M288 |
| *P. aghardii* OCP-ECN, C2 | | A | 1022.70 | 1548.70 | 935.50 | 37.20 | 0.96 | 0.04 | 20.43 | 26 | 0.30 | 0.474 | 0.32 | 0.16 | 36 | R9, N14, N29, N32, L56, I66, M68, Q73, Q77, V82, M83, R89, M117, V122, A123, P124, I125, P126, S127, G128, Y129, K130, Q140, L154, R182, D183, Q186, M204, M207, K231, R241, K251, V258, M288, E297, K299 |
| *Synechocystis* PCC6803 OCP-CAN, $P3_22_1$ | 4xb5 | A | 1089.50 | n/a | 956.10 | 37.30 | 0.96 | 0.04 | 20.45 | 26.04 | 0.44 | 0.885 | 0.62 | 0.26 | 1 | E118 |
| *Synechocystis* PCC6803 OCP-ECN, $P3_2$ | 3mg1 | A | 1095.70 | n/a | 943.00 | 40.40 | 0.96 | 0.04 | 20.36 | 25.87 | 0.39 | 0.829 | 0.58 | 0.28 | 11 | I25, D35, Q36, Y44, E46, M47, Q62, E70, M83, Q150, V273 |
| | | B | | | 940.60 | 40.30 | 0.96 | 0.04 | 20.36 | | 0.40 | | 0.56 | 0.25 | 10 | I25, D35, Y44, E46, S60, M74, Q150, I172, Q228, I251 |
| *Limnospira maxima* OCP-ECN, C2 | 5ui2 | A | 1186.10 | n/a | 965.50 | 34.50 | 0.97 | 0.03 | 20.58 | 26.43 | 0.45 | 1.546 | 0.57 | 0.28 | 0 | |
| | | B | | | 968.90 | 31.10 | 0.97 | 0.03 | 20.61 | | 0.50 | | 0.63 | 0.28 | 0 | |
| *Anabaena* PCC7120 OCP-CAN, *P*1 | 5hgr | A | 1089.50 | n/a | 955.00 | 41.90 | 0.96 | 0.04 | 20.48 | 26.68 | 0.46 | 2.287 | 0.58 | 0.28 | 0 | |
| | | B | | | 947.20 | 43.90 | 0.96 | 0.04 | 20.54 | | 0.55 | | 0.60 | 0.29 | 0 | |
| *Tolipothrix* OCP-CAN, $P3_22_1$ | 6pq1 | A | 856.90 | n/a | 964.30 | 24.70 | 0.98 | 0.02 | 20.69 | 26.77 | 0.52 | 1.983 | 0.52 | 0.24 | 2 | N156, S157 |
| Mean | | | 1050.80 | 1034.88 | 954.48 | 36.49 | 0.96 | 0.04 | 20.52 | 26.28 | 0.44 | 0.49 | 0.49 | 0.24 | 10.36 | |
| Standard deviation | | | 89.61 | 333.20 | 12.19 | 5.87 | 0.01 | 0.01 | 0.11 | 0.31 | 0.16 | 0.16 | 0.16 | 0.08 | 13.91 | |

| Examined structure | Contribution of secondary structure elements to the dimerisation interface | | | | | | | |
|---|---|---|---|---|---|---|---|---|
| | αC-αD | αD | αE | αG + αG-αH loop | linker | αM-β4 loop | β5-β6 loop | all |
| | buried surface area contribution to the dimerization interface (Å2) | | | | | | | |
| *$P2_1$* ECN | 374.57 | 108.54 | 51.72 | 315.29 | 104.28 | 43.07 | 49.18 | 1046.65 |
| *$P2_1$* CAN | 375.32 | 122.45 | 51.76 | 297.18 | 85.52 | 35.21 | 42.07 | 1009.51 |
| | fraction of the total BSA | | | | | | | |
| *$P2_1$* ECN | 0.36 | 0.10 | 0.05 | 0.30 | 0.10 | 0.04 | 0.05 | 1.00 |
| *$P2_1$* CAN | 0.37 | 0.12 | 0.05 | 0.29 | 0.08 | 0.03 | 0.04 | 1.00 |
| | % change in P21-ECN as compared to P21-CAN | | | | | | | |
| | -0.20 | -11.36 | -0.08 | 6.09 | 21.94 | 22.32 | 16.90 | 3.68 |
| | buried surface area contribution to the dimerization interface (Å2) | | | | | | | |
| C2 ECN | 303.57 | 335.79 | 152.89 | 331.49 | 323.09 | 68.83 | 31.77 | 1547.43 |
| C2 CAN | 349.13 | 343.01 | 171.52 | 337.92 | 326.91 | 88.24 | 27.83 | 1644.56 |
| | fraction of the total BSA | | | | | | | |
| C2 ECN | 0.20 | 0.22 | 0.10 | 0.21 | 0.21 | 0.04 | 0.02 | 1.00 |
| C2 CAN | 0.21 | 0.21 | 0.10 | 0.21 | 0.20 | 0.05 | 0.02 | 1.00 |
| | % change in P21-ECN as compared to *$P2_1$*-CAN | | | | | | | |
| | -13.05 | -2.10 | -10.86 | -1.90 | -1.17 | -22.00 | 14.16 | -5.91 |
| | % change in *$P2_1$*-ECN as compared to C2-ECN | | | | | | | |
| | 23.39 | -67.68 | -66.17 | -4.89 | -67.72 | -37.43 | 54.80 | -32.36 |
| | % change in *$P2_1$*-CAN as compared to C2-CAN | | | | | | | |
| | 7.50 | -64.30 | -69.82 | -12.06 | -73.84 | -60.10 | 51.17 | -38.62 |

| Structure | PDB id | Chain | Status of channel #1 | L37(CD2) -- M117(SD) | M83(SD) -- I125(CD1) | L37(CB) -- I125(CD1) | M83(SD) -- M117(SD) | Status of channel #2 | W41(CZ3) -- Y44(CE2) | Status of channel #3 | Y44(CE1) -- Y111(CD1) * | Status of channel #4 | K49(CA) -- C147(CA) * | Met47(SD) -- Y44 (CE1) | Met47(SD) -- F280(CZ) * |
|---|---|---|---|---|---|---|---|---|---|---|---|---|---|---|---|
| P. aghardii OCP-CAN, $P2_1$ | | A | open | 7.3 | 6.8 | 4.3 | 4.5 | open | 6.1 | open | 5.1 | close | 8.1 | 6.6 | 5.3 |
| | | B | open | 7.5 | 6.8 | 4.3 | 4.7 | open | 5.8 | close | 5.2 | close | 8.4 | 5.3 | 4.6 |
| P. aghardii OCP-ECN, $P2_1$ | | A | open | 7.5 | 6.7 | 4.3 | 4.7 | open | 8 | open | 5.1 | close | 8.4 | 5.5 | 4.3 |
| | | B | open | 7.3 | 6.3 | 4.4 | 4.5 | open | 5.9 | close | 5.2 | close | 8.2 | 5.4 | 4.2 |
| P. aghardii OCP-CAN, C2 | | A | open | 7.5 | 6.1 | 3.9 | 4.5 | close | 3.9 | close | 5.6 | open | 13.2 | 6 | 5.1 |
| P. aghardii OCP-ECN, C2 | | A | open | 7.5 | 4.7 | 5.4 | 4.7 | close | 3.8 | close | 5.1 | close | 13.9 | 5.9 | 5.3 |
| S. PCC6803 OCP-CAN, $P3_22_1$ | 4xb5 | A | open | 7.2 | 6.6 | 4.2 | 4.3 | close | 4 | open | 7.2 | open | 12.7 | 8.5 | 6.6 |
| S. PCC6803 OCP-ECN, $P3_2$ | 3mg1 | A | open | 7.5 | 6.4 | 4.2 | 4.3 | close | 6.1 | open | 5.8 | open | 12.6 | 6.4 | 5.2 |
| | | B | open | 7.5 | 6.4 | 4.2 | 4.2 | close | 6 | open | 5.9 | open | 12.6 | 7 | 6.7 |
| L. maxima OCP-ECN, C2 | 5ui2 | A | open | 7.6 | 6.4 | 4.6 | 4.4 | open | 6.2 | open | 5.4 | close | 8.1 | 6 | 3.8 |
| | | B | open | 7.3 | 6.6 | 4.3 | 4.1 | open | 6.2 | open | 5.3 | close | 7.9 | 5.9 | 6.3 |
| A. PCC7120 OCP-CAN, $P1$ | 5hgr | A | open | 7.4 | 6.8 | 4.4 | 4.6 | close | 4.9 | open | 4.2 | close | 11.7 | 5.7 | 4.9 |
| | | B | open | 7.4 | 6.9 | 4.5 | 4.5 | close | 4.5 | close | 6.5 | close | 12.1 | 7.7 | 4.7 |
| Toplipothrix OCP-CAN | 6pq1 | A | open | 7 | 6.7 | 4.4 | 4.3 | open | 4.5 | open | 7.3 | close | 11.8 | 6.9 | 7 |
| Mean | | | | 7.39 | 6.44 | 4.39 | 4.45 | | 5.42 | | 5.64 | | 10.69 | 6.34 | 5.29 |
| Standard deviation | | | | 0.16 | 0.55 | 0.33 | 0.19 | | 1.19 | | 0.86 | | 2.32 | 0.92 | 1.00 |
